# CD4^+^ T cells drive corneal nerve damage but not epitheliopathy in an acute aqueous-deficient dry eye model

**DOI:** 10.1101/2024.03.22.586336

**Authors:** Alexia Vereertbrugghen, Manuela Pizzano, Agostina Cernutto, Florencia Sabbione, Irene A Keitelman, Douglas Vera Aguilar, Ariel Podhorzer, Federico Fuentes, Celia Corral-Vázquez, Mauricio Guzmán, Mirta N Giordano, Analía Trevani, Cintia S de Paiva, Jeremías G Galletti

**Author notes:** **Corresponding Author:** Jeremías G Galletti. **Author Contributions:** AV, MP, MNG, AT, CDP, and JGG contributed to the conception and design of the work; AV, MP, AC, FS, IAK, DVA, AP, FF, CCV, MNG, and JGG contributed to the acquisition and analysis of data; AV, MP, MNG, AT, CDP, and JGG contributed to the interpretation of data; AV, MG, AT, MNG, CDP, and JGG drafted and/or substantively revised the manuscript. All authors read and approved the final manuscript. **Competing Interest Statement:** All authors declare that there is no conflict of interest.

## Abstract

Dry eye disease (DED) is characterized by a dysfunctional tear film in which the cornea epithelium and its abundant nerves are affected by ocular desiccation and inflammation. Although adaptive immunity and specifically CD4^+^ T cells play a role in DED pathogenesis, the exact contribution of these cells to corneal epithelial and neural damage remains undetermined. To address this, we explored the progression of a surgical DED model in wild-type (WT) and T cell-deficient mice. We observed that adaptive immune-deficient mice developed all aspects of DED comparably to WT mice except for the absence of functional and morphological corneal nerve changes, nerve damage-associated transcriptomic signature in the trigeminal ganglia, and sustained tear cytokine levels. Adoptive transfer of CD4^+^ T cells from WT DED mice to T cell-deficient mice reproduced corneal nerve damage but not epitheliopathy. Conversely, T cell-deficient mice reconstituted solely with naive CD4^+^ T cells developed corneal nerve impairment and epitheliopathy upon DED induction, thus replicating the WT DED phenotype. Collectively, our data show that while corneal neuropathy is driven by CD4^+^ T cells in DED, corneal epithelia damage develops independently of the adaptive immune response. These findings have implications for T cell-targeting therapies currently in use for DED.

**Graphical abstract:** 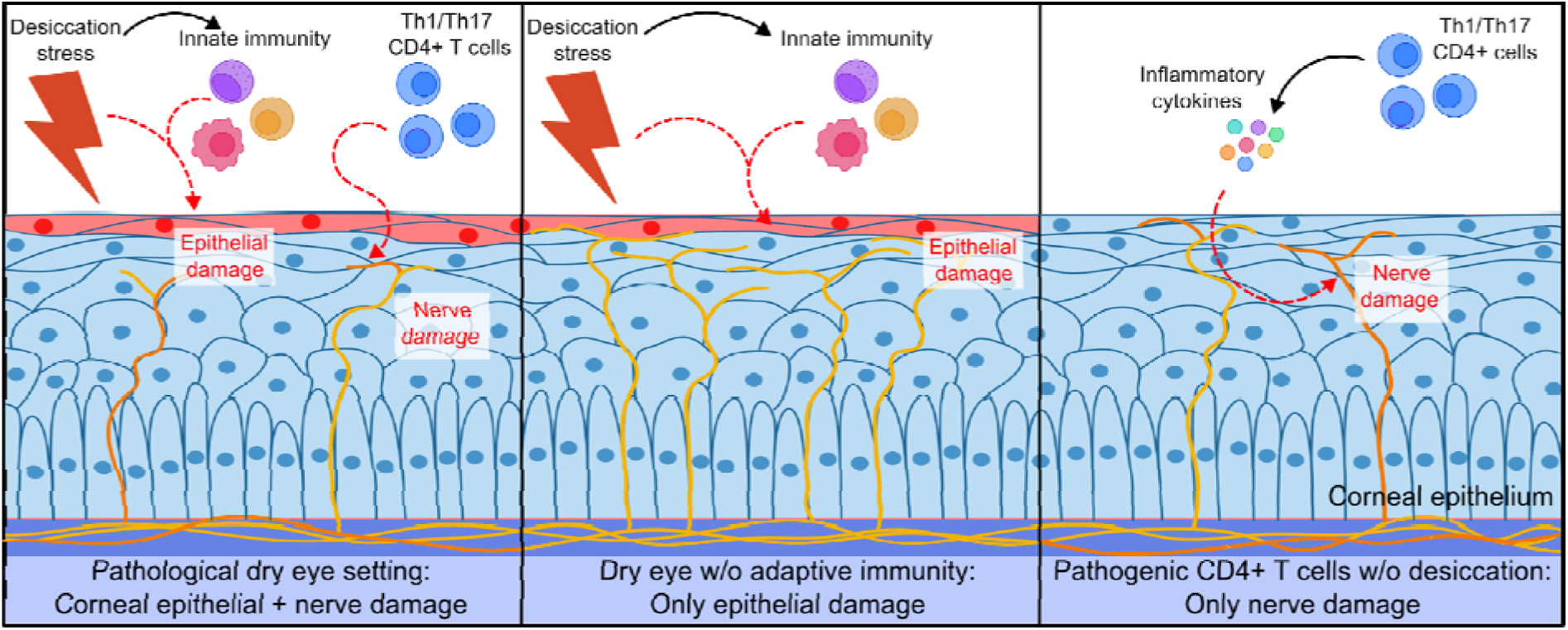

**Significance Statement:** Dry eye is a frequent ocular disorder in which damage to the corneal epithelium and nerves is triggered by inadequate lubrication. The local CD4^+^ T cell-predominant immune response aggravates ocular surface impairment but the exact contribution of these cells to corneal epithelial and neural disease remains undetermined. Using adoptive transfer of T cells into T cell-deficient mice, trigeminal transcriptomics, and tear cytokine analysis, we delineate the pathogenic role of CD4^+^ T cells, revealing that they drive corneal nerve damage but are dispensable for epithelial disease to develop in response to desiccation. CD4^+^ T cells promote corneal neuropathy possibly by releasing proinflammatory cytokines onto the ocular surface. These findings have implications for T cell-targeting therapies currently used for dry eye.

## Introduction

The ocular surface is the mucosal microenvironment that encircles the cornea and keeps it moist and protected for proper sight under homeostatic conditions(1, 2). Conversely, dry eye disease (DED) is an increasingly prevalent ocular surface disorder characterized by a dysfunctional tear film, visual disturbance, local inflammation and tissue damage, neurosensory abnormalities, and symptoms ranging from ocular discomfort to overt pain(3). DED has heterogeneous clinical presentations. While some patients have overt ocular surface epitheliopathy accompanied by discomfort or pain, others present mainly with symptoms derived from neurosensory alterations and little or no ocular epithelial damage(4, 5). Ocular surface inflammation plays a pivotal role in perpetuating the disease by favoring tissue damage, which in turn elicits more inflammation and creates a vicious cycle(6–8).

CD4^+^ T cells orchestrate the local adaptive immune response and drive DED-associated ocular inflammation(7, 8). DED patients have more conjunctival CD4^+^ T cells(9) and two FDA-approved drugs (cyclosporine and lifitegrast) for topical anti-inflammatory therapy selectively block T cell activation locally(7, 10). In animal models, the adoptive transfer of CD4^+^ T cells from DED mice reproduces conjunctival goblet cell loss and lacrimal gland infiltration (11, 12) whereas depletion of ocular surface antigen-presenting cells prevents local CD4^+^ T cell activation and goblet cell loss(13). DED increases T-helper (Th)1 and Th17 CD4^+^ T cells in the eye-draining lymph nodes of mice(14, 15), and both interferon (IFN)-γ and interleukin (IL)-17 are increased in the tears and conjunctiva of DED patients(16–19). IFN-γ, a hallmark type 1 cytokine, promotes squamous metaplasia and apoptosis in ocular surface epithelial cells(12, 20–24). IL-17, a hallmark type 3 cytokine, induces matrix metalloproteinase secretion and intercellular junction degradation, causing corneal barrier disruption(19, 25). However, CD4^+^ T cells are not the only source of IFN-γ and IL-17 in the ocular surface because CD8^+^ T cells, γδ T cells, NK cells, and other innate lymphoid cells also secrete these cytokines(26–31). Thus, while type 1 and 3 immunity contributes to DED-associated corneal epitheliopathy, it may be mediated by CD4^+^ T cells and/or innate immune cells(29).

By contrast, the pathophysiology of neurosensory abnormalities in DED is unclear(32, 33). The intraepithelial endings of the corneal nerves are almost in direct contact with the tear film and are impacted the most in DED(32). Since these nerves rely entirely on corneal epithelial cells for support(34) and become affected early in the course of the disease and coinciding with epithelial damage(35), it is assumed that corneal neuropathy in DED is secondary to corneal epitheliopathy and caused by the same mechanisms. In mice, tear hyperosmolarity elicits corneal nerve changes without corneal barrier disruption(36) and corneal neuropathy may develop independently of corneal epitheliopathy if a type 1 immune response is present in the ocular surface(37). However, since both models lack ocular desiccation, the defining disease feature that initiates corneal pathology(3, 4), their conclusions may not apply to DED because additional mechanisms could be at play. The disease setting determines the pathogenic impact (or lack thereof) of the local immune response on the corneal nerves: herpetic keratitis and ocular graft-versus-host-disease but not ocular allergy exhibit corneal neuropathy despite all three being immune-driven ocular surface disorders(38). In the case of DED, this issue has not been addressed. We hypothesized that the local CD4^+^ T cell-coordinated adaptive immune response in DED induces corneal neuropathy independently from corneal epitheliopathy. Therefore, we explored the relative contribution of the adaptive immune response, and more specifically of CD4^+^ T cells, to corneal epithelial and nerve damage in a murine model of DED.

## Results

### 1. Impact of adaptive immunity deficiency on DED phenotype development

CD4^+^ T cells are pathogenic in DED because they enhance desiccation-induced ocular surface inflammation(7, 9–12, 14, 15), but whether a CD4^+^ T cell-coordinated adaptive immune response is required for either corneal epitheliopathy or neuropathy to develop in this disorder remains undetermined. Therefore, we compared DED progression in immunocompetent wild-type (WT) and adaptive immunity-deficient *Rag1*KO (lacking CD4^+^ and CD8^+^ T, NKT, and B cells) mice using a surgical model (Figure 1A) that involves the bilateral excision of the extraorbital lacrimal gland(39). The resulting desiccation of the ocular surface triggers a pathogenic immune response, thus mimicking aqueous-deficient DED(40, 41). Tear deficiency 5 days after surgery was comparable in both strains (Figure 1B) and baseline tear production did not differ between strains nor sexes (sex factor, <2% total variation, p=0.50). Also, WT and *Rag1*KO mice exhibited similar external eye phenotype 10 days after DED induction (Figure 1C). Next, we quantified conjunctival goblet cells because their loss is a clinically validated finding in DED that correlates with disease severity(42, 43). Total conjunctival goblet cell area did not differ between control mice while DED induction led to a comparable loss of goblet cells in both strains (Figures 1D-E). Altogether this data shows that both mouse strains develop comparable DED severity in terms of non-corneal phenotype. Thus, the *Rag1*KO DED model is valid for assessing the impact of the lack of adaptive immunity on the corneal aspects of the disease.

**Figure 1.**
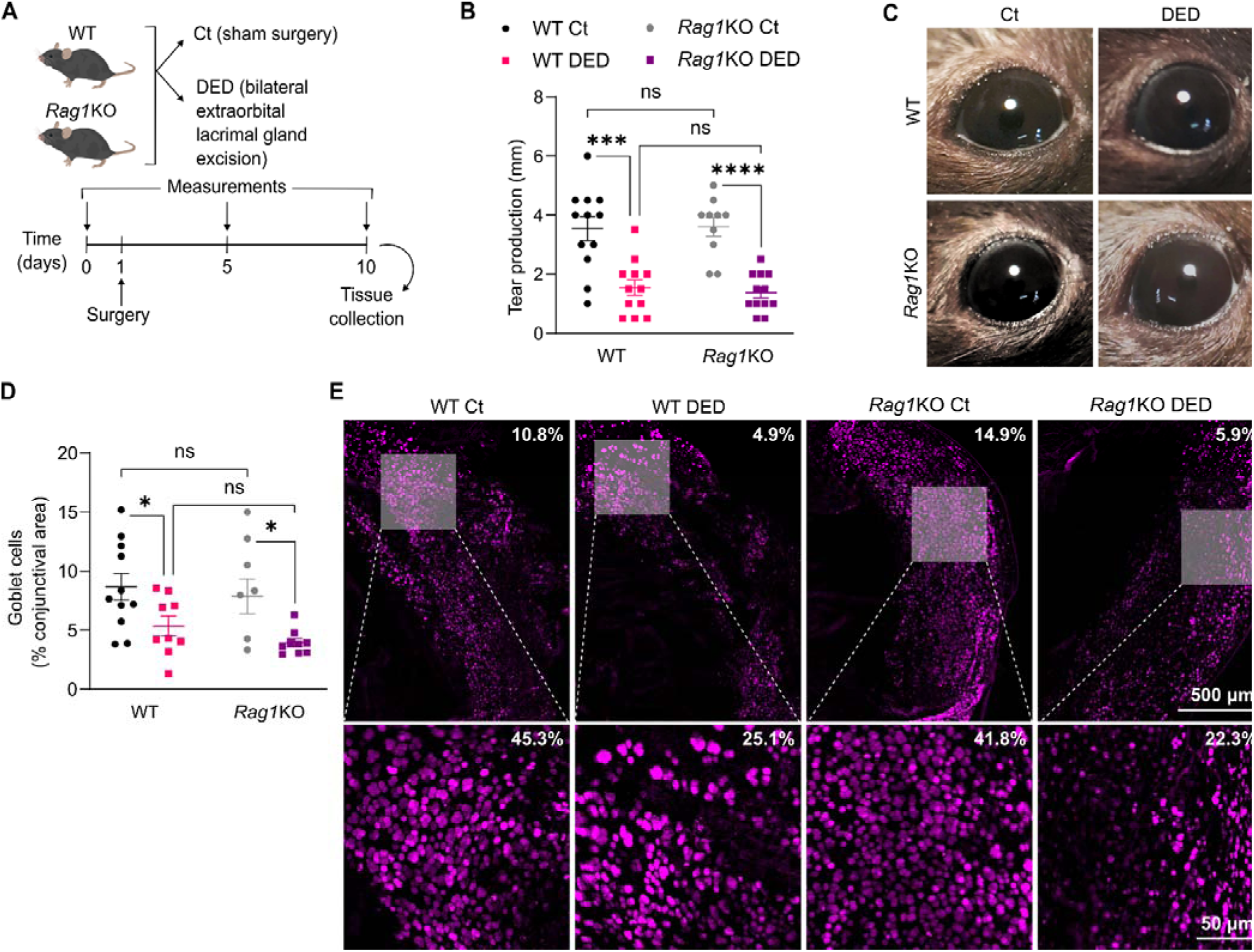
Impact of adaptive immune deficiency on dry eye phenotype development. **A)** Dry eye disease (DED) was surgically induced in wild-type (WT) or recombination-activating gene 1-knockout (*Rag1*KO) mice of both sexes through bilateral excision of the extraorbital lacrimal gland. Sham-operated animals were included as controls (Ct). **B)** Tear production on day 5 as measured by phenol red-paper wetting length. **C)** External eye appearance 10 days after surgery. **D)** Total conjunctival area occupied by goblet cells 10 days after DED induction. **E)** Representative micrographs of wheat germ agglutinin-stained goblet cells in conjunctival whole-mounts used for quantification. Low-magnification micrographs (top) of whole-mounted conjunctival strips used for measurements shown in D, with their corresponding high-magnification insets of the goblet cell-rich area (bottom) provided only as a reference. The actual goblet cell-occupied area is shown in the top right corner of each image. All experiments were performed twice or more with 6 mice/group/experiment. To compare means, two-way ANOVA was used for C and D (strain and treatment) with Sidak’s post hoc test. * indicates p<0.05, *** indicates p<0.001, **** indicates p<0.0001, and ns indicates not significant.

### 2. Adaptive immune deficiency does not impede DED-associated corneal epitheliopathy development

Corneal epitheliopathy is a hallmark DED sign, characterized by epithelial barrier dysfunction and accelerated cell turnover(4). Type 1 and 3 immunity promotes corneal epithelial damage(12, 19, 20, 23, 25), though whether the adaptive component of the immune response is required for corneal epitheliopathy to develop in DED is unclear. We first probed corneal barrier function through dye uptake(4, 44). In DED, increased epithelial cell shedding and matrix metalloproteinase activity disrupt tight junctions, thus allowing the intercellular diffusion of fluorescein-tagged dextran(44). Remarkably, DED mice from both strains developed a comparable time-dependent increase in corneal dye uptake(4, 44)(Figures 2A-B). Sub-analysis (Figure S1A, 3-way ANOVA with factors strain, sex, and time) revealed that time had the largest effect on corneal epitheliopathy development (60.7% of total variation, p<0.0001), female mice had worse epithelial disease than males (sex factor accounted for 9.1% of total variation, p<0.0001), and strain had no significant overall effect (1.4% of total variation, p=0.08). Female WT DED mice had more corneal dye uptake on day 5 (p=0.01) than female *Rag1*KO DED mice but this difference disappeared by day 10 (p=0.59), suggesting a faster tempo (Figure S1A). Next, we quantified the number of proliferating corneal epithelial cells as an indicator of the increased turnover rate in DED(45). Both WT and *Rag1*KO DED mice had a comparably increased number of proliferating (Ki67^+^) corneal epithelial cells (Figures 2C and S1B). Proliferating cells were restricted to the basal epithelial layer in control mice while they extended to the suprabasal layer in DED mice of both strains (Figure S1C), a pattern consistent with DED-induced corneal epitheliopathy(45). These results suggest that the absence of an adaptive immune response does not condition the development of corneal epitheliopathy in this DED model.

**Figure 2.**
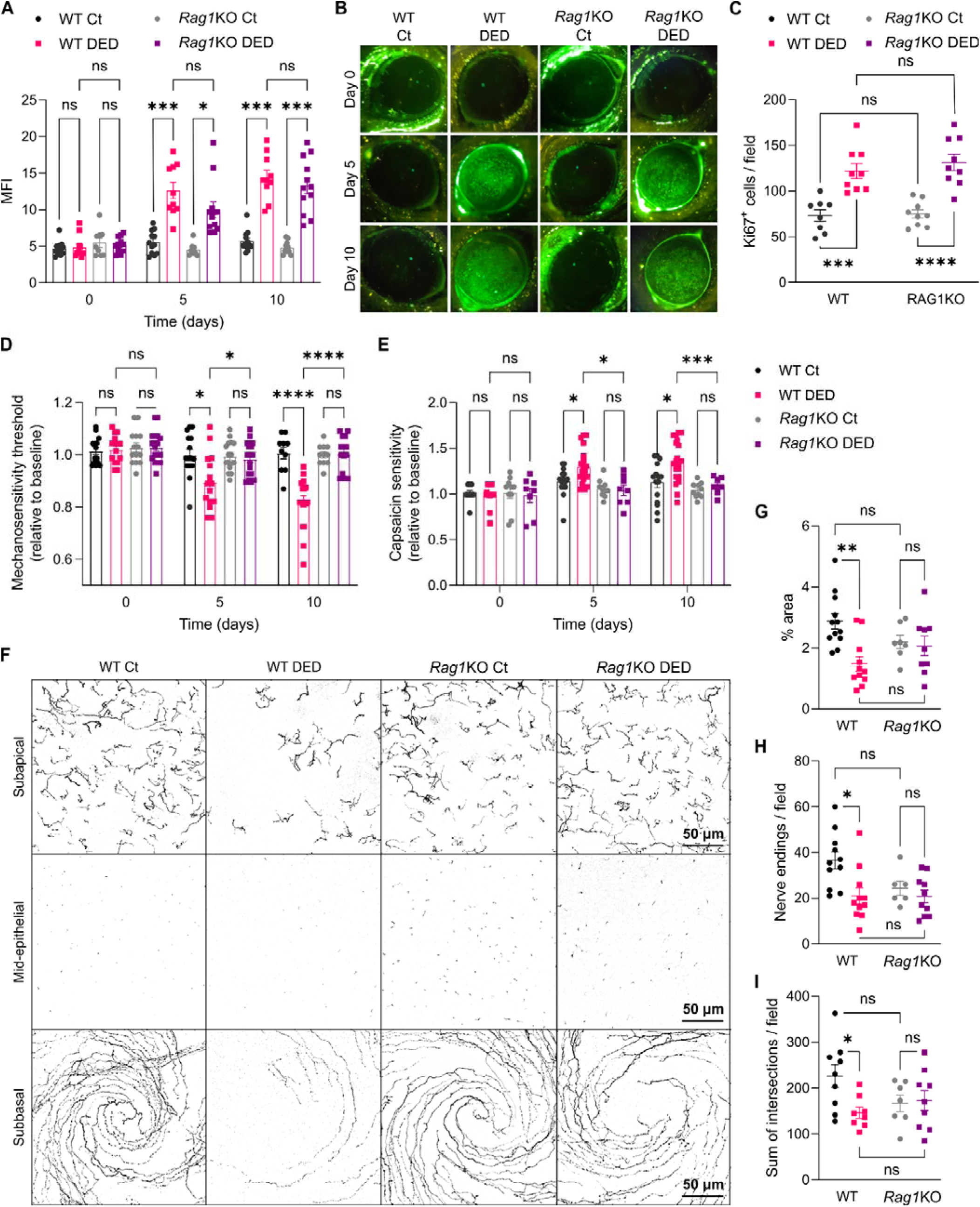
Lack of adaptive immunity does not hinder corneal epitheliopathy but prevents corneal neuropathy development in dry eye. Dry eye disease (DED) was surgically induced in wild-type (WT) or recombination-activating gene 1-knockout (*Rag1*KO) mice of both sexes through bilateral excision of the extraorbital lacrimal gland. Sham-operated animals were included as controls (Ct). **A)** Cumulative data and **B)** representative micrographs of corneal dextran-fluorescein uptake shown as the mean fluorescence intensity (MFI, see Methods). **C)** Number of proliferating (Ki67+) cells within the epithelial basal layer of corneal whole-mounts obtained 10 days after DED induction. **D)** Corneal mechanosensitivity and **E)** capsaicin thresholds in mice from both strains on days 0, 5, and 10 of DED induction. **F)** Representative micrographs and **G-H-I)** density of intraepithelial corneal innervation analyzed at three different levels by Alexa Fluor 488 anti-tubulin β3 staining: subapical (G, % area occupied by nerve endings) and mid-epithelial (H, count of nerve endings/field) nerve ending density and complexity of subbasal nerves (I, sum of intersections at all Sholl radii). All experiments were performed twice or more with 6 mice/group/experiment. To compare means, two-way ANOVA was used for A, D, and E (group and time) and C, G, H, and I (strain and treatment) with Sidak’s post hoc test. * indicates p<0.05, *** indicates p<0.001, **** indicates p<0.0001, and ns indicates not significant.

### 3. Adaptive immune deficiency prevents DED-associated corneal neuropathy development

Antigen-driven CD4^+^ T cell activation in a non-desiccated ocular surface is sufficient to damage corneal nerves(37). However, other factors must also determine whether corneal neuropathy occurs in the context of an ocular surface immune response since CD4^+^ T cell activation affects corneal nerves in herpetic keratitis and ocular graft-versus-host disease but not in allergic conjunctivitis mouse models(38). DED involves ocular desiccation and innate immune activation followed by an adaptive immune response. Thus, if local adaptive immunity is required for DED-associated corneal nerve dysfunction to develop is unknown. We first measured corneal sensitivity to mechanical and capsaicin stimulation to probe two different types of nerve fibers. At baseline, *Rag1*KO mice showed slightly reduced corneal mechanosensitivity (-4.4±0.9%, Figure S2A) and increased capsaicin sensitivity (Figure S2B). Sex had no effect (p=0.58, Figure S2C) on baseline mechanosensitivity thresholds in both strains. Therefore, when evaluating the effects of DED in each strain, the results were normalized to each strain-specific baseline (Figures 2D-E). Corneal mechanosensitivity progressively decreased and capsaicin sensitivity increased in WT DED mice(35, 39) while they remained unchanged in *Rag1*KO DED mice. Sub-analysis (Figure S2D, 3-way ANOVA with factors strain, sex, and time) revealed that strain had the largest effect (22.3% of total variation, p<0.0001) on mechanosensitivity followed by time (14.3% of total variation, p<0.0001) while sex had a small significant effect (2.9% of total variation, p=0.03). In agreement with their lower baseline corneal mechanosensitivity, control *Rag1*KO mice had lower corneal nerve density than control WT mice (Figures S2E-F). However, after 10 days of DED and consistent with the functional impairment, only WT DED mice displayed reduced nerve density at all levels of the corneal intraepithelial innervation (Figures 2F-I and S2G)(37, 39). By contrast, nerve density in *Rag1*KO DED mice did not vary. DED-induced corneal nerve changes in WT mice decreased in magnitude from the superficial subapical level to the deeper subbasal level whereas *Rag1*KO mice showed a similar non-significant trend (Figure S2H). Altogether, the data indicates that DED-associated corneal nerve damage requires an adaptive immune response to develop.

### 4. DED induces a trigeminal transcriptomic signature related to corneal nerve damage only in adaptive immune-sufficient mice

Corneal nerve fibers originate from somatosensory neurons in the trigeminal ganglion that may exhibit reactive gene expression changes in response to peripheral nerve injury(46–48). To further study the impact of adaptive immune deficiency on corneal neuropathy development, we analyzed the trigeminal transcriptional profiles by bulk RNA-Seq. First, we determined the effect of DED on trigeminal gene expression in each strain. After 10 days of DED induction, there were 172 differentially expressed genes (DEGs) in WT mice (77 up- and 95 down-regulated) but only 7 DEGs (5 up- and 2 down-regulated) in *Rag1*KO mice (Figures 3A-B and Dataset S1). In WT DED mice, up-regulated genes included *Atf3*, *Sprr1a*, and *Fos*, regeneration-associated genes induced by axonal injury in peripheral nervous system neurons(49), while down-regulated genes included nociceptive and mechanosensitive channels and voltage-gated ion channels. Comparison between WT and *Rag1*KO DED mice revealed 34 up- and 39 down-regulated genes in WT mice (Figure S3). Several up-regulated DEGs in WT mice corresponded to the absent adaptive immune response in the *Rag1*KO strain and were mostly B-cell related. The 3 DEGs that were up-regulated in WT DED mice relative to both control WT and *Rag1*KO DED mice were *Npy* (neuropeptide Y), *Dbp*, and *mt-Atp6*, all of which are associated with trigeminal neuropathic pain(47, 50). Down-regulated genes in WT DED mice included nociceptive and voltage-gated channels, some of which were also down-regulated relative to control WT mice. Gene set enrichment analysis identified 189 pathways in WT mice and 24 pathways in *Rag1*KO mice (Figure S4). In WT mice, the 10 most significantly activated pathways by DED pertained to pre- and postsynaptic translation, the humoral immune response, and antigen processing and presentation while the 10 most significantly suppressed pathways related to microtubule-based transport, exocytosis, detection of abiotic stimuli, and those involved in sensory perception of pain. In *Rag1*KO mice, the 10 most significant pathways activated by DED related to presynaptic translation and regulation of antigen processing and presentation whereas the 10 most suppressed ones were dendrite development, regulation of transporter activity, synapse assembly, and potassium ion transport, albeit the statistical significance was 10-fold lower than in WT mice. Finally, qPCR for 5 selected genes confirmed RNA-Seq findings in trigeminal WT samples (Figure S5). Altogether, these findings indicate that DED-induced trigeminal gene expression changes are larger and more related to corneal nerve damage in WT than in *Rag1*KO mice, which is consistent with the lack of corneal neuropathy development in the latter.

**Figure 3.**
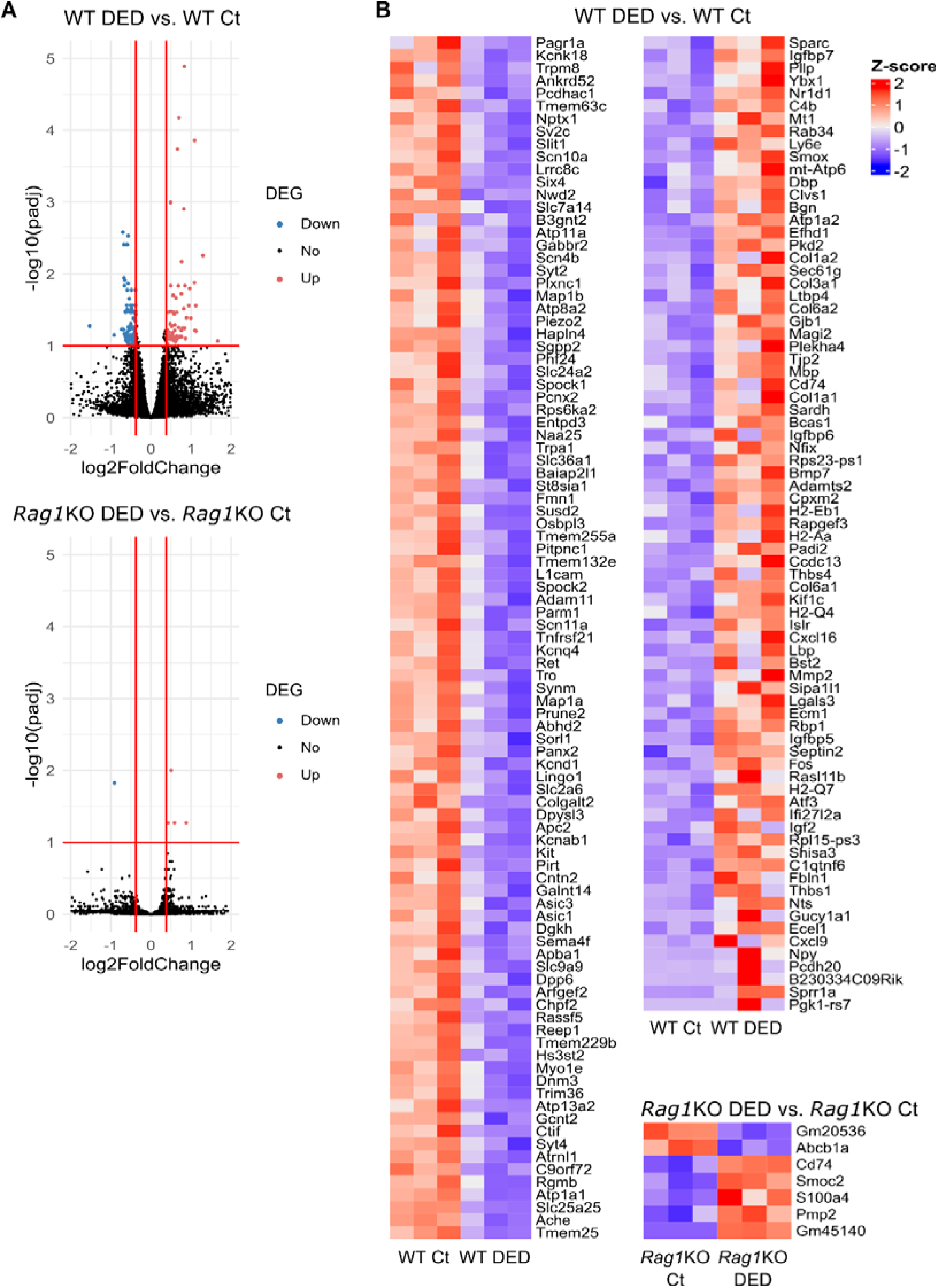
Gene expression changes induced by dry eye disease in the trigeminal ganglion. Bulk RNA-Seq analysis of trigeminal ganglia harvested 10 days after surgical induction of dry eye disease (DED) in wild-type (WT) or recombination-activating gene 1-knockout (*Rag1*KO) mice (female mice, n=3 per group). Differentially expressed genes (DEGs) were calculated between the sham-operated (Ct) and DED mice of each strain (fold change > 1.4, adjusted p-value < 0.1). **A)** Volcano plots of DEGs (red: up-, blue: down-regulated in DED mice, black: unchanged). **B)** Heatmaps (normalized counts, Z score) of DEGs.

### 5. DED leads to sustained proinflammatory cytokine levels in tears only in adaptive immune-sufficient mice

The adaptive immune response, and specifically its CD4^+^ T cells, sustain ocular surface inflammation in DED by releasing Th1 and Th17-associated pathogenic cytokines(13, 15, 19, 51–55). Because corneal nerves were spared in adaptive immune-deficient DED mice, we hypothesized that the specific lack of CD4^+^ T cells and their associated cytokines in this strain could explain the absence of corneal neuropathy. Therefore, we measured tear cytokine levels after 10 days of DED, once the CD4^+^ T cell response has fully ensued in WT mice(11, 53)(Figure 4). WT DED mice had increased levels of 6 of the 7 tested cytokines (IFN-γ, TNF, IL-6, IL-10, IL-17A, and IL-22) while *Rag1*KO DED mice showed no change. Subanalysis by sex of the WT strain data revealed that female DED mice had higher tear levels of IL-17A (p=0.03) and a similar trend for IFN-γ (p=0.07) while the other tested cytokines did not differ between sexes. Of note, the number of conjunctival macrophages and neutrophils (Figure S6) and adaptive immune cells in the trigeminal ganglia (Figure S7) of WT mice was not modified by DED. Our data indicates that the adaptive immune response plays a pivotal role in amplifying and sustaining ocular surface inflammation in DED through local cytokine production.

**Figure 4.**
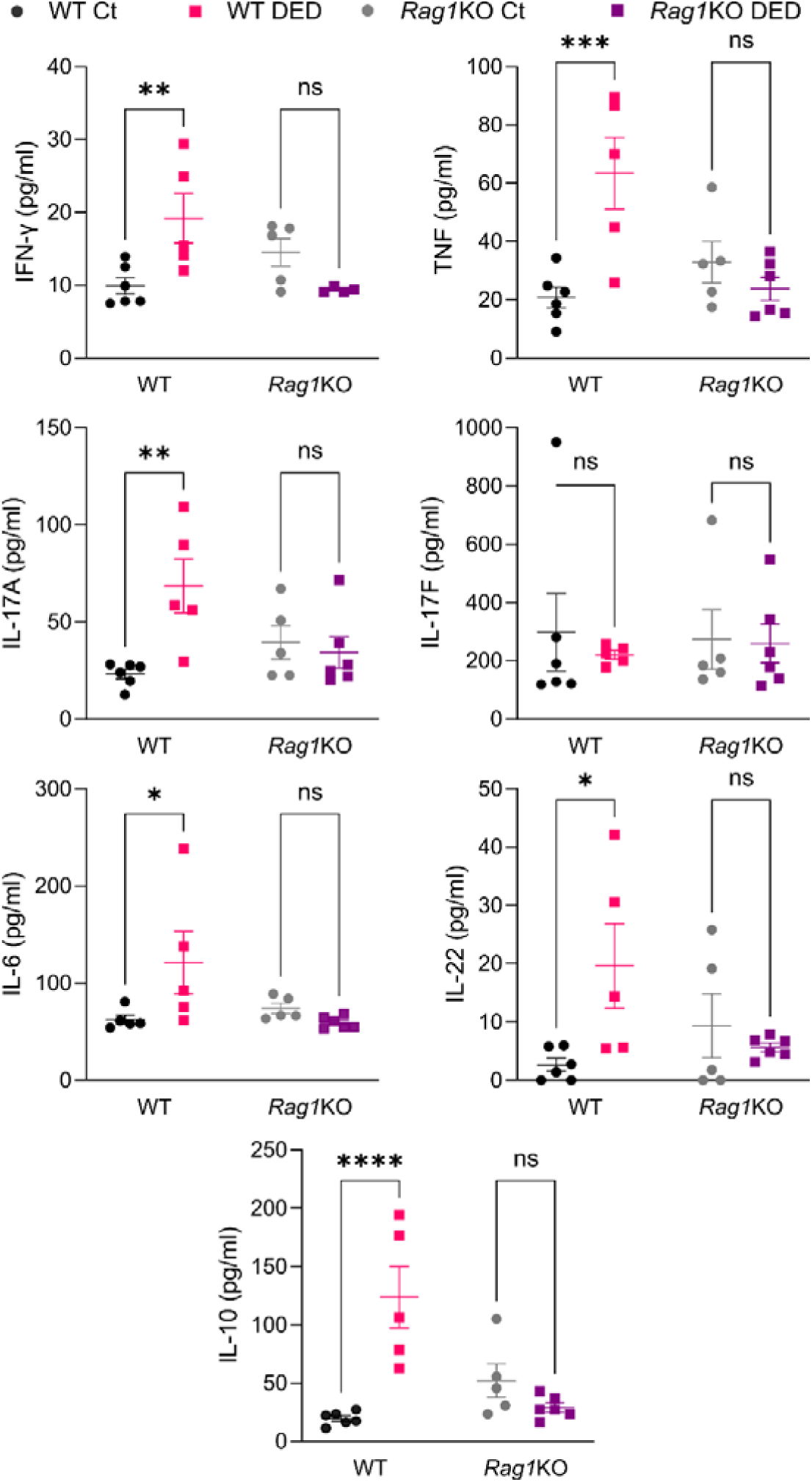
Cytokine levels in tear washings of wild-type and T cell-deficient mice with dry eye. Tear washings were collected after 10 days of dry eye disease (DED) induction in wild-type (WT) or recombination-activating gene 1-knockout (*Rag1*KO) mice of both sexes and analyzed by a bead-based multiplex assay for interferon (IFN)-γ, tumor necrosis factor (TNF), interleukin (IL)-6, -10, -17A, -17F, and -22 levels. Results from one representative experiment with 6 mice/group are shown (pg/ml, mean±SEM). Two-way ANOVA (treatment and strain) with Sidak’s post hoc test was used to compare means. * indicates p<0.05, ** indicates p<0.01, *** indicates p<0.001, **** indicates p<0.0001, and ns indicates not significant.

### 6. Adoptive transfer of CD4^+^ T cells from DED mice recapitulates ocular phenotype and corneal nerve damage but not epitheliopathy in the recipients

Our results so far suggested that the adaptive immune response drives corneal nerve damage in DED. Since isolated ocular CD4^+^ T cell activation is capable of inducing corneal neuropathy(37), we hypothesized that the lack of CD4^+^ T cells in *Rag1*KO DED mice was responsible for their resistance to develop corneal nerve damage. To test this, splenic CD4^+^ T cells from WT mice were isolated 10 days after sham or DED-inducing surgery and transferred into sex-matched naïve *Rag1*KO mice, which were then assessed weekly for DED pathology (Figure 5A). First, we validated the model. Control or DED-induced CD4^+^ T cells did not significantly differ in the proportion of Th1 or Th17 cells, although there was a trend towards more IFN and IL-17A production in the latter group (Figure S5A). After 4 weeks, the lymph nodes of the recipient *Rag1*KO mice only had CD4^+^ T cells, ruling out any potentially confounding effect from contaminating CD8^+^ T or B cells that may proliferate rapidly in the lymphopenic host (Figure 5B). As previously reported(11), DED-induced CD4^+^ T cells homed more efficiently to the conjunctiva than control CD4^+^ T cells (Figure 5C and S5B). Despite the conjunctival infiltration, the external eye phenotype remained unchanged as in the surgically induced DED model (Figures 5D and 1B). Finally, the pathogenicity of the DED-induced CD4^+^ T cells was confirmed by the decreased tear production (Figure 5E) and reduced conjunctival goblet cell density (Figures 5F-G) in the recipient *Rag1*KO mice. However, there was no detectable extraorbital lacrimal gland infiltration in either recipient group and tear cytokine levels were similar to those in control *Rag1*KO mice (Figures S9-10 and 4). Thus, the adoptive transfer of DED-induced CD4^+^ T cells reproduced all the non-corneal findings observed in WT DED mice in the *Rag1*KO recipients.

**Figure 5.**
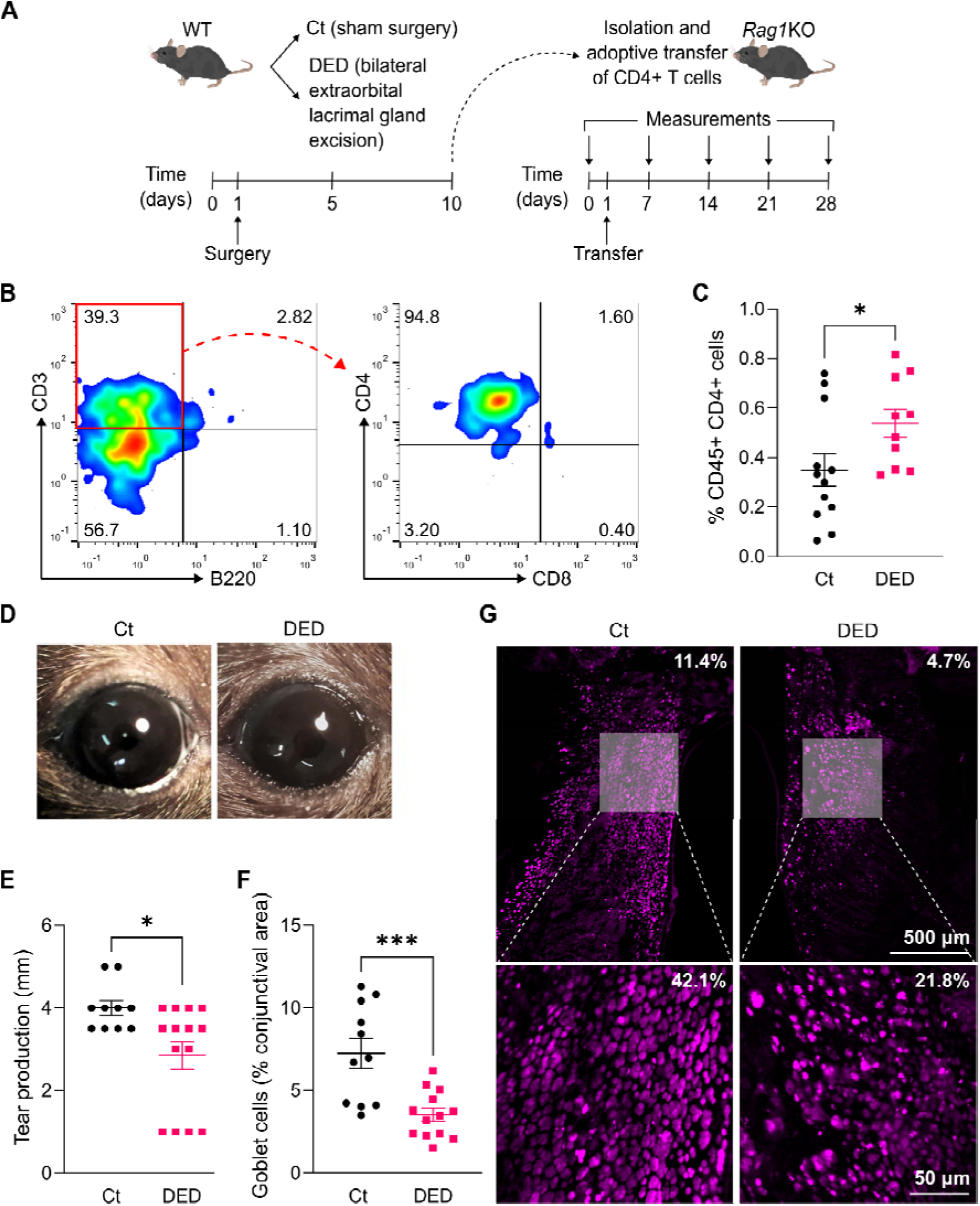
Adoptive transfer of dry eye-induced CD4^+^ T cells reproduces ocular surface phenotype. **A)** Experimental design: CD4^+^ T cells were isolated from the spleens and lymph nodes of wild-type (WT) mice of both sexes 10 days after surgical induction of dry eye disease (DED) and then adoptively transferred into sex-matched recombination-activating gene 1-knockout (*Rag1*KO) mice, which were evaluated over the course of 4 weeks. Sham-operated WT mice were used as a source of control (Ct) CD4^+^ T cells. **B)** Flow cytometry of cervical lymph node cells (representative example) obtained from a CD4^+^ T cell-reconstituted *Rag1*KO mouse 4 weeks after transfer. **C)** Conjunctival CD4^+^ T cells in *Rag1*KO mice 4 weeks after transfer as assessed by flow cytometry (gated on total viable cells). **D)** External eye appearance (representative micrographs) of *Rag1*KO recipients 4 weeks after transfer. **E)** Tear production in Ct- and DED-induced CD4^+^ T cell recipients measured on days 25-27 after transfer. **F)** Total conjunctival area occupied by goblet cells in Ct- and DED-induced CD4^+^ T cell recipients 4 weeks after transfer. **G)** Representative micrographs of wheat germ agglutinin-stained goblet cells in conjunctival whole-mounts used for quantification. Low-magnification micrographs (top) of whole-mounted conjunctival strips used for measurements shown in F, with their corresponding high-magnification insets of the goblet cell-rich area (bottom) provided only as a reference. The actual goblet cell-occupied area is shown in the top right corner of each image. All experiments were performed twice or more with 6 mice/group/experiment. Student’s t test was used to compare means. * indicates p<0.05 and *** indicates p<0.001.

Next, we assessed the capacity of the pathogenic CD4^+^ T cells to cause corneal epithelial damage in the recipient mice in the absence of desiccating stress. Neither group of recipient mice (control and DED-induced CD4^+^ T cells) showed a change from baseline levels of corneal dextran-FITC uptake (Figures 6A-B). Sub-analysis (Figure S6A, 3-way ANOVA with factors transferred cells, sex, and time) confirmed that no factor had a significant effect on corneal epitheliopathy development (<5% of total variation, p>0.05). Moreover, both groups had comparable numbers of proliferating corneal epithelial cells (Figures 6C and S6B) that were restricted to the basal layer (Figure S6C), as observed in homeostatic conditions. Thus, these results indicate that in the absence of desiccating stress, CD4^+^ T cells from DED mice are not sufficient to induce corneal epitheliopathy in *Rag1*KO recipient mice.

**Figure 6.**
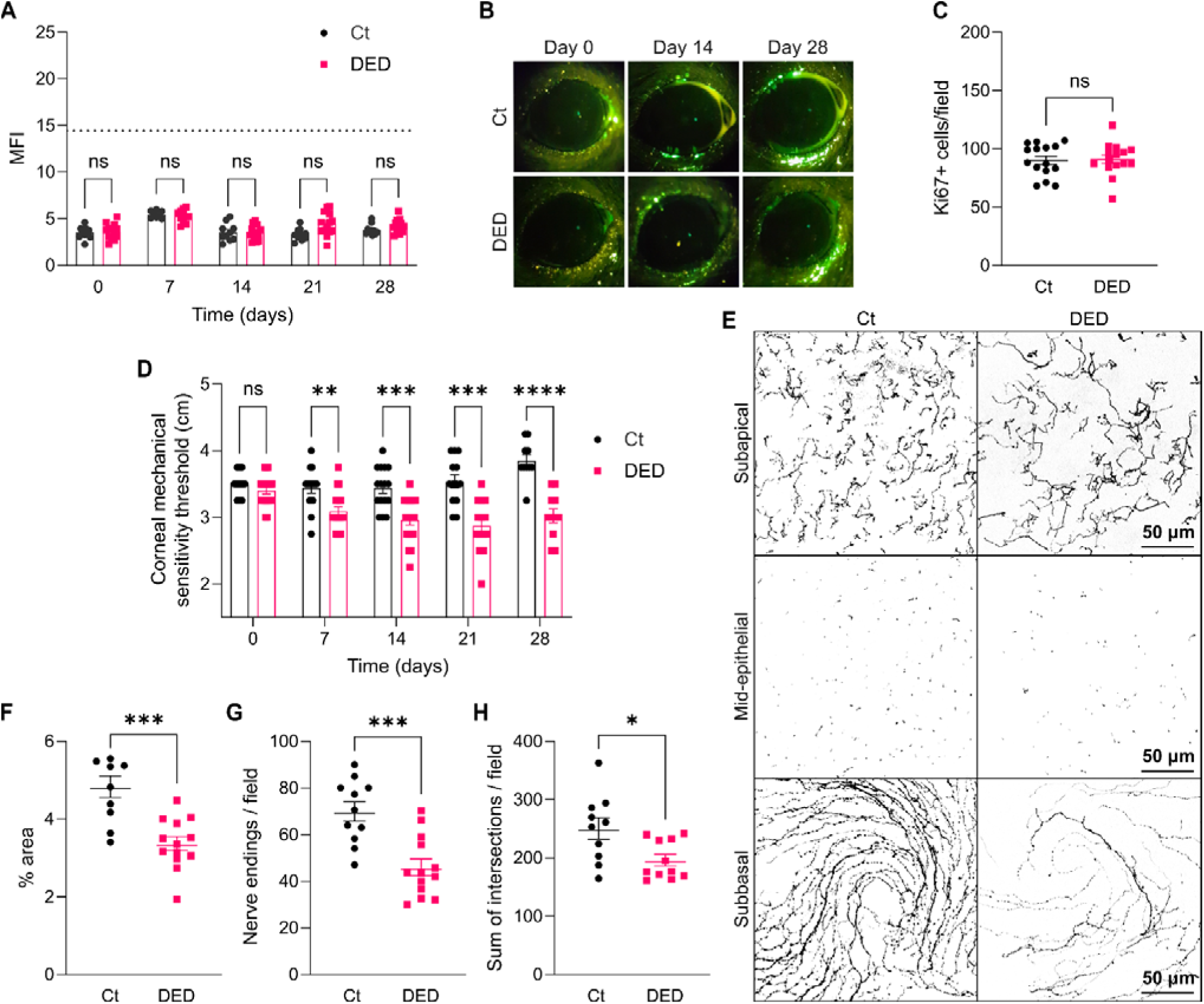
Adoptive transfer of dry eye-induced CD4^+^ T cells reproduces corneal neuropathy but not epitheliopathy. The experimental design was described in detail in Figure 5A and involved the adoptive transfer of control (Ct) or dry eye disease (DED)-induced CD4^+^ T cells from WT mice into recombination-activating gene 1-knockout (*Rag1*KO) mice. **A)** Cumulative and **B)** representative micrographs of corneal dextran-fluorescein uptake in Ct- and DED-induced CD4^+^ T cell recipients. Data shown as the mean fluorescence intensity (MFI, see Methods). The dotted line represents the mean corneal uptake observed in WT DED mice. **C)** Quantification of proliferating cells within the epithelial basal layer of corneal whole-mounts obtained from Ct- and DED-induced CD4^+^ T cell-recipient mice. **D)** Corneal mechanosensitivity thresholds in Ct- and DED-induced CD4^+^ T cell recipients. **E)** Representative micrographs and **F-H)** density of intraepithelial corneal innervation analyzed at three different levels by Alexa Fluor 488 anti-tubulin β3 staining: subapical (F, % area occupied by nerve endings) and mid-epithelial (G, count of nerve endings/field) nerve ending density and complexity of subbasal nerves (H, sum of intersections at all Sholl radii). All experiments were performed twice or more with 6 mice/group/experiment. To compare means, two-way ANOVA (donor group and time) with Sidak’s post hoc test was used for A and D, and Student’s t test was applied in C, F, G, and H. * indicates p<0.05, ** indicates p<0.01, *** indicates p<0.001, **** indicates p<0.0001, and ns indicates not significant.

By contrast, corneal mechanosensitivity significantly decreased in the DED-induced CD4^+^ T cell-recipient mice as early as 7 days after transfer (Figure 6D) and did not worsen significantly from this time point. Sub-analysis (Figure S6D, 3-way ANOVA with factors group, sex, and time) revealed that sex had no effect (p=0.88) on mechanosensitivity, in contrast with the findings from the surgical DED model (Figure S2A). Consistently, corneal intraepithelial nerve density at all levels was lower 28 days post-transfer in the DED-induced CD4^+^ T cell-recipient mice (Figures 6E-H). The pattern of change in this group was similar to that in WT DED mice: a larger relative reduction in the subapical nerve endings than in the subbasal corneal nerves (Figure S6E). Finally, we compared the transcriptomic signatures in the trigeminal ganglia of female Ct- and DED-induced CD4^+^ T cell-recipient mice 28 days after transfer (Figure S12). There were 658 DEGs (180 up- and 478 down-regulated) in DED-induced CD4^+^ T cell-recipient mice (Dataset S2). Eight of these genes were also differentially expressed in WT DED mice, including *Ache* (acetylcholinesterase) and *Asic3* (acid-sensing ion channel subunit 3) which have been linked to trigeminal pain perception(56, 57). Gene set enrichment analysis identified 199 pathways (Figure S13). The 10 most significantly activated pathways in DED-induced CD4^+^ T cell-recipient mice were related to innate immune activation and type I immunity while the 10 most significantly suppressed pathways were associated with oxidative phosphorylation, glycolysis, and mitochondrial activity.

Altogether these results show that adoptive transfer of pathogenic DED-induced CD4^+^ T cells reproduces corneal neuropathy and trigeminal neuroinflammation but not epitheliopathy in recipient mice.

### 7. Adaptive immune-deficient mice reconstituted with naive CD4^+^ T cells gain the ability to develop corneal nerve damage upon DED induction

As a proof-of-concept that CD4^+^ T cells drive corneal nerve damage in DED, we tested if reconstitution of *Rag1*KO mice with naive CD4^+^ T cells was sufficient to enable corneal neuropathy development upon surgical DED induction. We adoptively transferred CD4^+^ T cells from naïve WT mice into *Rag1*KO mice that, one week later, underwent either sham- or DED-inducing surgery (Figure 7A). First, we confirmed that T cell reconstitution in the cervical lymph nodes was effective and limited to CD4^+^ T cells (Figure 7B), and then, we tested for the development of corneal epitheliopathy and neuropathy after disease induction. Corneal barrier disruption progressively increased on days 5 and 10 of DED induction comparably to WT and *Rag1*KO DED mice (Figures 7C-D and 2A). But contrasting with *Rag1*KO mice lacking T and B cells, CD4^+^ T cell-reconstituted *Rag1*KO DED mice developed progressive corneal mechanosensitivity loss and exhibited reduced corneal nerve density at all intraepithelial levels (Figures 7E-I). The pattern of change showed a larger relative reduction in the subapical nerve endings than in the subbasal corneal nerves (Figure 7J). These results altogether show that reconstitution of *Rag1*KO mice with naïve CD4^+^ T cells from WT mice followed by surgical DED induction enables corneal neuropathy onset and progression, thus confirming that CD4^+^ T cells are required for corneal nerve damage to develop in DED.

**Figure 7.**
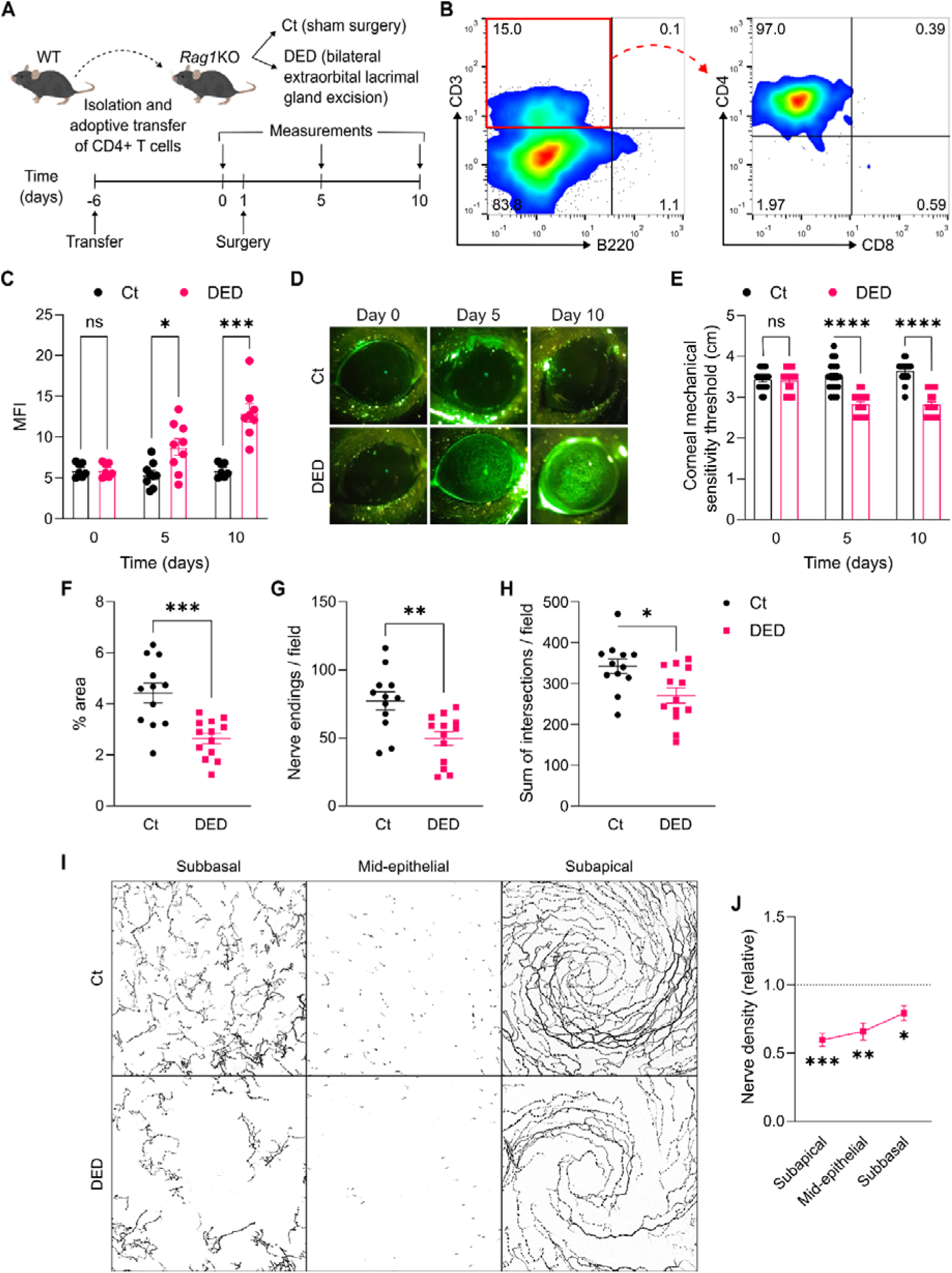
Reconstitution of T cell-deficient with CD4^+^ T cells enables corneal neuropathy development upon dry eye induction. **A)** Experimental design: CD4^+^ T cells were isolated from the spleens and lymph nodes of wild-type (WT) mice of both sexes and then adoptively transferred into sex-matched recombination-activating gene 1-knockout (*Rag1*KO) mice, which one week later underwent either sham (Ct) or dry eye disease (DED)-inducing surgery and were evaluated over the course of 10 days. **B)** Flow cytometry of cervical lymph node cells (representative example) obtained from a CD4^+^ T cell-reconstituted *Rag1*KO mouse 10 days after surgery. **C)** Cumulative data and **D)** representative micrographs of corneal dextran-fluorescein uptake in Ct and DED CD4^+^ T cell-reconstituted *Rag1*KO mice. Data shown as the mean fluorescence intensity (MFI, see Methods). **E)** Corneal mechanosensitivity thresholds in Ct and DED CD4^+^ T cell-reconstituted *Rag1*KO mice. **F-H)** Quantification and **I)** representative micrographs of intraepithelial corneal innervation analyzed at three different levels by Alexa Fluor 488 anti-tubulin β3 staining: subapical (F, % area occupied by nerve endings) and mid-epithelial (G, count of nerve endings/field) nerve ending density and complexity of subbasal nerves (H, sum of intersections at all Sholl radii). **J)** Distribution of change in nerve density over the three levels of intraepithelial corneal innervation in DED CD4^+^ T cell-reconstituted *Rag1*KO mice (expressed relative to Ct CD4^+^ T cell-reconstituted mice). All experiments were performed twice or more with 6 mice/group/experiment. To compare means, two-way ANOVA with Sidak’s post hoc test was used in C and E (treatment and time), and Student’s t test was applied in F, G, H, and J. * indicates p<0.05, ** indicates p<0.01, *** indicates p<0.001, **** indicates p<0.0001, and ns indicates not significant.

## Discussion

Our current understanding of DED involves dysfunctional tear film-instigated corneal epitheliopathy and neurosensory abnormalities as defining clinical features and ocular surface inflammation as a key driver of the disease process(3, 4, 6). Contrasting with the pathogenesis of corneal epitheliopathy in DED, the mechanisms underlying corneal nerve damage are poorly understood(32). A CD4^+^ T cell-driven adaptive immune response perpetuates ocular surface inflammation(8) and thus interfering with CD4^+^ T activation is the target of several approved therapies. However, whether CD4^+^ T cells are required for either DED-associated corneal epitheliopathy or neuropathy to occur is undetermined. Herein we show that while DED-associated corneal epitheliopathy develops unimpeded in the absence of CD4^+^ T cells, corneal nerve damage does not. Thus, our results indicate that the mechanisms behind these two defining aspects of the disease are different and that therapies that target one may not necessarily tackle the other.

Our findings on corneal epitheliopathy expand our knowledge of DED pathogenesis(7, 8). The contribution of the adaptive immune response to disease progression was established by showing that CD4^+^ T cells from DED mice reproduce lacrimal gland- and conjunctival infiltration in T cell-deficient recipients(11). Subsequent investigations demonstrated the pathogenic relevance of type 1 (Th1) and type 3 (Th17) immunity in many aspects of DED including corneal epitheliopathy(13, 15, 19, 25, 28, 51–55, 58), but none indicated that corneal epitheliopathy does not occur in the absence of CD4^+^ T cell activation. Consistently, local depletion of antigen presenting cells prevented conjunctival CD4^+^ T cell infiltration and goblet cell loss but no effect on corneal epitheliopathy-related signs was described(13). Our results agree with these reports: adoptive transfer of DED-induced CD4^+^ T cells reproduces lacrimal gland dysfunction, conjunctival infiltration, and goblet cell loss in T cell-deficient recipient mice (Figures 5-6). Also, the adoptive transfer of effector memory CD4^+^ T cells from female DED mice into naïve recipients was shown to cause more severe corneal epitheliopathy than naïve or effector CD4^+^ T cells but only after desiccating stress exposure(53). Consistently, female WT mice in our study developed more severe corneal epitheliopathy than female *Rag1*KO mice on day 5 of DED induction, but the differences disappeared by day 10 among strains and sexes (Figure S1). Thus, our findings demonstrate that CD4^+^ T cells are dispensable for DED-associated corneal epitheliopathy onset, but these cells may accelerate or worsen this aspect of the disease in females.

Nonetheless, our data does not imply that type 1 and 3 immunity are not involved in DED-associated corneal epitheliopathy. IFN-γ and IL-17 are pathogenic to the corneal epithelium(19, 22, 24), and conjunctival NK cells and other innate immune cells represent a significant source of these cytokines in the ocular surface(25, 28–31). Conjunctival IFN-γ and Th1-associated chemokines increase in *Rag1*KO mice after 5 days of desiccating stress but return to baseline levels by day 10(29). Conjunctival goblet cell loss is also detectable in *Rag1*KO mice after 5 days of DED induction, indicating that early IFN-γ production by innate immune cells in the ocular surface is sufficient to cause this typical DED finding(21, 29). The present study complements these reports by showing that DED-associated conjunctival goblet cell apoptosis, which is caused mainly by IFN-γ(12, 21), occurs unhindered in the absence of CD4^+^ T cells (Figures 1D-E). Increased tear levels of IFN-γ and IL-17 were not detected in T cell-deficient mice after 10 days of DED induction (as opposed to WT mice, Figure 5), indicating that CD4^+^ T cells are required for sustained production of these pathogenic cytokines(53). Thus, our findings also suggest that the early innate components of type 1 and type 3 responses are competent to prime the proinflammatory ocular surface state and conjunctival goblet cell dysfunction that contribute to corneal epithelial damage.

By contrast, the onset of corneal nerve damage in DED depends on the presence of CD4^+^ T cells. Since corneal epithelial cells ensheath nerve fibers(34), it is striking that desiccation-induced pathologic changes in the former may occur while the latter remain unaffected in the absence of CD4^+^ T cells. However, we have found that the converse is also true: corneal nerves may be damaged by CD4^+^ T cell activation in the ocular surface while the corneal epithelium remains unaffected in the absence of desiccating stress(37). Thus, the present findings extend the notion of an epithelial-neural divide in corneal pathology to DED. This might translate into variations in the immune and non-immune components contributing to the diverse presentations of the disease: little difference in corneal epithelial changes but worse corneal nerve alterations were reported between Sjögren’s and non-Sjögren’s syndrome DED patients(59, 60). The sensitivity of corneal nerves to CD4^+^ T cell activation could also explain the corneal neuropathy findings that correlate with the severity of the systemic autoimmune disease in rheumatoid arthritis patients(61) or that are observed after recovery in COVID-19 patients(62). Of note, corneal nerves are also affected during HIV infection, when CD4^+^ T cell function is impaired, by viral replication and immune activation in the trigeminal ganglion(63, 64). Thus, trigeminal neuroinflammation *per se* may also impact corneal nerve endings independently of CD4^+^ T cell activity. In line with this, increased corneal nociceptor signaling is associated with trigeminal neuroinflammation and propagation of DED-induced corneal neuropathy(39, 65).

The trigeminal transcriptomic signatures observed after 10 days of DED induction in WT mice but not in *Rag1*KO mice are consistent with the extent of corneal nerve damage and correspond to the common transcriptomic signatures of peripheral nerve injury and regeneration and neuropathic pain (Figures 3, S3-4). Based on the DEG numbers and expression fold-change, the magnitude of the DED-induced trigeminal transcriptomic signature is smaller than in other models of trigeminal nerve injury(46–48, 66). However, most models involve whole nerve ligation or injection of inflammatory agents whereas DED targets the peripheral nerve endings of only 2% of trigeminal neurons(32). A combined analysis of bulk- and single-cell RNA-Seq studies of dorsal root ganglia in neuropathic pain or nerve injury models showed that most gene expression changes take place in nociceptor neurons and that the reduced magnitude of expression changes in whole ganglion studies probably relates to the diluting effect of other cell types(67). In light of this, the qualitatively comparable trigeminal transcriptomic signature observed in our study probably indicates a pain maintenance period as that reported after 10 days of partial infraorbital nerve transection(66). Functional impairment of mechanical sensitivity, capsaicin hypersensitivity, spontaneous ocular pain, and morphological alterations in corneal nerve fibers are well established at this time point in our model(39). These findings are commensurate with the gene expression changes found in the trigeminal ganglia of WT DED mice, which are associated with a neuropathic pain state in other models(66, 67). Supporting this hypothesis, the up-regulation of the macrophage-specific marker *Cd74* in both WT and *Rag1*KO DED mice probably reflects the increase in neuron-associated macrophages in the trigeminal ganglion upon axonal injury that relates to neuropathic pain(68, 69). Thus, our data support the notion that adaptive immune response contributes to the development of neuropathic pain in DED.

Although our study does not delve into the exact process by which CD4^+^ T cells promote corneal neuropathy, the persistently elevated tear cytokine levels probably play a role (Figure 5). CD4^+^ T cells coordinate immune responses in tissues mostly by secreting cytokines that have diverse effects on innate immune cells and non-immune cells(70, 71). In DED, conjunctival CD4^+^ T cells increase in number and constitute a likely source of the Th1 and Th17 cytokines that reach the cornea through the tear film. We observed more severe disruption of the superficial subapical nerve endings than of subbasal nerve fibers in the three models tested, one of which lacked ocular desiccation and relied entirely on CD4^+^ T cell activation for pathogenicity. This pattern of nerve fiber impairment is compatible with a soluble factor diffusing from the tear film into the corneal epithelium, which could be an action-at-distance effect of conjunctival CD4^+^ T cells. Supporting this hypothesis, DED-induced CD4^+^ T cells preferably homed to the conjunctiva (Figure 6C). Nonetheless, a smaller number of T cells patrol the corneal epithelium under homeostatic conditions in humans and mice(72) and may even establish corneal tissue residency in pathologic settings(73). Therefore, it remains to be established whether CD4^+^ T cells must enter the cornea to exert their pathogenic action on corneal nerves. Alternatively, and even though these populations do not change in number in our DED model, modulation of conjunctival myeloid cells(74) or trigeminal resident immune cells(75, 76) could represent other mechanisms by which CD4^+^ T cells damage corneal nerve fibers indirectly. The trigeminal transcriptomic signature in DED-induced CD4^+^ T cell-recipient mice included innate immune-driven neuroinflammation, cytokine responses, and metabolic impairment, suggesting a local pathogenic effect within the trigeminal ganglion. Intriguingly, tear cytokine levels in these CD4^+^ T cell-recipient mice did not rise above those in control *Rag1*KO mice (Figure S10) despite their corneal nerve impairment, which may be due to a limitation of the technique, a matter of timing, or even a difference in the underlying pathophysiology between the two models.

The lower corneal mechanosensitivity, higher capsaicin sensitivity, and reduced nerve density at baseline in *Rag1*KO mice (Figures S2A-B, S2E-F) indicate that the lack of an adaptive immune system has a developmentally significant effect on corneal nerves. There are known effects of adaptive immunity on olfactory function(77), and more specifically of CD4^+^ T cells, on anxiety-like behavior in mice(78). While it could be argued that these baseline differences and not the lack of CD4^+^ T cells in *Rag1*KO mice might be responsible for their resistance to DED-induced corneal neuropathy, the adoptive transfer and CD4^+^ T cell-restricted reconstitution experiments favor the latter (Figures 2, 6-7). Moreover, repopulation of adult *Rag1*KO mice with naïve CD4*^+^*T cells does not revert their impaired emotional behavior(78) but does render them susceptible to DED-induced corneal neuropathy (Figure 7). By the same token, the absence of B cells, CD8*^+^* T cells, and NKT cells being responsible for the lack of a corneal neural phenotype in *Rag1*KO DED mice is ruled out by the recapitulation of corneal neuropathy in the aforementioned experiments (Figures 2, 6-7). Nonetheless, our findings do not imply that adaptive immune cells other than CD4*^+^* T cells do not contribute to non-neural aspects of DED, as already demonstrated by others(79–81). Another intriguing finding was the lack of lacrimal gland infiltration after the adoptive transfer of DED-induced CD4^+^ T cells (Figure S9). We hypothesize that the resulting mild tear hyposecretion was due to dysregulated tearing reflexes caused by the impaired corneal nerve sensory function (Figures 5E, 6)(32), and not by a primary lacrimal gland defect. Thus, this model could represent the short tear film breakup time-subtype of DED seen in patients with normal tear secretion, no corneal epitheliopathy, and neuropathic pain(4).

Finally, female sex is the most influential risk factor for DED in patients(82, 83), and our findings also shed light on this association: female mice develop worse DED in terms of corneal epithelial and neural impairment in part due to greater pathogenic activity of CD4^+^ T cells in this sex (Figures S1-S2). However, this difference cannot be solely attributed to these cells because female mice adoptively transferred with CD4^+^ T cells from female DED mice do not develop worse corneal neuropathy than males (Figure S6A). This indicates that sex influences other pathogenic factors that are triggered by desiccation in the ocular surface. A reduced regulatory role of ocular surface neutrophils in female mice has been linked to an amplified effector CD4^+^ T cell response in the ocular surface and worse DED phenotype(84). Alternatively, the more severe corneal mechanosensitivity impairment in female DED mice (Figure S2A) may involve greater susceptibility of corneal nerves to immune-mediated damage. Supporting this idea, corneal neural repair mechanisms are conditioned by sex as female mice exhibit faster corneal neuroregeneration than males(85).

In summary, the present study enhances our understanding of the pathophysiology of DED-associated corneal epithelial and nerve damage by showing that desiccation-induced changes in the distal subapical endings of corneal nerves require the activation of CD4^+^ T cells while those in corneal epithelial cells occur independently of the adaptive immune response. One limitation of our study (and of DED animal models in general) is that the onset of desiccation is abrupt, as opposed to DED patients who typically suffer a progressive impairment of the tear film, which eventually elicits CD4^+^ T cell-sustained ocular surface inflammation that aggravates the disease(86). Thus, ocular surface CD4^+^ T cell activity may contribute indirectly to corneal epithelial damage during the initial phases of the disease in patients, as suggested by the faster tempo of corneal epitheliopathy in female WT mice (Figure S1A). By contrast, the resistance of corneal nerves to experimental desiccation in the absence of CD4^+^ T cells indicates that corneal neuropathy in DED is mostly immune-mediated. We have previously shown that corneal nerves are particularly sensitive to the activation of Th1 CD4^+^ T cells in the ocular surface in the absence of desiccating stress(37), and on the other hand, that signaling through transient receptor potential vanilloid-1 channels in the corneal tissue is required to propagate desiccation-initiated corneal nerve damage in DED(39). Therefore, the sustained levels of Th1 cytokines from CD4^+^ T cells likely prime corneal nerve changes that later propagate proximally through overactivation of transient receptor potential vanilloid-1 channels. Overall, our findings have implications for current DED management because they help explain why therapies aimed at reducing or controlling CD4^+^ T cell recruitment and activation in the ocular surface may be more impactful on certain manifestations of the disease and not on others. At the same time, the present study also highlights the need for further research to elucidate the precise mechanisms underlying corneal epithelial and nerve damage in DED and to identify potential therapeutic targets for these conditions.

## Materials and methods

See Supporting Information Text.

## Data availability

The data underlying Figures 3, S3, S4, S12 and S13 are openly available in ArrayExpress (https://www.ebi.ac.uk/biostudies/arrayexpress), accession E-MTAB-13945.

## Funding

This work was supported by research grants from Wellcome Trust (221859/Z/20/Z) and Agencia Nacional de Promoción Científica y Tecnológica (FONCyT PICT 2018-02911, PICT 2020-00138, PICT 2021-00109) awarded to JGG.

## Notes

https://www.ebi.ac.uk/biostudies/arrayexpress

### Competing Interest Statement

The authors have declared no competing interest.

### Summary of Updates

Text revised, 7 supplemental figures added, 1 dataset added

