## Supporting Information for "CD4^+^ T cells drive corneal nerve damage but not epitheliopathy in an acute aqueous-deficient dry eye model"

**Affiliations:**
1 Innate Immunity Laboratory, Institute of Experimental Medicine (CONICET/National Academy of Medicine of Buenos Aires), Buenos Aires, Argentina

2 Flow Cytometry Unit, Institute of Experimental Medicine (CONICET/National Academy of Medicine of Buenos Aires), Buenos Aires, Argentina

3 Confocal Microscopy Unit, Institute of Experimental Medicine (CONICET/National Academy of Medicine of Buenos Aires), Buenos Aires, Argentina

4 Translational Clinical Research Program, Hospital del Mar Research Institute, Barcelona, Spain

5 Ocular Surface Center, Baylor College of Medicine, Houston, Texas, USA

Corresponding author: Jeremías G Galletti


**This PDF file includes:**

Supporting text
Figures S1 to S13
Tables S1 to S2
Legends for Datasets S1 to S2
SI References

**Other supporting materials for this manuscript include the following:**

Datasets S1 to S2

**Supporting Text**

### Extended Methods

**Mice.** C57BL/6 (C57BL/6NCrl) mice were originally obtained from Charles River Laboratories (Wilmington, MA, USA) and recombination-activating gene 1 (*Rag1*)-knockout (*Rag1*KO (B6.129S7-Rag1tm1Mom/J, JAX stock #002216) mice were purchased from The Jackson Laboratory (Bar Harbor, ME, USA). Mice were bred and maintained at the Institute of Experimental Medicine's conventional animal facility. All mice were 6-8 weeks old at the beginning of the experiments and both male and female mice were included. All protocols were approved by the Institute of Experimental Medicine animal ethics committee (approval #084/2020) and adhered to the Association for Research in Vision and Ophthalmology Statement for the Use of Animals in Ophthalmic and Vision Research. Since DED is more prevalent and more severe in women(1), and consistently, female mice develop worse DED than their male counterparts(2), we performed all experiments using equal numbers of female and male animals to consider this difference and we included sex as a variable in the analysis.

**Reagents and antibodies.** Table S1 lists all antibodies and the most significant reagents. Unless otherwise specified, all chemical and biological reagents were from Sigma-Aldrich (Buenos Aires, Argentina).

**Lacrimal gland excision surgery.** Mice were anesthetized by i.p. injection of ketamine (100 mg/kg) and xylazine (10 mg/kg) and placed on a heated pad. Excision surgery was performed as previously reported(3).

**Tear production, corneal epithelial barrier function, and mechanical and capsaicin sensitivity measurements.** All DED-related measurements were performed as previously described(3, 4). Corneal fluorescein uptake of dextran-fluorescein isothiocyanate (average molecular weight 3000–5000, 10 mg/ml in PBS) was used as an indicator of epithelial barrier integrity. Corneal mechanical thresholds were determined using nylon 6-0 monofilament(3–6) in the morning (8–11 AM) before any other experimental manipulation.

**Adoptive transfer of $CD4^+$ T cells.** $CD4^+$ T cells (>98% purity) were isolated by negative selection with the aid of magnetic beads (MojoSort Mouse CD4+ T Cell Isolation Kit, BioLegend #480033) from splenocyte suspensions and each *Rag1*KO mouse received $1x10^6$ cells/0.5 ml PBS(3, 4).

**Lymph node cells and flow cytometry analysis.** Mechanically dissociated cervical lymph node cells were stained for surface markers B220, CD3, CD4, and CD8(3, 4). The entire cell suspension resulting from one mouse was stained and acquired as one independent sample on a Cyflow space cytometer (Partec, Germany) and analyzed using FlowJo software (FlowJo v10.3, Treestar, Ashland, OR, USA).

**Preparation and flow cytometry analysis of conjunctival and trigeminal cell suspensions.** Following euthanasia and cardiac perfusion with PBS to remove contaminating blood cells, conjunctivas and trigeminal ganglia were collected in serum-free RPMI 1640 medium, minced with the aid of scissors, and then digested with collagenase and DNAse, filtered, stained, and fixed as previously described(3, 4). The entire cell suspension resulting from one mouse was stained and acquired as one independent sample.

**Collection of eye tissue for imaging.** After euthanasia, the conjunctival tissue of each eye was excised as two strips (superior and inferior) under a dissection microscope and collected in an ice-cold fixative solution. Immediately after, enucleation was performed by gently proptosing the eye globe and cutting the optic nerve with curved scissors. The two eyes of each mouse were collected in ice-cold fixative solution. Mice were euthanized one at a time so that all ocular tissue was collected within 5 min of the time of death to ensure adequate corneal nerve preservation(7).

**Conjunctival sample staining, spinning disk scanning microscopy, and image analysis.** Excised conjunctivas were fixed in pre-chilled 4% formaldehyde in PBS for 1 h, then washed 3 times with PBS, and stored at 4°C until processing. Conjunctiva samples were simultaneously blocked and permeabilized with a 1% BSA and 0.3% Triton X-100 solution overnight at 4°C, then washed, and stained overnight with wheat germ agglutinin Alexa Fluor® 647 Conjugate. The stained conjunctivas were washed three times in PBS for 20 min, mounted in Aqua-Poly/Mount (PolySciences), and stored at 4°C until imaged. Image acquisition was performed with an Olympus IX83 inverted motorized microscope (Olympus, Tokyo, Japan) equipped with a UPlanSapo 10x/0.4 objective and a Disk Scanning Unit. Composite images (3-μm step size Z-stacks) spanning the entire conjunctival strips were obtained using the multiple image alignment module of the cellSens Dimensions software (Olympus) to account for the non-uniform distribution of goblet cells within the conjunctiva(8). To assess the goblet cell area, maximum intensity projections including all epithelial slices within the stacks were created using ImageJ. Then, the entire conjunctival surface was demarcated as the area of interest using the polygon selection tool, a background correction (20-pixel rolling ball radius) was applied, the image was thresholded, and finally, the percentage of the selected area occupied by the wheat germ agglutinin-derived signal was measured by the software.

**Corneal immunostaining and confocal laser scanning microscopy acquisition.** Eyes were processed as previously described(3, 4). After blocking, the dissected corneas were stained overnight with anti-tubulin β3, anti-mouse/human Ki-67, and anti-mouse/human CD324 (E-cadherin) antibodies. Image acquisition was performed with a FluoView FV1000 confocal microscope (Olympus, Tokyo, Japan) equipped with Plapon 60X/1.42 and UPlanSapo 20X/0.75 objectives. Z stacks spanning the entire corneal epithelium were acquired at the corneal center (defined as the center of the nerve whorl or the center of the disorganized area in those samples with highly disrupted nerve whorls) and at two opposite locations 600 μm from the center, and analyzed at three different levels as previously described(3, 4). For subapical and mid-epithelial nerve analysis, the average of the three locations was used, whereas for subbasal nerves, only the central Z stack was quantified. For epithelial cell turnover analysis, a blind observer (AV) selected a single section encompassing the basal epithelial cells from one of the peripheral Z-stacks and manually counted the number of Ki-67$^{+}$ cells. Side views were obtained using the Orthogonal Views function of ImageJ.

**RNA isolation from trigeminal ganglia and RNA-Seq analysis.** The trigeminal ganglia were dissected after euthanasia and cardiac perfusion with PBS to remove contaminating blood cells(9), collected in ice-cold TRI Reagent, and stored at -80°C until processing. For RNA isolation, both trigeminal ganglia from one mouse were homogenized in 1 ml of TRI Reagent, and then 0.2 ml of isopropanol was added. After centrifugation, RNA was purified from the aqueous phase using the Direct-zol RNA MiniPrep kit (Cat #R2052, Zymo Research, Irvine, CA, USA) following the manufacturer's instructions. The concentration and purity of RNA were assessed with a NanoDrop 1000 spectrophotometer (ThermoFisher Scientific, Waltham, MA, USA). RNA-Seq was performed by NovoGene (Sacramento, CA, USA) using the Illumina NovaSeq platform to generate 150 bp paired-end reads. The sequenced reads were mapped to the mouse reference genome (assembly GRCm38/mm10) using STAR v2.7.11a, and the quantification of reads per gene was estimated by RSEM v1.3.1. Subsequent analyses were performed in R (v4.3.1). Gene filtering was performed by selecting features with >=10 reads in at least 3 samples. For quality controls, the filtered data were subjected to a principal component analysis and hierarchical clustering (Ward.D2 method, Euclidean distance) after trimmed mean of M-values normalization (edgeR v4.0.16). Gene information (ENSEMBL ID, external gene name and gene biotype) was extracted using biomaRt (v2.58.2). Differential gene expression analyses were performed by the DESeq2 methodology (v1.42.1). Genes with log2 fold change > 0.378 and adjusted p-value < 0.1 were considered as upregulated, and genes with log2 fold change < -0.378 and adjusted p-value < 0.1 were categorized as downregulated. The list of genes was ranked by log10 adjusted p-value * sign(log2 fold change) and subjected to a Gene Set Enrichment Analysis with the Gene Ontology database, using enrichGO and simplify (clusterProfiler v4.10.1). Enriched Gene Ontology terms with adjusted p-value of 0.05 were considered significant. All raw data files are available at ArrayExpress (https://www.ebi.ac.uk/biostudies/arrayexpress), accession E-MTAB-13945.

**RT-qPCR analysis of trigeminal gene expression.** Trigeminal RNA was extracted as for RNA-Seq and reverse transcription was performed as previously described(3). Real-time quantitative PCR (qPCR) was performed with 50 ng cDNA, SsoFast EvaGreen Supermix (Bio-Rad, USA), and primers in a final reaction volume of 20 µL. Primers (Table S2) were designed using the Primer3 software, purchased from Ruralex-Fagos (Buenos Aires, Argentina), and used at 400 nM. The reaction was performed in a CFX Connect Real-Time PCR Detection System (Bio-Rad, USA). The glyceraldehyde 3-phosphate dehydrogenase gene was used for normalization of the results for each sample and then the fold-change in specific mRNA levels between groups was calculated by using the $2^{-\Delta\Delta Ct}$ method.

**Tear cytokine levels.** To collect a tear washing sample, each mouse was manually restrained, then 5 µl of PBS+BSA 0.1% was applied on one eye using a 10-µl pipette tip, and finally the same volume was gently collected 20 seconds later from the ocular surface. The right and left eye washings from one mouse were pooled as one sample and stored at -80°C until processing. Cytokine levels were determined using a bead-based multiplex assay (LEGENDplex, #741048, BioLegend) following the manufacturer's instructions and are expressed as mean±SEM (pg/ml).

**Lacrimal gland histopathology.** Formalin-fixed extraorbital lacrimal gland specimens were embedded in paraffin, serially cut into 5-μm-thick sections, and stained with hematoxylin and eosin. At least four sections from each sample were examined to evaluate gland structure and inflammatory cell infiltration.

**Statistical analysis.** Student's t-test and one- or two-way analysis of variance (ANOVA) with Sidak's post hoc tests were used to compare the means of two or more samples, respectively. Significance was set at $p<0.05$ and two-tailed tests were used in all experiments. All data are shown as mean±standard error of measurement and each data point represents one animal (average of the two eyes). Calculations were performed using GraphPad Prism version 9 software (GraphPad Software, La Jolla, CA, USA).

**Figures S1 to S13**

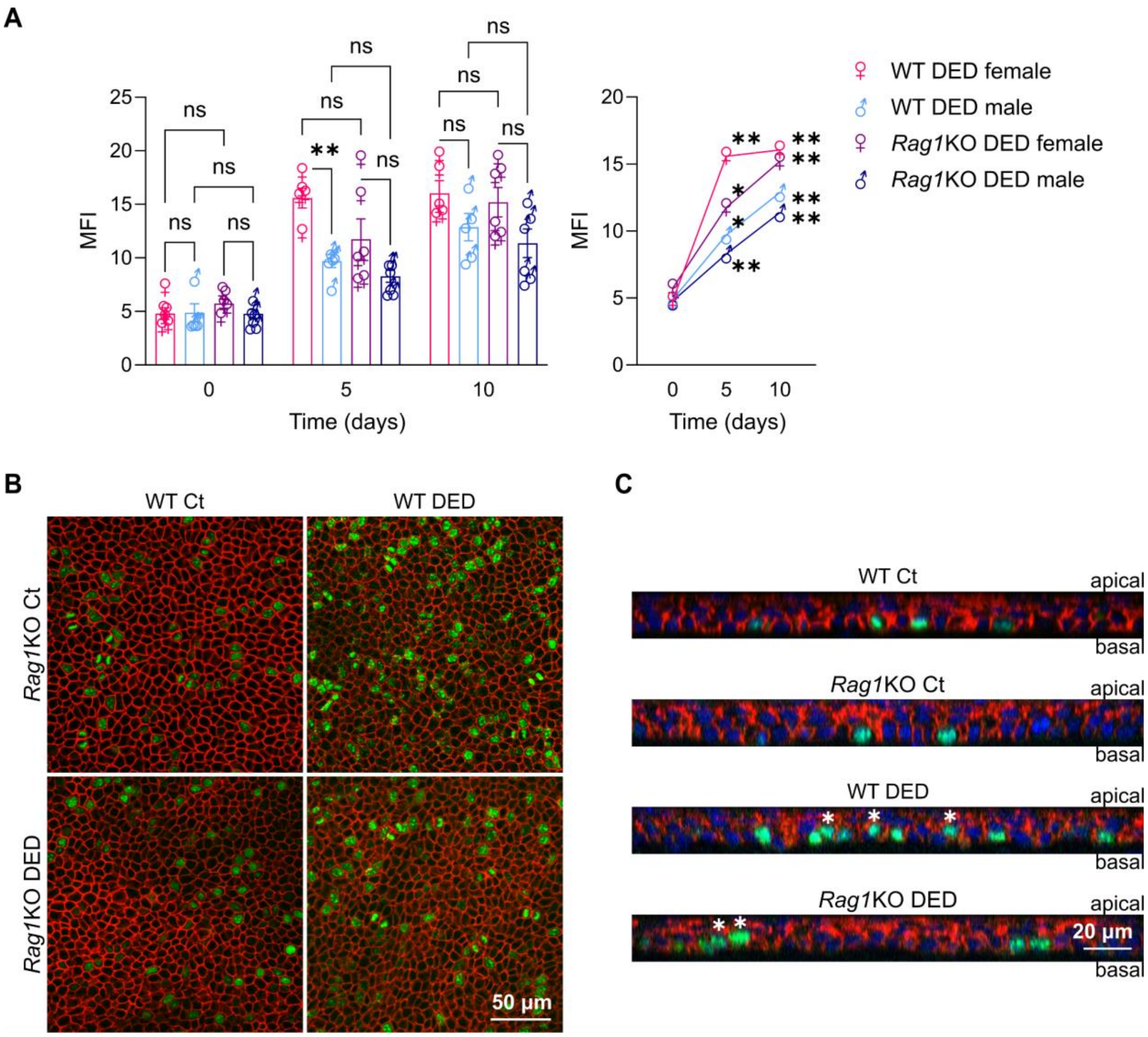


**Figure S1 - Effect of sex and epithelial cell turnover in dry eye-induced corneal epitheliopathy progression.** Dry eye disease (DED) was surgically induced in wild-type (WT) or recombination-activating gene 1-knockout (*Rag1*KO) mice of both sexes through bilateral excision of the extraorbital lacrimal gland. **A)** Corneal dextran-fluorescein uptake in both strains was quantified as mean fluorescence intensity (MFI) calculated with ImageJ software (Methods section). Individual data (left) and progression curves (right) from two independent experiments. **B)** Representative micrographs of proliferating cells within the basal layer and **C)** axial reconstructions of the entire epithelium of corneal whole-mounts obtained 10 days after DED induction and stained with Ki67 (green) and E-cadherin (red). Nuclei are shown in blue whereas suprabasal Ki67+ epithelial cells are marked with white asterisks in the axial reconstructions. To compare means, three-way ANOVA was used (strain, sex, and time) with Sidak's post hoc test. ♀ and ♂ indicate female and male, respectively; * indicates $p<0.05$, ** indicates $p<0.01$, and ns indicates not significant.

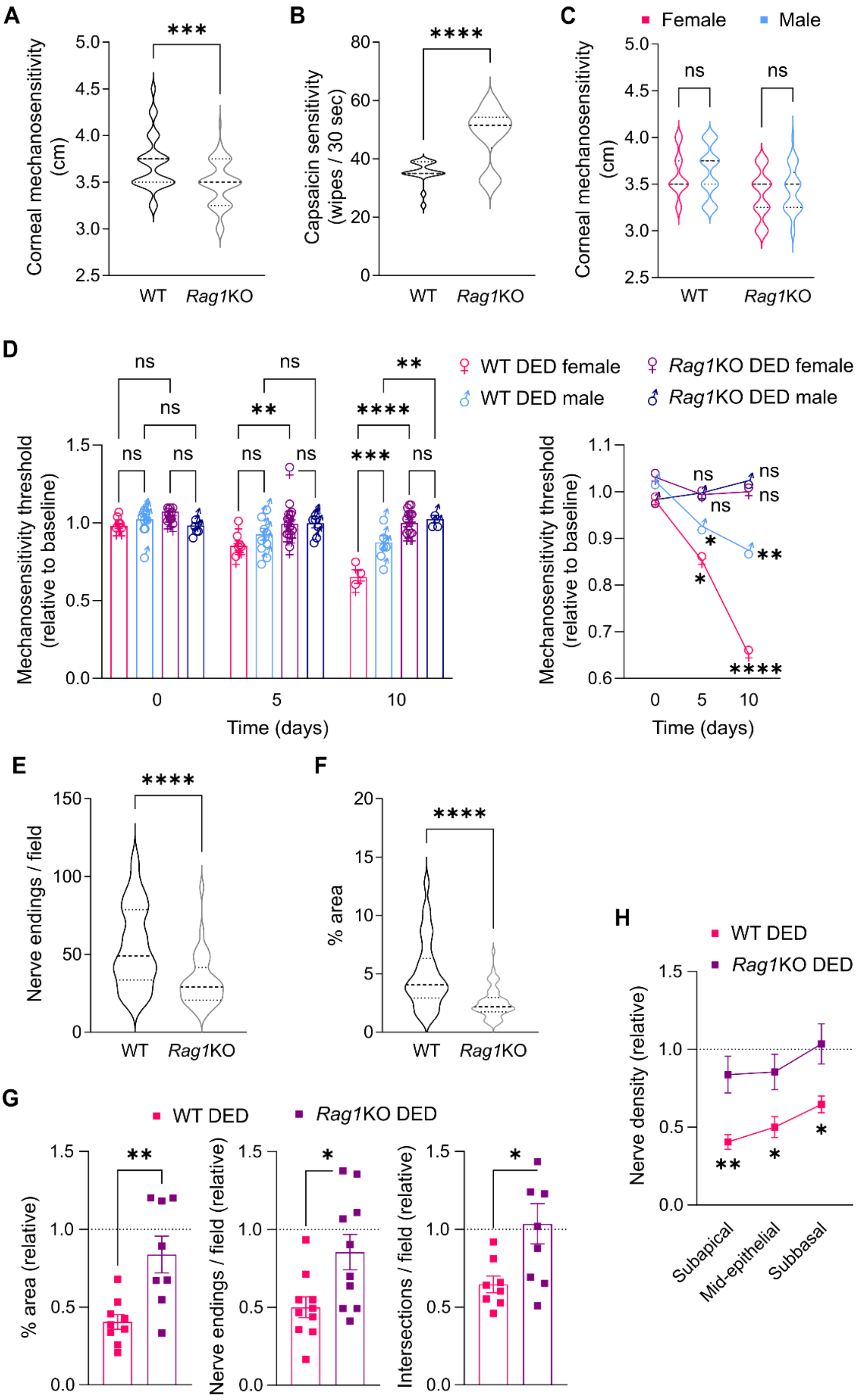


**Figure S2 - Strain-specific baseline differences and effect of sex and relative change in nerve density in dry eye-induced corneal neuropathy.** Dry eye disease (DED) was surgically induced in wild-type (WT) or recombination-activating gene 1-knockout (*Rag1*KO) mice of both sexes through bilateral excision of the extraorbital lacrimal gland. Corneal mechanosensitivity and capsaicin sensitivity were measured using nylon 6-0 filaments or 100 μM capsaicin instillation during the experiment whereas nerve morphology was assessed by confocal microscopy and quantified as described in Methods. **A)** Baseline mechanosensitivity thresholds (pooled data) from 39 WT and 56 *Rag1*KO mice. **B)** Baseline capsaicin sensitivity thresholds (pooled data) from 34 WT and 18 *Rag1*KO mice. **C)** Baseline mechanosensitivity thresholds (pooled data) aggregated by sex from 53 WT (22 female and 31 male) and 48 *Rag1*KO mice (19 female and 29 male). **D)** Mechanosensitivity thresholds in female and male DED mice from both strains on days 0, 5, and 10 of DED induction. Individual data (left) and progression curves (right) from two independent experiments. **E-F)** Pooled intraepithelial corneal nerve density data from sham-operated WT (n=21) and *Rag1*KO mice (n=19) analyzed at the subapical (E, % area occupied by nerve endings) and mid-epithelial (F, count of nerve endings/field) levels. **G)** Nerve density after 10 days of DED induction in mice from both strains (females and males combined) shown relative to sham-operated mice of the same strain as analyzed at three different levels of intraepithelial corneal innervation: subapical endings (left), mid-epithelial fibers (middle), and subbasal nerves (right). **H)** Distribution of change in nerve density in DED mice (day 10 after surgery, relative to sham-operated mice of the same strain, both sexes combined) over the three levels of intraepithelial corneal innervation. To compare means, three-way ANOVA was used (strain, sex, and time) with Sidak's post hoc test. ♀ and ♂ indicate female and male, respectively; * indicates $p<0.05$, ** indicates $p<0.01$, and ns indicates not significant.

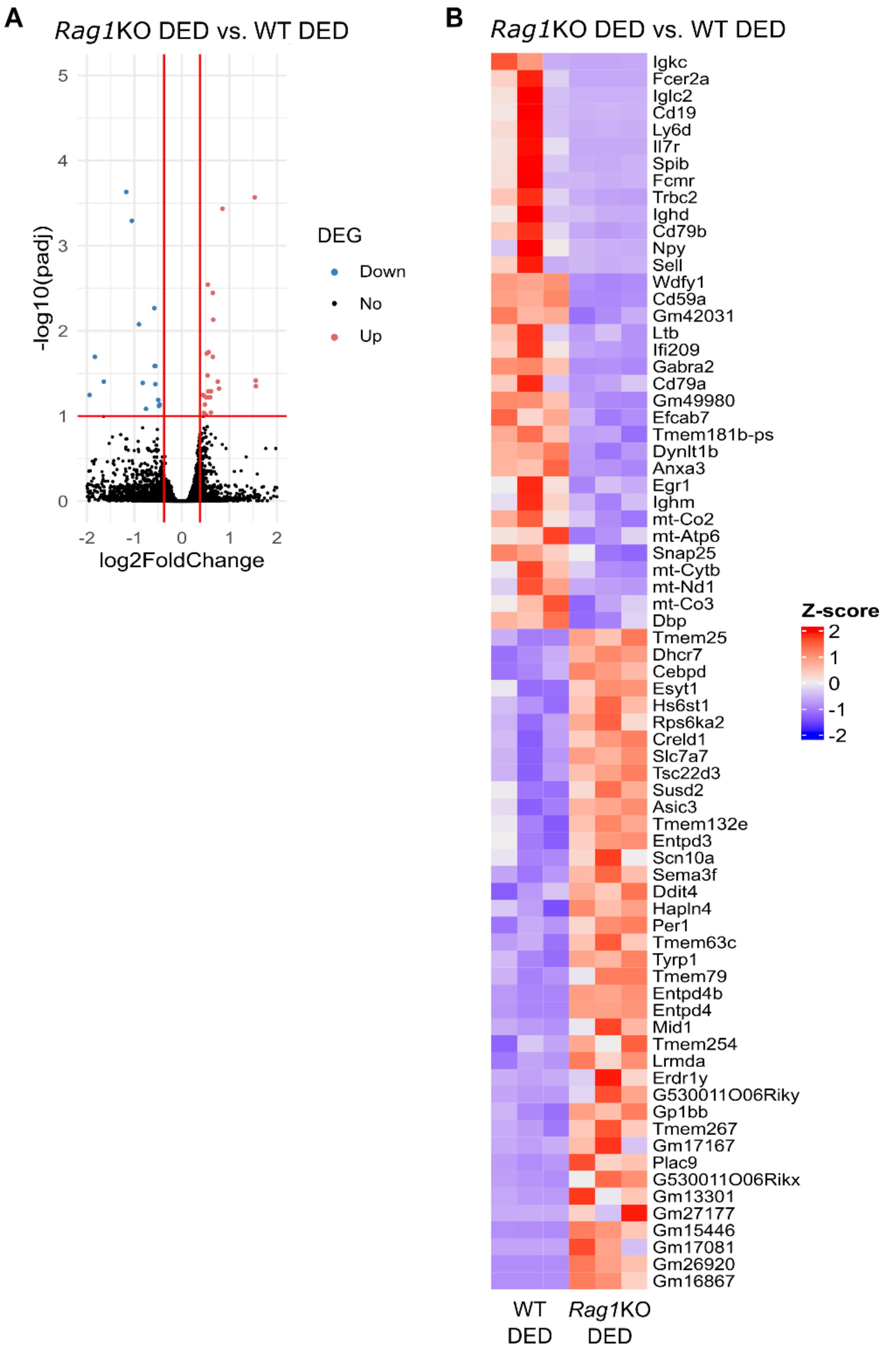


**Figure S3 - Comparison of dry eye-induced transcriptomic signatures in the trigeminal ganglion of wild-type and adaptive immune-deficient mice.** Dry eye disease (DED) was surgically induced in wild-type (WT) or recombination-activating gene 1-knockout (*Rag1*KO) mice for 10 days and then the trigeminal ganglia were harvested for bulk RNA-Seq analysis (female mice, n=3 per group). Differential gene expression was calculated between the sham-operated (Ct) and DED mice of each strain. **A)** Volcano plots of differentially expressed genes in DED *Rag1*KO vs WT mice. Up-regulated genes in DED *Rag1*KO mice are shown in red and down-regulated genes are shown in blue. **B)** Heatmaps (normalized counts, Z score) of the differentially expressed genes (fold change > 1.4, adjusted p-value < 0.1) in DED *Rag1*KO vs WT mice.

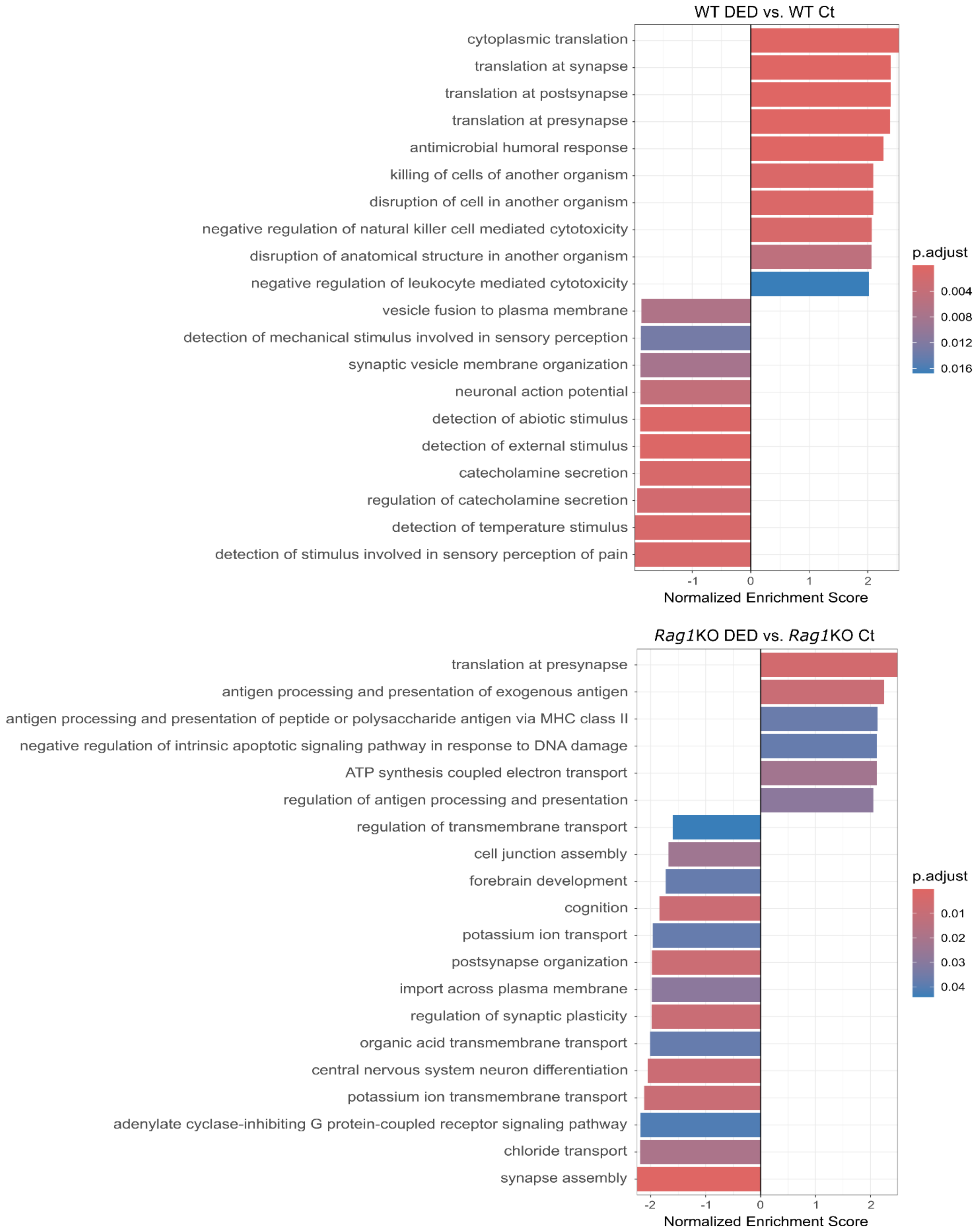


**Figure S4 - Biological pathways enriched in the dry eye-induced transcriptomic signatures in the trigeminal ganglion of wild-type and adaptive immune-deficient mice.** Dry eye disease (DED) was surgically induced in wild-type (WT) or recombination-activating gene 1-knockout (*Rag1*KO) mice for 10 days and then the trigeminal ganglia were harvested for bulk RNA-Seq analysis (female mice, n=3 per group). Differential gene expression was calculated between the sham-operated (Ct) and DED mice of each strain followed by gene set enrichment analysis (Gene Ontology). The 10 most significantly activated and suppressed pathways for each strain are shown.

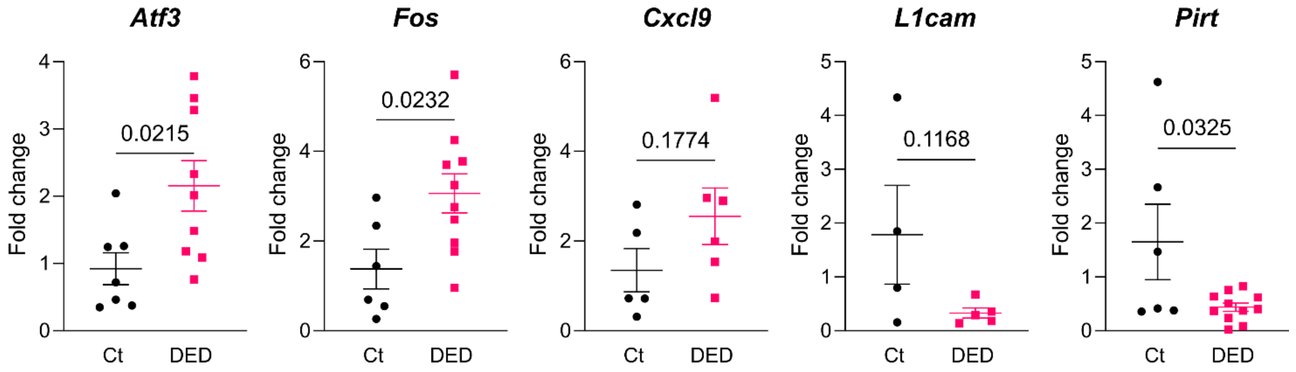


**Figure S5 - qPCR validation of trigeminal gene expression changes induced by dry eye.** Dry eye disease (DED) was surgically induced in wild-type (WT) mice for 10 days and then the trigeminal ganglia were harvested and assayed by qPCR for mRNA expression levels of five selected genes that were found to be differentially expressed by RNA-Seq. Sham-operated mice were included as controls (Ct). Mean±standard error of measurement is shown and Student's t test was applied to compare means.

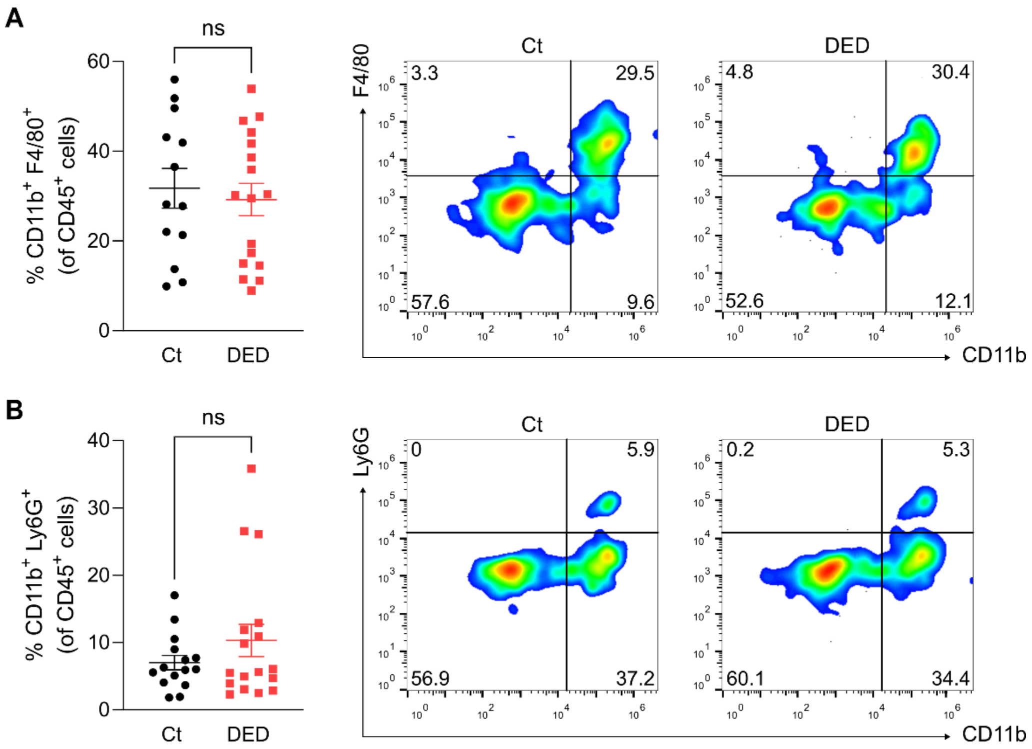


**Figure S6 – Conjunctival myeloid cells in dry eye mice.** Dry eye disease (DED) was surgically induced in wild-type (WT) mice for 10 days and then the conjunctiva was harvested, digested into a cell suspension and analyzed by flow cytometry for myeloid cells. Sham-operated mice were included as controls (Ct). The gating strategy was the same as in Figure S8. Cumulative data and representative dot plots for **A)** conjunctival macrophages, defined as $CD45^+$ $CD11b^+$ $F4/80^+$ $Ly6G^-$ cells, and **B)** neutrophils, defined as $CD45^+$ $CD11b^+$ $F4/80^-$ $Ly6G^+$ cells. The experiment was performed twice with 6 mice/group/experiment. Student's t test was used to compare means and ns indicates not significant.

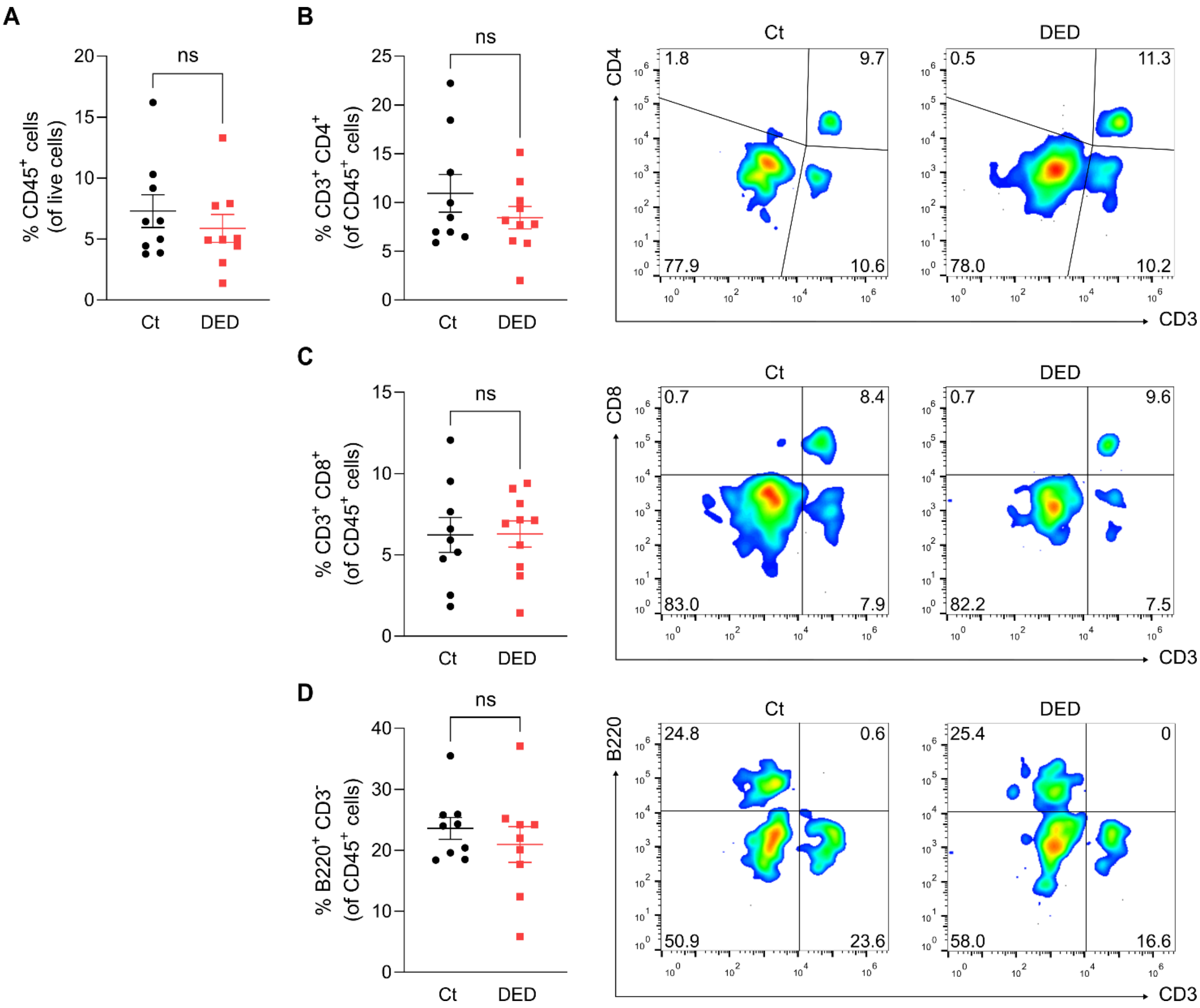


**Figure S7 – Adaptive immune cells in the trigeminal ganglia of dry eye mice.** Dry eye disease (DED) was surgically induced in wild-type (WT) mice for 10 days and then the trigeminal ganglia were harvested, digested into a cell suspension and analyzed by flow cytometry for T and B cells. Sham-operated mice were included as controls (Ct). The gating strategy was the same as in Figure S8. **A)** Number of trigeminal $CD45^+$ cells. Cumulative data and representative dot plots for **B)** $CD4^+$ T cells, defined as $CD45^+$ $CD3^+$ $CD4^+$ $CD8^-$ $B220^-$ cells, **C)** CD8+ T cells, defined as $CD45^+$ $CD3^+$ $CD4^-$ $CD8^+$ $B220^-$ cells, and **D)** B cells, defined as $CD45^+$ $CD3^+$ $CD4^-$ $CD8^-$ $B220^+$ cells. The experiment was performed twice with 4-5 mice/group/experiment. Student's t test was used to compare means and ns indicates not significant.

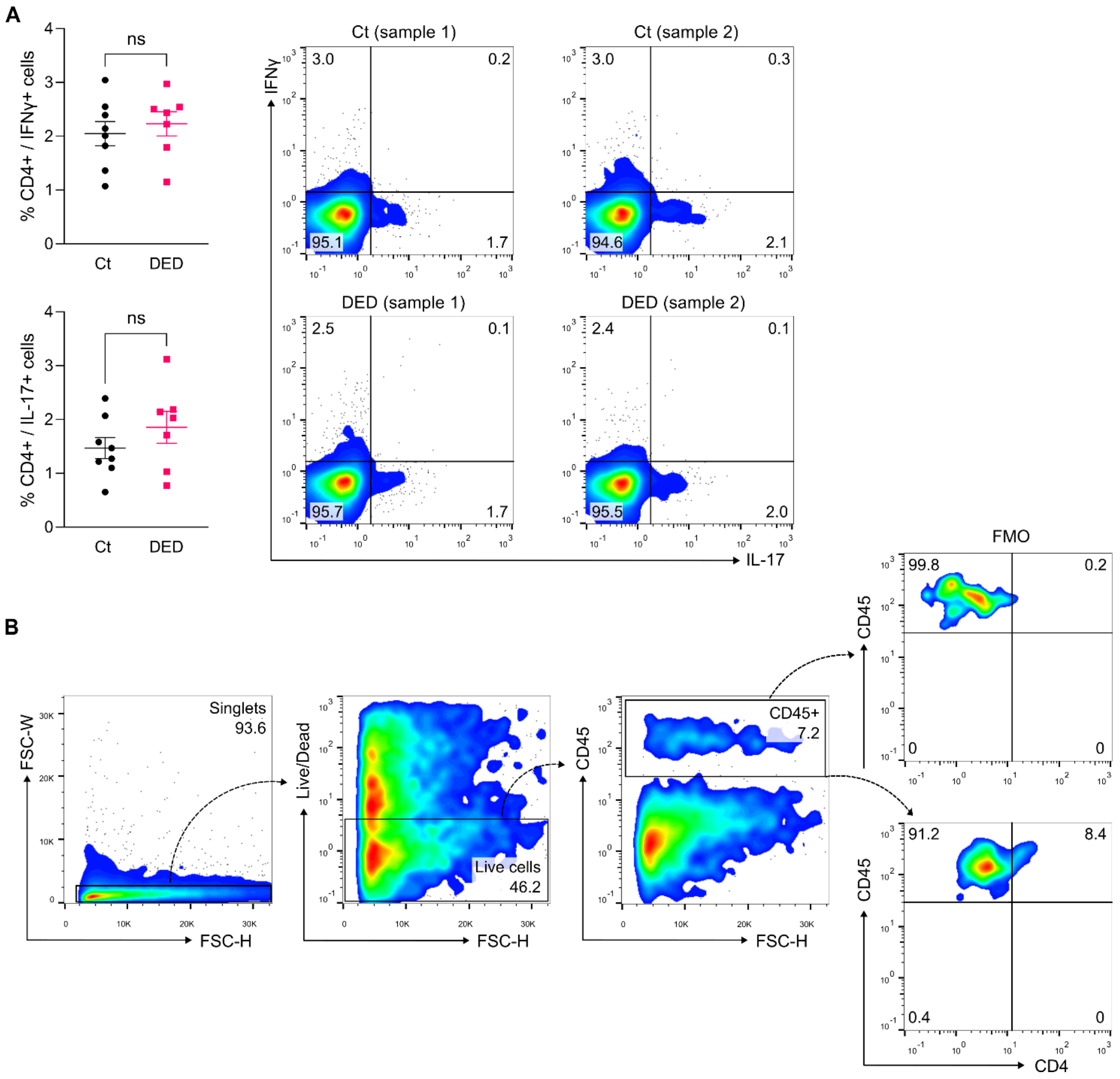


**Figure S8 - Adoptive transfer of dry eye-induced CD4$^+$ T cells reproduces corneal neuropathy but not epitheliopathy.** The experimental design was described in detail in Figure 5A and involved the adoptive transfer of control (Ct) or dry eye disease (DED)-induced CD4$^+$ T cells from WT mice into recombination-activating gene 1-knockout (*Rag1*KO) mice of the same sex that were subsequently evaluated over 4 weeks. **A)** Production of interferon (IFN)-γ and interleukin (IL)-17 by Ct or DED-induced CD4$^+$ T cells from WT mice after 10 days of induction as assessed by intracellular flow cytometry before the adoptive transfer into *Rag1*KO mice. Data from one experiment (left panel, n=7-8/group) and representative dot plots (right panel, 2 samples from each group). **B)** Gating strategy with representative example of flow cytometry analysis of conjunctival CD4$^+$ T cells in *Rag1*KO mice 4 weeks after adoptive transfer. Student's t test was applied to compared means in A. FSC-H indicates forward side height, FSC-W indicates forward scatter width, FMO indicates fluorescence-minus-one control, and ns indicates not significant.

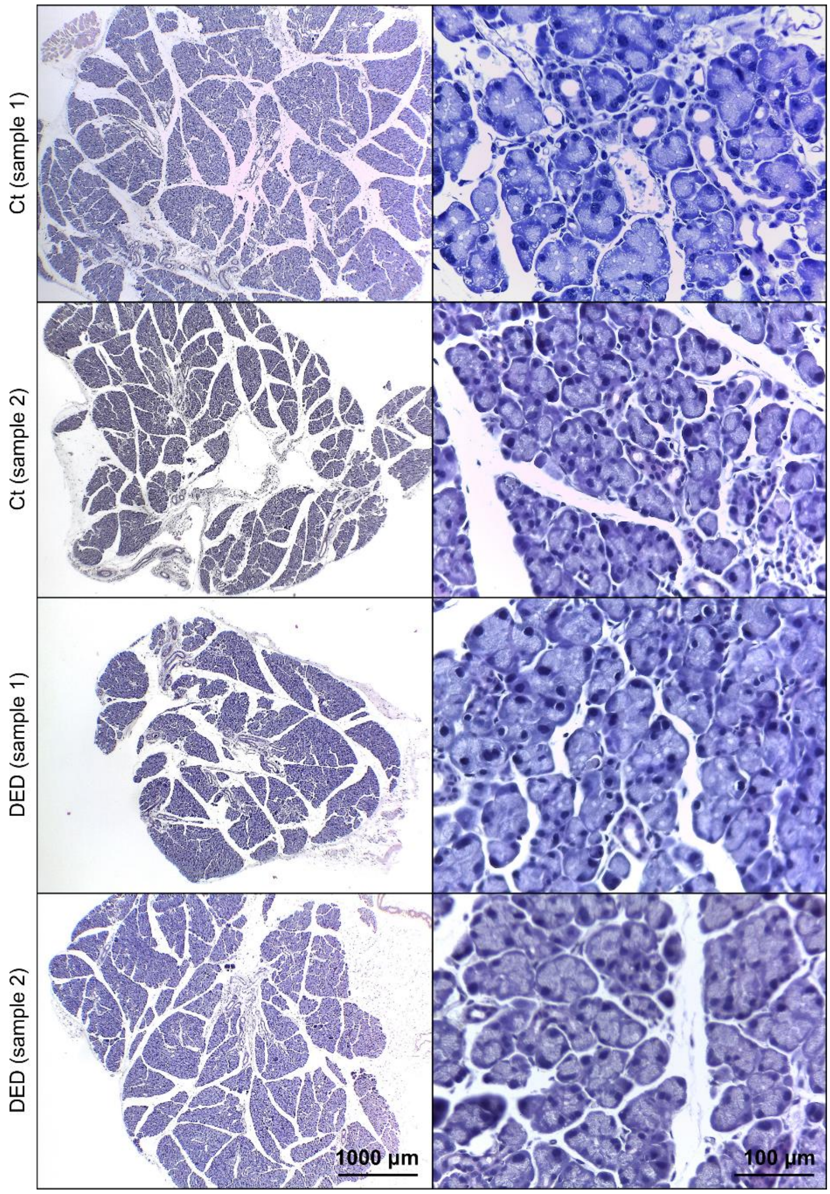


**Figure S9 – Lacrimal gland histopathology after adoptive transfer of dry eye-induced CD4$^+$ T cells.** The experimental design was described in detail in Figure 5A and involved the adoptive transfer of control (Ct) or dry eye disease (DED)-induced CD4$^+$ T cells from WT mice into same-sex recombination-activating gene 1-knockout (*Rag1*KO) mice. Low- (left) and high-magnification (right) representative histopathology micrographs of extraorbital lacrimal glands harvested 4 weeks after adoptive transfer (2 samples from each group, hematoxylin-eosin stain).

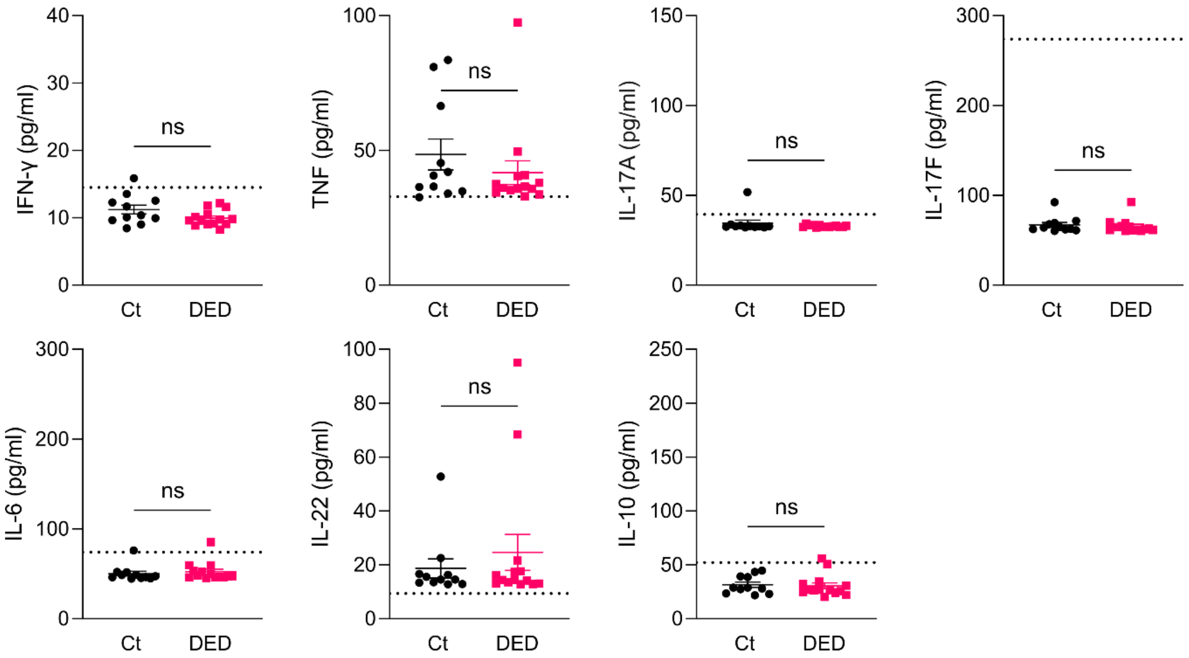


**Figure S10 - Cytokine levels in tear washings after adoptive transfer of dry eye-induced CD4$^+$ T cells.** Tear washings were collected 4 weeks after adoptive transfer of control (Ct) or dry eye disease (DED)-induced CD4$^+$ T cells from WT mice into same-sex recombination-activating gene 1-knockout (*Rag1*KO) mice, as described in Figure 5A. Samples were analyzed by a bead-based multiplex assay for interferon (IFN)-γ, tumor necrosis factor (TNF), interleukin (IL)-6, -10, -17A, -17F, and -22 levels. Cumulative data from two experiments with 6 mice/group are shown in pg/ml (mean±SEM). The dashed line in each graph represents the mean cytokine concentration previously found in sham-operated *Rag1*KO mice (Figure 4). Student's t test was used to compare means and ns indicates not significant.

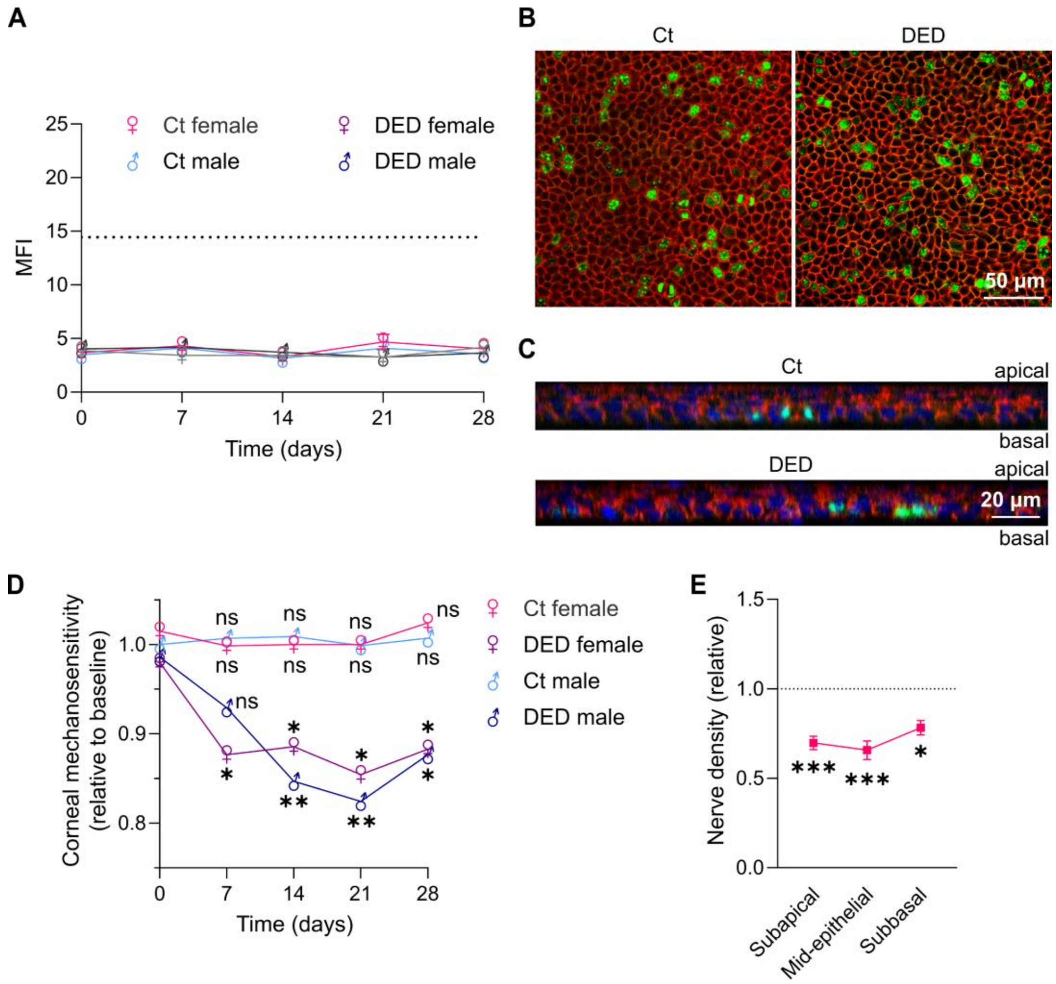


**Figure S11 - Adoptive transfer of dry eye-induced CD4⁺ T cells reproduces corneal neuropathy but not epitheliopathy.** The experimental design was described in detail in Figure 5A and involved the adoptive transfer of control (Ct) or dry eye disease (DED)-induced $CD4^+$ T cells from WT mice into recombination-activating gene 1-knockout (*Rag1*KO) mice of the same sex that were subsequently evaluated over 4 weeks. **A)** Sex-aggregated data of corneal dextran-fluorescein uptake in Ct- and DED-induced $CD4^+$ T cell recipients. Data shown as the mean fluorescence intensity (MFI) calculated with ImageJ software (Methods section). **B)** Representative micrographs of proliferating cells within the basal layer and **C)** axial reconstructions of the entire epithelium of corneal whole-mounts obtained 4 weeks after adoptive transfer induction and stained with Ki67 (green) and E-cadherin (red). Nuclei are shown in blue in the axial reconstructions. **D)** Corneal mechanosensitivity thresholds in Ct- and DED-induced $CD4^+$ T cell recipients aggregated by sex. **E)** Distribution of change in nerve density in DED-induced $CD4^+$ T cell-recipients (both sexes combined) over the three levels of intraepithelial corneal innervation (quantified as for Figure 6 and expressed relative to the Ct $CD4^+$ T cell recipients that are represented by the dotted line). All experiments were performed twice or more with 6 mice/group/experiment. To compare means, three-way ANOVA was used (strain, sex, and time) with Sidak's post hoc test was used for A and D, and Student's t test was applied in E. ♀ and ♂ indicate female and male, respectively; * indicates p<0.05, ** indicates p<0.01, *** indicates p<0.001, and ns indicates not significant.

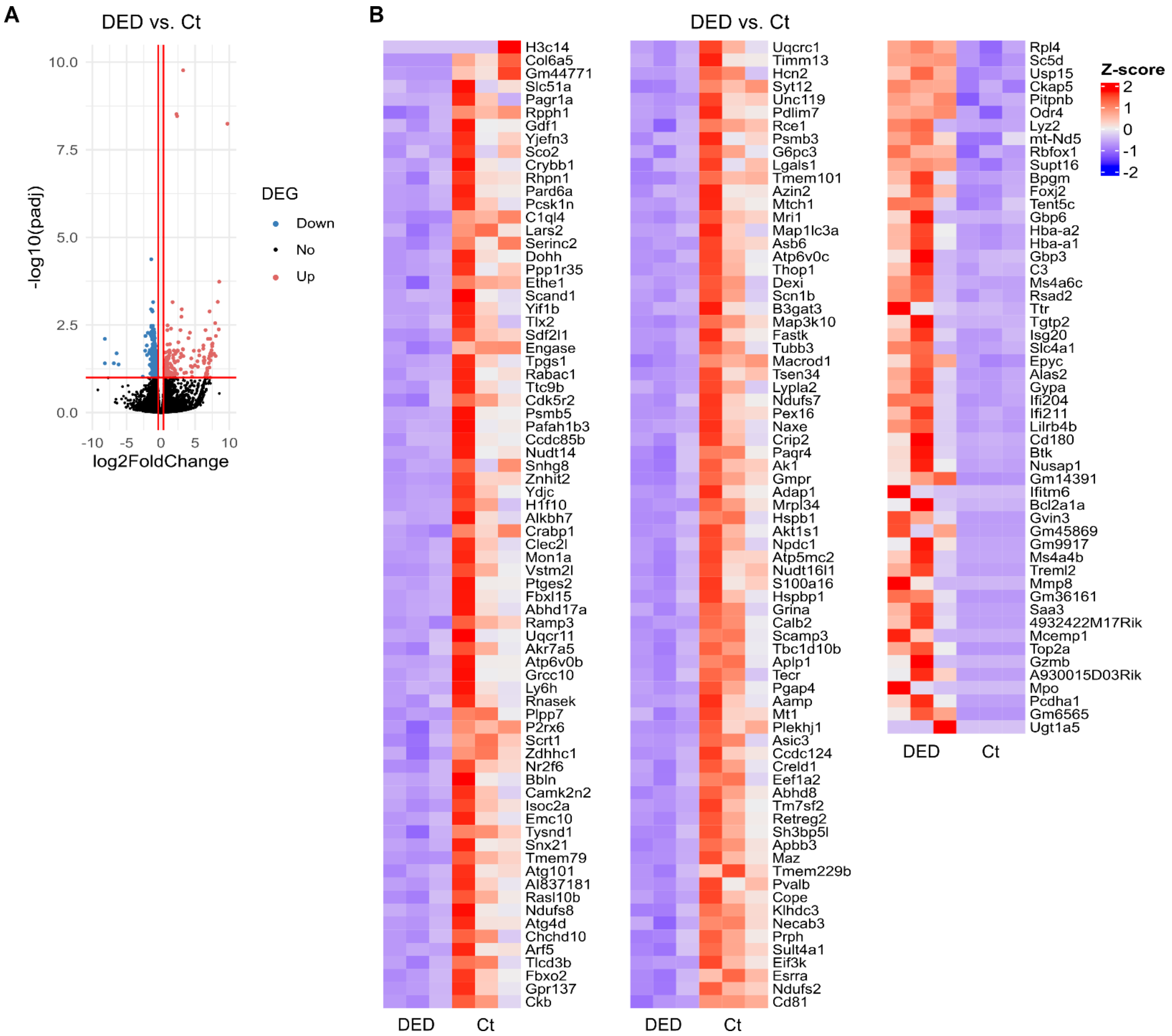


**Figure S12 - Trigeminal gene expression changes induced by adoptive transfer of dry eye-induced CD4⁺ T cells.** CD4⁺ T cells were isolated from the spleens and lymph nodes of female wild-type mice 10 days after surgical induction of dry eye disease (DED) and then adoptively transferred into sex-matched recombination-activating gene 1-knockout (*Rag1*KO) mice. Sham-operated WT mice were used as a source of control (Ct) CD4⁺ T cells. After 4 weeks, the trigeminal ganglia of the recipient *Rag1*KO mice were harvested for bulk RNA-Seq analysis (n=3 per group). Differential gene expression was calculated between the mice receiving Ct and DED-induced CD4⁺ T cells. **A)** Volcano plots of differentially expressed genes. Up-regulated genes in DED-induced CD4⁺ T cell-recipient mice are shown in red and down-regulated genes are shown in blue. **B)** Heatmaps (normalized counts, Z score) of the differentially expressed genes (fold change > 1.4, adjusted p-value < 0.1).

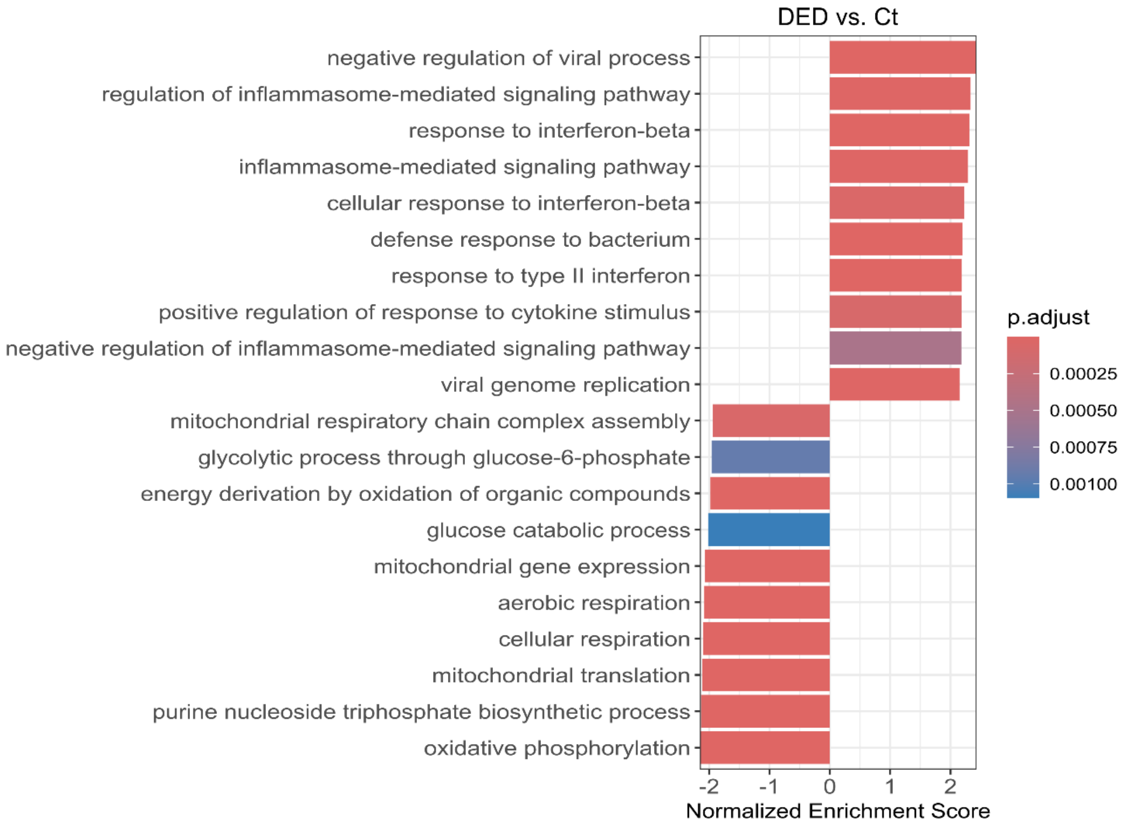


**Figure S13 - Biological pathways enriched in the trigeminal transcriptomic signatures after adoptive transfer of dry eye-induced CD4$^+$ T cells.** CD4$^+$ T cells were isolated from the spleens and lymph nodes of female wild-type mice 10 days after surgical induction of dry eye disease (DED) and then adoptively transferred into sex-matched recombination-activating gene 1-knockout (*Rag1*KO) mice. Sham-operated WT mice were used as a source of control (Ct) CD4$^+$ T cells. After 4 weeks, the trigeminal ganglia of the recipient *Rag1*KO mice were harvested for bulk RNA-Seq analysis (n=3 per group). Differential gene expression was calculated between the mice receiving Ct and DED-induced CD4$^+$ T cells followed by gene set enrichment analysis (Gene Ontology). The 10 most significantly activated and suppressed pathways for each strain are shown.

**Tables S1 to S2**

### Table S1 – Reagents and antibodies

| Reagent/Antibody | Product # | Concentration/dilution | Supplier |
|---|---|---|---|
| Dextran-fluorescein isothiocyanate | FD4-100MG | 10 mg/ml in PBS | Sigma-Aldrich (Buenos Aires, Argentina) |
| Ketamine | Ketonal 50 | 100 mg/kg | Richmond Vet (Grand Bourg, Argentina) |
| Xylazine | Xilacina 20 | 10 mg/kg | Richmond Vet (Grand Bourg, Argentina) |
| Alexa Fluor® 488 anti-tubulin β3 | 801203 | 2.5-3.5 µg/ml | BioLegend (San Diego, CA, USA) |
| Alexa Fluor® 594 anti-mouse/human Ki-67 | 151213 | 0.5 µl/100 µl | BioLegend (San Diego, CA, USA) |
| Alexa Fluor® 647 anti-mouse/human CD324 (E-cadherin) | 147308 | 2.5 µg/ml | BioLegend (San Diego, CA, USA) |
| Wheat Germ Agglutinin (WGA) Alexa Fluor® 647 Conjugate | W32466 | 10 µg/ml | ThermoFisher (Buenos Aires, Argentina) |
| Live/Dead Fixable Dead Cell Stain | L10119 | 1:500 in PBS | ThermoFisher (Buenos Aires, Argentina) |
| CD45 APC | 103112 | 0.5 µl/100 µl | BioLegend (San Diego, CA, USA) |
| CD4 FITC | 100406 | 0.5 µl/100 µl | BioLegend (San Diego, CA, USA) |
| CD8 Pacific Blue | 100725 | 0.5 µl/100 µl | BioLegend (San Diego, CA, USA) |
| CD3 PE | 100206 | 0.5 µl/100 µl | BioLegend (San Diego, CA, USA) |
| B220 PE-Cy7 | 103210 | 0.5 µl/100 µl | BioLegend (San Diego, CA, USA) |
| F4/80 PE-Cy7 | 123114 | 0.5 µl/100 µl | BioLegend (San Diego, CA, USA) |
| Ly6G FITC | 127606 | 0.5 µl/100 µl | BioLegend (San Diego, CA, USA) |
| Fc Block | 158002 | 0.5 µl/100 µl | BioLegend (San Diego, CA, USA) |

**Table S2 - qPCR primers**

| Gene name | Gene symbol | Forward primer 5' > 3' | Reverse Primer 5' >3' | Predicted size (bp) |
|---|---|---|---|---|
| Glyceraldehyde 3-phosphate dehydrogenase | *Gadph* | CTCCCACTCTTCCACCTTCG | CCACCACCCTGTTGCTGTAG | 110 |
| Fos proto-oncogene, AP-1 transcription factor subunit | *Fos* | GGGAATGGTGAAGACCGTGTCA | GCAGCCATCTTATTCCGTTCCC | 126 |
| L1 cell adhesion molecule | *L1cam* | AGTTCCGCTGGACGAAAGATG | CGATAGATGCCCTGAAACCTCT | 139 |
| Phosphoinositide-Interacting Regulator of Transient receptor potential channels | *Pirt* | GCAAGTGCTCAGGATGATAGGG | GGAGGAAGAACTTGAGGCTTTGG | 146 |
| Activating transcription factor 3 | *Atf3* | GAAGATGAGAGGAAAAGGAGGCG | GCTCAGCATTCACACTCTCCAG | 120 |
| Chemokine (C-X-C motif) ligand 9 | *Cxcl9* | CCTAGTGATAAGGAATGCACGATG | CTAGGCAGGTTTGATCTCCGTTC | 158 |

**Legends for Datasets S1 to S2**

**Dataset S1** (separate file). List of differentially expressed genes described in Figures 3, S3, and S4.

**Dataset S2** (separate file). List of differentially expressed genes described in Figures S12 and S13.

**Supporting Information References**
