## Supplementary material for "CD4^+^ T cells drive corneal nerve damage but not epitheliopathy in an acute aqueous-deficient dry eye model": Dataset S1

DEGs WT DED vs WT Ct

| baseMean | log2FoldChange | lfcSE | stat | pvalue | padj | DE | gene_id | external_gene_name |
|---|---|---|---|---|---|---|---|---|
| 63.24515692 | 6.022567907 | 0.79446937 | 7.580616871 | 3.43916E-14 | 5.5704E-10 | Up | ENSMUSG00000050359.7 | Sprr1a |
| 2042.412329 | 0.833785908 | 0.138176077 | 6.034227688 | 1.59725E-09 | 1.29353E-05 | Up | ENSMUSG00000026185.8 | Igfbp5 |
| 1630.138853 | 0.698929193 | 0.122733091 | 5.694708637 | 1.23583E-08 | 6.67223E-05 | Up | ENSMUSG00000037206.15 | Islr |
| 295.9440323 | 1.089629283 | 0.197438104 | 5.518839886 | 3.41245E-08 | 0.000138179 | Up | ENSMUSG00000050010.8 | Shisa3 |
| 665.4800013 | 0.662051563 | 0.121902008 | 5.4310144 | 5.60346E-08 | 0.000181519 | Up | ENSMUSG00000001119.7 | Col6a1 |
| 6950.687647 | 0.49269047 | 0.096958783 | 5.081442412 | 3.7458E-07 | 0.001011178 | Up | ENSMUSG00000007097.14 | Atp1a2 |
| 365.9217621 | 0.821380602 | 0.163923286 | 5.010762187 | 5.42149E-07 | 0.001254455 | Up | ENSMUSG00000046402.10 | Rbp1 |
| 16.04099591 | 19.03594086 | 3.909601721 | 4.86902304 | 1.12151E-06 | 0.002270644 | Up | ENSMUSG00000066632.3 | Pgk1-rs7 |
| 164.4725483 | 1.29948547 | 0.284112155 | 4.573846796 | 4.7885E-06 | 0.005539953 | Up | ENSMUSG00000019890.4 | Nts |
| 923.2564482 | 0.766323952 | 0.169633615 | 4.517524143 | 6.25669E-06 | 0.006755973 | Up | ENSMUSG00000042700.16 | Sipa1l1 |
| 201.6322358 | 1.085465053 | 0.250403312 | 4.334866995 | 1.45848E-05 | 0.013123914 | Up | ENSMUSG00000061167.9 | Rpl15-ps3 |
| 250.2697775 | 0.848263148 | 0.198310143 | 4.277457194 | 1.8904E-05 | 0.014562002 | Up | ENSMUSG00000021250.13 | Fos |
| 931.2515491 | 0.495044105 | 0.116245262 | 4.258617479 | 2.05695E-05 | 0.014562002 | Up | ENSMUSG00000026255.15 | Efhd1 |
| 40.08142563 | 2.950644259 | 0.693056269 | 4.257438237 | 2.06783E-05 | 0.014562002 | Up | ENSMUSG00000029819.6 | Npy |
| 1457.627648 | 0.686379997 | 0.161633515 | 4.246520269 | 2.17116E-05 | 0.014652626 | Up | ENSMUSG00000020821.17 | Kif1c |
| 411.3786911 | 0.967943414 | 0.229487176 | 4.217854053 | 2.46638E-05 | 0.015979207 | Up | ENSMUSG00000079017.3 | Ifi27l2a |
| 917.0029595 | 0.760053762 | 0.18322287 | 4.148247224 | 3.3503E-05 | 0.018712031 | Up | ENSMUSG00000031740.8 | Mmp2 |
| 98752.54939 | 0.553503828 | 0.134112067 | 4.12717394 | 3.67249E-05 | 0.019827753 | Up | ENSMUSG00000041607.17 | Mbp |
| 371.8271379 | 0.625544188 | 0.152522095 | 4.101334887 | 4.10774E-05 | 0.021462256 | Up | ENSMUSG00000022469.17 | Rapgef3 |
| 151.1641099 | 1.119334859 | 0.280127423 | 3.99580607 | 6.44745E-05 | 0.027481406 | Up | ENSMUSG00000006369.14 | Fbln1 |
| 193.4913916 | 0.944714821 | 0.236180549 | 3.99996877 | 6.33508E-05 | 0.027481406 | Up | ENSMUSG00000026628.13 | Atf3 |
| 220.4414475 | 0.818342167 | 0.206354308 | 3.965713995 | 7.31766E-05 | 0.030390791 | Up | ENSMUSG00000028108.15 | Ecm1 |
| 1285.943831 | 0.463732116 | 0.119132343 | 3.892579507 | 9.9184E-05 | 0.0334648 | Up | ENSMUSG00000022587.14 | Ly6e |
| 1765.708525 | 0.466003574 | 0.120796071 | 3.857770969 | 0.000114426 | 0.0334648 | Up | ENSMUSG00000027333.18 | Smox |
| 1531.865768 | 0.510205638 | 0.131720131 | 3.87340669 | 0.000107325 | 0.0334648 | Up | ENSMUSG00000078974.10 | Sec61g |
| 20.82182591 | 4.669249539 | 1.205656853 | 3.872784804 | 0.000107599 | 0.0334648 | Up | ENSMUSG00000104597.1 | B230334C09Rik |
| 410.0040518 | 0.574446871 | 0.150258555 | 3.823056004 | 0.000131808 | 0.034263656 | Up | ENSMUSG00000009614.16 | Sardh |
| 1136.766625 | 0.432135374 | 0.112810822 | 3.830619837 | 0.000127821 | 0.034263656 | Up | ENSMUSG00000020889.11 | Nr1d1 |
| 417.8934686 | 0.644499789 | 0.168702566 | 3.820331864 | 0.000133272 | 0.034263656 | Up | ENSMUSG00000028927.6 | Padi2 |
| 3070.093662 | 0.578646493 | 0.152145684 | 3.803239605 | 0.000142816 | 0.035952349 | Up | ENSMUSG00000013523.13 | Bcas1 |
| 27.72215531 | 3.476597223 | 0.914722224 | 3.800713628 | 0.00014428 | 0.035952349 | Up | ENSMUSG00000050505.7 | Pcdh20 |
| 181.5506327 | 0.888293935 | 0.236035805 | 3.763386391 | 0.000167628 | 0.040523419 | Up | ENSMUSG00000060550.16 | H2-Q7 |
| 565.9656142 | 0.992912072 | 0.26429427 | 3.756842974 | 0.00017207 | 0.040985649 | Up | ENSMUSG00000048583.16 | Igf2 |
| 1872.781501 | 0.479043975 | 0.129832405 | 3.689710401 | 0.000224509 | 0.049813426 | Up | ENSMUSG00000059824.12 | Dbp |
| 414.7363249 | 0.596566229 | 0.162170428 | 3.678637581 | 0.000234483 | 0.051323301 | Up | ENSMUSG00000008999.7 | Bmp7 |
| 667.2188558 | 0.519001736 | 0.141788774 | 3.660386647 | 0.000251835 | 0.053294892 | Up | ENSMUSG00000020241.13 | Col6a2 |
| 427.3723825 | 0.692753513 | 0.190791127 | 3.630952456 | 0.000282377 | 0.055546025 | Up | ENSMUSG00000035929.11 | H2-Q4 |
| 196.3838757 | 0.841423879 | 0.232977526 | 3.611609637 | 0.000304302 | 0.05665271 | Up | ENSMUSG00000026276.20 | Septin2 |
| 368.4057275 | 0.791582076 | 0.219041028 | 3.613853004 | 0.00030168 | 0.05665271 | Up | ENSMUSG00000050335.17 | Lgals3 |
| 1140.973333 | 0.410010991 | 0.114591617 | 3.578019063 | 0.000346208 | 0.060296068 | Up | ENSMUSG00000036256.13 | Igfbp7 |
| 576.621154 | 0.593265912 | 0.166124027 | 3.571222792 | 0.000355318 | 0.061224399 | Up | ENSMUSG00000001911.16 | Nfix |
| 71.02192722 | 1.100129596 | 0.308544575 | 3.56554509 | 0.000363101 | 0.061261901 | Up | ENSMUSG00000022440.9 | C1qtnf6 |
| 445.2705391 | 0.758106825 | 0.212460373 | 3.568226941 | 0.000359405 | 0.061261901 | Up | ENSMUSG00000046718.8 | Bst2 |
| 135.3737106 | 1.122172835 | 0.315924052 | 3.552033563 | 0.000382266 | 0.06254105 | Up | ENSMUSG00000040152.8 | Thbs1 |
| 2463.207292 | 0.446678432 | 0.126133039 | 3.541327771 | 0.000398119 | 0.064483282 | Up | ENSMUSG00000031765.8 | Mt1 |
| 451.6398365 | 0.536814478 | 0.152093122 | 3.529511869 | 0.000416327 | 0.066110283 | Up | ENSMUSG00000024812.11 | Tjp2 |
| 32.54918966 | 2.649702425 | 0.756873569 | 3.500852102 | 0.000463773 | 0.069553089 | Up | ENSMUSG00000029417.9 | Cxcl9 |
| 17.12415297 | 2.634348306 | 0.758968143 | 3.470960314 | 0.000518601 | 0.071130127 | Up | ENSMUSG00000026247.13 | Ecel1 |
| 628.4937193 | 0.525825972 | 0.151582958 | 3.468899009 | 0.000522596 | 0.071130127 | Up | ENSMUSG00000047797.14 | Gjb1 |
| 784.8345033 | 0.443160796 | 0.12765275 | 3.471611828 | 0.000517344 | 0.071130127 | Up | ENSMUSG00000073418.4 | C4b |
| 2027.125582 | 0.51051423 | 0.147718765 | 3.455987659 | 0.00054828 | 0.07304563 | Up | ENSMUSG00000026043.18 | Col3a1 |
| 641.0101043 | 0.481497671 | 0.139383217 | 3.454488146 | 0.000551338 | 0.07304563 | Up | ENSMUSG00000041216.15 | Clvs1 |
| 301.7679007 | 0.644865391 | 0.186763513 | 3.452844613 | 0.000554708 | 0.07304563 | Up | ENSMUSG00000079235.10 | Ccdc13 |
| 249.0849728 | 0.75790721 | 0.220428176 | 3.438340887 | 0.00058529 | 0.074645263 | Up | ENSMUSG00000016024.9 | Lbp |
| 404.0697884 | 0.715567768 | 0.208313414 | 3.435053719 | 0.000592436 | 0.074966344 | Up | ENSMUSG00000018920.11 | Cxcl16 |
| 2571.005628 | 0.572100314 | 0.166755302 | 3.430777355 | 0.000601854 | 0.074986423 | Up | ENSMUSG00000001506.10 | Col1a1 |
| 590.6855023 | 0.45428744 | 0.13234367 | 3.432634434 | 0.000597747 | 0.074986423 | Up | ENSMUSG00000002059.18 | Rab34 |
| 564.3952642 | 0.511929087 | 0.14953358 | 3.423505846 | 0.000618189 | 0.075402536 | Up | ENSMUSG00000040488.18 | Ltbp4 |
| 446.2094505 | 0.527616899 | 0.15448989 | 3.415219589 | 0.000637306 | 0.076711783 | Up | ENSMUSG00000040003.18 | Magi2 |
| 424.5194458 | 0.659387178 | 0.195268709 | 3.376819471 | 0.000733292 | 0.078656452 | Up | ENSMUSG00000021702.7 | Thbs4 |
| 3969.52041 | 0.505020336 | 0.149547446 | 3.376990711 | 0.000732835 | 0.078656452 | Up | ENSMUSG00000029661.16 | Col1a2 |
| 1396.581485 | 0.486562203 | 0.143471996 | 3.391339185 | 0.00069552 | 0.078656452 | Up | ENSMUSG00000031375.17 | Bgn |
| 392.5742703 | 0.503511046 | 0.149075275 | 3.377562419 | 0.000731314 | 0.078656452 | Up | ENSMUSG00000034462.9 | Pkd2 |
| 439.7490323 | 0.861230916 | 0.255028496 | 3.376998759 | 0.000732814 | 0.078656452 | Up | ENSMUSG00000049907.8 | Rasl11b |
| 2673.269669 | 0.624498105 | 0.184199997 | 3.390326365 | 0.000698095 | 0.078656452 | Up | ENSMUSG00000060586.11 | H2-Eb1 |
| 43766.40754 | 0.475150405 | 0.140080528 | 3.391980385 | 0.000693894 | 0.078656452 | Up | ENSMUSG00000064357.1 | mt-Atp6 |

| | | | | | | | | |
|---|---|---|---|---|---|---|---|---|
| 269.9461742 | 0.596219135 | 0.17621546 | 3.383466672 | 0.000715769 | 0.078656452 | Up | ENSMUSG00000100755.1 | Rps23-ps1 |
| 5815.019039 | 0.421524393 | 0.125070866 | 3.370284444 | 0.000750906 | 0.079753121 | Up | ENSMUSG00000028639.14 | Ybx1 |
| 2481.845477 | 0.641142744 | 0.190627407 | 3.363329295 | 0.000770084 | 0.080471342 | Up | ENSMUSG00000036594.15 | H2-Aa |
| 8386.427843 | 0.536057589 | 0.159491425 | 3.361043328 | 0.000776486 | 0.080620198 | Up | ENSMUSG00000040428.19 | Plekha4 |
| 6581.158849 | 0.569476305 | 0.170340869 | 3.34315722 | 0.00082831 | 0.082815618 | Up | ENSMUSG00000024610.15 | Cd74 |
| 228.6193777 | 0.602797499 | 0.180553648 | 3.338606034 | 0.000841999 | 0.083667827 | Up | ENSMUSG00000036545.9 | Adamts2 |
| 2065.980848 | 0.587946615 | 0.177119298 | 3.319494946 | 0.000901804 | 0.085897143 | Up | ENSMUSG00000023046.6 | Igfbp6 |
| 82.00383828 | 1.667041397 | 0.502433604 | 3.317933722 | 0.00090686 | 0.085897143 | Up | ENSMUSG00000033910.13 | Gucy1a1 |
| 258.529481 | 0.613736477 | 0.185849588 | 3.302328956 | 0.000958855 | 0.087881824 | Up | ENSMUSG00000030862.13 | Cpxm2 |
| 3997.958437 | 0.411795536 | 0.125670955 | 3.276775734 | 0.001049997 | 0.093443994 | Up | ENSMUSG00000031775.5 | Pllp |
| 23392.95803 | 0.405628854 | 0.124048888 | 3.269911255 | 0.001075812 | 0.093989171 | Up | ENSMUSG00000018593.13 | Sparc |
| 700.1460373 | -0.701394591 | 0.145617553 | -4.816689861 | 1.45959E-06 | 0.002626781 | Down | ENSMUSG00000014498.9 | Ankrd52 |
| 1302.133911 | -0.567875308 | 0.118984932 | -4.77266572 | 1.81803E-06 | 0.00294467 | Down | ENSMUSG00000023809.10 | Rps6ka2 |
| 1329.107816 | -0.675287679 | 0.144097331 | -4.686330241 | 2.78147E-06 | 0.003902114 | Down | ENSMUSG00000025582.4 | Nptx1 |
| 1993.815461 | -0.589682822 | 0.126487795 | -4.66197409 | 3.13191E-06 | 0.003902114 | Down | ENSMUSG00000032908.9 | Sgpp2 |
| 4808.856688 | -0.604134745 | 0.12951936 | -4.664435827 | 3.09465E-06 | 0.003902114 | Down | ENSMUSG00000046480.6 | Scn4b |
| 2864.079309 | -0.670219159 | 0.15257492 | -4.392721678 | 1.1194E-05 | 0.011331866 | Down | ENSMUSG00000051111.16 | Sv2c |
| 469.1282845 | -0.650008467 | 0.148914858 | -4.364967165 | 1.27142E-05 | 0.012113627 | Down | ENSMUSG00000054720.12 | Lrrc8c |
| 2006.150626 | -0.570545109 | 0.132183445 | -4.316312903 | 1.58657E-05 | 0.013525095 | Down | ENSMUSG00000056222.15 | Spock1 |
| 3123.108638 | -0.662122188 | 0.155067103 | -4.269907518 | 1.95554E-05 | 0.014562002 | Down | ENSMUSG00000034533.10 | Scn10a |
| 7843.157751 | -0.484235598 | 0.115333092 | -4.198583346 | 2.6859E-05 | 0.016583146 | Down | ENSMUSG00000027254.13 | Map1a |
| 24468.42919 | -0.385032093 | 0.091848018 | -4.192056635 | 2.76437E-05 | 0.016583146 | Down | ENSMUSG00000033161.10 | Atp1a1 |
| 6573.210462 | -0.589173948 | 0.140832811 | -4.18349918 | 2.87056E-05 | 0.016605172 | Down | ENSMUSG00000036062.13 | Phf24 |
| 3970.491026 | -0.540752894 | 0.132269828 | -4.088255824 | 4.34629E-05 | 0.021999003 | Down | ENSMUSG00000006342.15 | Susd2 |
| 2239.798927 | -0.6017432 | 0.149936679 | -4.013315504 | 5.98718E-05 | 0.027257085 | Down | ENSMUSG00000026452.15 | Syt2 |
| 1433.890515 | -0.382804193 | 0.095351356 | -4.014669619 | 5.95291E-05 | 0.027257085 | Down | ENSMUSG00000026819.15 | Slc25a25 |
| 3833.180326 | -0.511655164 | 0.127577932 | -4.010530327 | 6.05825E-05 | 0.027257085 | Down | ENSMUSG00000034981.9 | Parm1 |
| 5437.492512 | -0.459825969 | 0.114387895 | -4.019883122 | 5.8227E-05 | 0.027257085 | Down | ENSMUSG00000048070.4 | Pirt |
| 2023.008547 | -0.479063993 | 0.121014963 | -3.958717004 | 7.53535E-05 | 0.030512499 | Down | ENSMUSG00000009731.4 | Kcnd1 |
| 6678.292121 | -0.490492065 | 0.124156115 | -3.950607402 | 7.79531E-05 | 0.03079528 | Down | ENSMUSG00000030110.13 | Ret |
| 878.5955349 | -0.590148053 | 0.149739077 | -3.941175969 | 8.10831E-05 | 0.031269124 | Down | ENSMUSG00000007594.10 | Hapln4 |
| 1559.336605 | -0.533245716 | 0.135825267 | -3.925968458 | 8.63815E-05 | 0.032537692 | Down | ENSMUSG00000029822.15 | Osbpl3 |
| 2027.419364 | -0.381753945 | 0.097437031 | -3.917955426 | 8.93032E-05 | 0.032873731 | Down | ENSMUSG00000023328.14 | Ache |
| 303.3844777 | -0.638885413 | 0.1638665 | -3.898816482 | 9.6664E-05 | 0.03331206 | Down | ENSMUSG00000034460.9 | Six4 |
| 589.4206657 | -0.548349329 | 0.140323395 | -3.907754146 | 9.3158E-05 | 0.03331206 | Down | ENSMUSG00000044042.19 | Fmn1 |
| 4033.198212 | -0.425079248 | 0.108931126 | -3.902275365 | 9.52927E-05 | 0.03331206 | Down | ENSMUSG00000052852.8 | Reep1 |
| 1883.607414 | -0.557841056 | 0.144469011 | -3.861319833 | 0.000112776 | 0.0334648 | Down | ENSMUSG00000020261.15 | Slc36a1 |
| 1987.940336 | -0.523997521 | 0.135877466 | -3.856397509 | 0.00011507 | 0.0334648 | Down | ENSMUSG00000031391.18 | L1cam |
| 2464.508095 | -0.62311028 | 0.161495361 | -3.858378812 | 0.000114142 | 0.0334648 | Down | ENSMUSG00000031441.15 | Atp11a |
| 16710.55861 | -0.51752788 | 0.134246459 | -3.855057952 | 0.000115702 | 0.0334648 | Down | ENSMUSG00000058297.16 | Spock2 |
| 1298.058522 | -0.597909472 | 0.154299591 | -3.874990647 | 0.000106629 | 0.0334648 | Down | ENSMUSG00000074785.5 | Plxnc1 |
| 2081.878247 | -0.512077178 | 0.133720261 | -3.829465894 | 0.000128422 | 0.034263656 | Down | ENSMUSG00000020926.16 | Adam11 |
| 3308.652821 | -0.467939642 | 0.122062718 | -3.833600063 | 0.000126281 | 0.034263656 | Down | ENSMUSG00000024501.20 | Dpysl3 |
| 1274.154611 | -0.483610958 | 0.125833236 | -3.843268865 | 0.000121406 | 0.034263656 | Down | ENSMUSG00000039202.12 | Abhd2 |
| 613.6323492 | -0.629727011 | 0.164625363 | -3.825212592 | 0.000130659 | 0.034263656 | Down | ENSMUSG00000069072.9 | Slc7a14 |
| 1798.852502 | -0.378013263 | 0.099660606 | -3.793005847 | 0.000148835 | 0.036525376 | Down | ENSMUSG00000002032.17 | Tmem25 |
| 1775.606906 | -0.433482067 | 0.115507977 | -3.752832282 | 0.000174848 | 0.04104363 | Down | ENSMUSG00000031129.9 | Slc9a9 |
| 930.5688961 | -0.437843127 | 0.11682228 | -3.747941965 | 0.000178291 | 0.041254104 | Down | ENSMUSG00000000627.15 | Sema4f |
| 1132.867017 | -0.388876891 | 0.104548216 | -3.719593742 | 0.000199543 | 0.045521207 | Down | ENSMUSG00000028300.14 | C9orf72 |
| 1877.40397 | -0.500658777 | 0.135520219 | -3.694347476 | 0.000220452 | 0.049592553 | Down | ENSMUSG00000023915.5 | Tnfrsf21 |
| 70.60389507 | -1.530262723 | 0.417307187 | -3.666993455 | 0.000245419 | 0.053000703 | Down | ENSMUSG00000030680.6 | Pagr1a |
| 995.8679318 | -0.465911258 | 0.127733351 | -3.647530215 | 0.000264773 | 0.054574253 | Down | ENSMUSG00000020135.13 | Apc2 |
| 2223.065265 | -0.510688239 | 0.140309292 | -3.639732132 | 0.000272922 | 0.054574253 | Down | ENSMUSG00000034115.10 | Scn11a |
| 2804.401255 | -0.402470796 | 0.110501135 | -3.642232243 | 0.000270284 | 0.054574253 | Down | ENSMUSG00000036622.15 | Atp13a2 |
| 1365.235359 | -0.564171663 | 0.154884041 | -3.642542244 | 0.000269959 | 0.054574253 | Down | ENSMUSG00000041608.8 | Entpd3 |
| 6762.516399 | -0.483968068 | 0.133479026 | -3.625798632 | 0.00028807 | 0.055546025 | Down | ENSMUSG00000039126.10 | Prune2 |
| 2575.024802 | -0.419569979 | 0.115674575 | -3.627158158 | 0.000286558 | 0.055546025 | Down | ENSMUSG00000046157.13 | Tmem229b |
| 1190.788422 | -0.529075522 | 0.14649158 | -3.611644589 | 0.000304261 | 0.05665271 | Down | ENSMUSG00000036502.14 | Tmem255a |
| 1083.907961 | -0.425751957 | 0.118283447 | -3.599421303 | 0.000318926 | 0.058700529 | Down | ENSMUSG00000026430.16 | Rassf5 |
| 1257.999821 | -0.5245375 | 0.145959214 | -3.593726532 | 0.000325982 | 0.058874172 | Down | ENSMUSG00000020701.12 | Tmem132e |
| 285.758074 | -0.688091923 | 0.191519499 | -3.592803475 | 0.000327139 | 0.058874172 | Down | ENSMUSG00000103255.1 | Pcdhac1 |
| 413.522389 | -0.736081462 | 0.205043278 | -3.589883409 | 0.000330826 | 0.058883375 | Down | ENSMUSG00000036251.16 | Trpm8 |
| 309.388509 | -0.681354504 | 0.19018645 | -3.582560722 | 0.000340242 | 0.05990117 | Down | ENSMUSG00000034145.14 | Tmem63c |
| 472.4348936 | -0.570184638 | 0.160185516 | -3.559526797 | 0.000371524 | 0.062036785 | Down | ENSMUSG00000060212.13 | Pcnx2 |
| 2605.516569 | -0.399883513 | 0.11301499 | -3.538322783 | 0.000402677 | 0.064575912 | Down | ENSMUSG00000021360.16 | Gcnt2 |
| 1186.912056 | -0.606892115 | 0.172197746 | -3.524390591 | 0.000424458 | 0.066747061 | Down | ENSMUSG00000039809.10 | Gabbr2 |
| 351.609576 | -0.662586537 | 0.188792318 | -3.50960538 | 0.000448772 | 0.068573237 | Down | ENSMUSG00000025020.11 | Slit1 |
| 15023.74482 | -0.59748187 | 0.170451761 | -3.50528424 | 0.00045612 | 0.069044638 | Down | ENSMUSG00000052727.6 | Map1b |

| | | | | | | | | |
|---|---|---|---|---|---|---|---|---|
| 2645.886598 | -0.434369785 | 0.124187482 | -3.4976938 | 0.0004693 | 0.069736199 | Down | ENSMUSG00000024897.9 | Apba1 |
| 481.0137435 | -0.459915987 | 0.131923818 | -3.48622405 | 0.000489891 | 0.070846042 | Down | ENSMUSG00000005672.12 | Kit |
| 107.5050652 | -0.919802554 | 0.263702566 | -3.488030346 | 0.000486593 | 0.070846042 | Down | ENSMUSG00000040901.8 | Kcnk18 |
| 2720.199033 | -0.431606007 | 0.123732936 | -3.488206292 | 0.000486273 | 0.070846042 | Down | ENSMUSG00000061576.15 | Dpp6 |
| 2271.254727 | -0.488476832 | 0.140474676 | -3.477330185 | 0.000506434 | 0.071130127 | Down | ENSMUSG00000030554.16 | Synm |
| 492.5010232 | -0.469037453 | 0.135027335 | -3.473648148 | 0.000513434 | 0.071130127 | Down | ENSMUSG00000032649.14 | Colgalt2 |
| 1285.962636 | -0.52734557 | 0.151683289 | -3.47662273 | 0.000507772 | 0.071130127 | Down | ENSMUSG00000040430.18 | Pitpnc1 |
| 1352.67071 | -0.390643394 | 0.112235286 | -3.480575559 | 0.000500338 | 0.071130127 | Down | ENSMUSG00000054843.9 | Atrnl1 |
| 827.116068 | -0.490050542 | 0.141580832 | -3.461277462 | 0.000537618 | 0.072565044 | Down | ENSMUSG00000025272.16 | Tro |
| 260.0620402 | -0.635665659 | 0.18448086 | -3.445699788 | 0.000569583 | 0.073804255 | Down | ENSMUSG00000090061.9 | Nwd2 |
| 1932.142931 | -0.593488634 | 0.172368826 | -3.443132067 | 0.000575018 | 0.073917251 | Down | ENSMUSG00000021983.16 | Atp8a2 |
| 1036.41745 | -0.476026714 | 0.138846078 | -3.428449117 | 0.00060704 | 0.075055206 | Down | ENSMUSG00000049556.5 | Lingo1 |
| 1131.418565 | -0.495973446 | 0.145262168 | -3.414333216 | 0.000639383 | 0.076711783 | Down | ENSMUSG00000028631.7 | Kcnq4 |
| 2243.71339 | -0.411727739 | 0.120777051 | -3.408989826 | 0.000652039 | 0.077654982 | Down | ENSMUSG00000040265.16 | Dnm3 |
| 1094.317829 | -0.438268216 | 0.129027936 | -3.396692444 | 0.000682056 | 0.078656452 | Down | ENSMUSG00000023017.10 | Asic1 |
| 356.5713058 | -0.555260508 | 0.164148147 | -3.382679105 | 0.000717825 | 0.078656452 | Down | ENSMUSG00000030283.7 | St8sia1 |
| 1883.664872 | -0.441793039 | 0.130558024 | -3.383882704 | 0.000714685 | 0.078656452 | Down | ENSMUSG00000038276.12 | Asic3 |
| 1283.125727 | -0.482923984 | 0.14351664 | -3.364933734 | 0.00076562 | 0.080471342 | Down | ENSMUSG00000049313.8 | Sorl1 |
| 640.6829289 | -0.62328872 | 0.185967228 | -3.351605149 | 0.000803445 | 0.082363305 | Down | ENSMUSG00000051650.11 | B3gnt2 |
| 1020.563345 | -0.481361658 | 0.143565955 | -3.352895605 | 0.000799709 | 0.082363305 | Down | ENSMUSG00000058441.7 | Panx2 |
| 2008.598915 | -0.394044042 | 0.117720423 | -3.347286998 | 0.000816067 | 0.082611457 | Down | ENSMUSG00000024261.6 | Syt4 |
| 437.2601854 | -0.562830397 | 0.168086121 | -3.348464424 | 0.000812607 | 0.082611457 | Down | ENSMUSG00000042719.17 | Naa25 |
| 1193.714738 | -0.446412545 | 0.13381627 | -3.336010964 | 0.000849898 | 0.083937804 | Down | ENSMUSG00000053024.14 | Cntn2 |
| 1747.527372 | -0.41277193 | 0.124054837 | -3.32733442 | 0.000876811 | 0.085253518 | Down | ENSMUSG00000032220.10 | Myo1e |
| 530.4788535 | -0.556015253 | 0.166972283 | -3.329985327 | 0.000868506 | 0.085253518 | Down | ENSMUSG00000038859.7 | Baiap2l1 |
| 1172.764897 | -0.396034533 | 0.11904955 | -3.326636118 | 0.000879011 | 0.085253518 | Down | ENSMUSG00000052928.9 | Ctif |
| 978.4928614 | -0.430167964 | 0.129879822 | -3.31204614 | 0.000926163 | 0.087215449 | Down | ENSMUSG00000074582.10 | Arfgef2 |
| 2507.47632 | -0.404817244 | 0.12271898 | -3.298733766 | 0.00097122 | 0.087881824 | Down | ENSMUSG00000033949.12 | Trim36 |
| 2594.912849 | -0.438026353 | 0.132403491 | -3.30826892 | 0.000938746 | 0.087881824 | Down | ENSMUSG00000034731.11 | Dgkh |
| 1255.105969 | -0.582423062 | 0.176229544 | -3.304911593 | 0.000950064 | 0.087881824 | Down | ENSMUSG00000037996.17 | Slc24a2 |
| 960.7983002 | -0.590395765 | 0.178615256 | -3.305405036 | 0.000948392 | 0.087881824 | Down | ENSMUSG00000041482.17 | Piezo2 |
| 825.5260644 | -0.443572173 | 0.134671999 | -3.293722359 | 0.000988701 | 0.088966616 | Down | ENSMUSG00000024064.14 | Galnt14 |
| 861.7805526 | -0.462294464 | 0.141542196 | -3.266124716 | 0.001090302 | 0.093989171 | Down | ENSMUSG00000027827.17 | Kcnab1 |
| 420.0773754 | -0.558468993 | 0.170996941 | -3.265958969 | 0.001090941 | 0.093989171 | Down | ENSMUSG00000032769.5 | Trpa1 |
| 763.8245666 | -0.469760995 | 0.143720182 | -3.268580582 | 0.001080884 | 0.093989171 | Down | ENSMUSG00000036067.12 | Slc2a6 |
| 2406.544448 | -0.387096562 | 0.118353666 | -3.270676566 | 0.001072905 | 0.093989171 | Down | ENSMUSG00000048027.9 | Rgmb |
| 948.7202569 | -0.429552263 | 0.132380286 | -3.244835589 | 0.001175185 | 0.099974183 | Down | ENSMUSG00000038181.16 | Chpf2 |
| 618.7760125 | -0.418754633 | 0.129088669 | -3.243930209 | 0.001178926 | 0.099974183 | Down | ENSMUSG00000046321.8 | Hs3st2 |

DEGs Rag1KO DED vs Rag1KO Ct

| baseMean | log2FoldChange | lfcSE | stat | pvalue | padj | DE | gene_id | external_gene_name |
|---|---|---|---|---|---|---|---|---|
| 813.5626622 | 0.587328208 | 0.138709698 | 4.234225983 | 2.2934E-05 | 0.053072607 | Up | ENSMUSG00000001020.8 | S100a4 |
| 3703.868919 | 0.507207226 | 0.105670559 | 4.799891573 | 1.58752E-06 | 0.009975836 | Up | ENSMUSG00000023886.10 | Smoc2 |
| 7364.085924 | 0.429404891 | 0.100546607 | 4.270704933 | 1.94856E-05 | 0.053072607 | Up | ENSMUSG00000024610.15 | Cd74 |
| 235.2490367 | -0.910399181 | 0.196752679 | -4.627124706 | 3.70777E-06 | 0.015015553 | Down | ENSMUSG00000040584.8 | Abcb1a |
| 268.6780052 | 0.879582593 | 0.207383404 | 4.241335489 | 2.22194E-05 | 0.053072607 | Up | ENSMUSG00000052468.7 | Pmp2 |
| 20.26698415 | -3.445683496 | 0.722452038 | -4.769428717 | 1.84749E-06 | 0.009975836 | Down | ENSMUSG00000092386.1 | Gm20536 |
| 52.8573461 | 9.222531632 | 1.224961809 | 7.52883197 | 5.11962E-14 | 8.29327E-10 | Up | ENSMUSG00000107928.1 | Gm45140 |

DEGs Rag1KO DED vs WT DED

| baseMean | log2FoldChange | lfcSE | stat | pvalue | padj | DE | gene_id | external_gene_name |
|---|---|---|---|---|---|---|---|---|
| 947.7546889 | 1.121939377 | 0.131242875 | 8.548573617 | 1.24619E-17 | 3.36471E-14 | Up | ENSMUSG00000095463.8 | Entpd4 |
| 121.9515138 | 10.33391873 | 1.21365769 | 8.514689779 | 1.67037E-17 | 3.86571E-14 | Up | ENSMUSG00000093954.8 | Gm16867 |
| 942.7085222 | 1.110693572 | 0.131859021 | 8.42334159 | 3.65896E-17 | 7.40939E-14 | Up | ENSMUSG00000022066.16 | Entpd4b |
| 55.66838351 | 6.750405611 | 0.913161107 | 7.392349013 | 1.44257E-13 | 2.12452E-10 | Up | ENSMUSG00000090015.8 | Gm15446 |
| 170.9690142 | 2.175644666 | 0.348955966 | 6.234725526 | 4.52571E-10 | 5.23689E-07 | Up | ENSMUSG00000074634.12 | Tmem267 |
| 100.4813992 | 2.170414426 | 0.384001854 | 5.652093614 | 1.58505E-08 | 1.60487E-05 | Up | ENSMUSG00000050761.4 | Gp1bb |
| 73.43747656 | 5.905416587 | 1.103061534 | 5.35366016 | 8.61927E-08 | 8.21365E-05 | Up | ENSMUSG00000098975.7 | Gm27177 |
| 13.42880102 | 7.15124481 | 1.377864074 | 5.190094544 | 2.10187E-07 | 0.000179212 | Up | ENSMUSG00000058447.8 | Gm26920 |
| 157.4976234 | 1.531937509 | 0.300731247 | 5.094041695 | 3.5051E-07 | 0.000270393 | Up | ENSMUSG00000035299.16 | Mid1 |
| 1208.24652 | 0.853340574 | 0.169755882 | 5.026868952 | 4.98553E-07 | 0.000367116 | Up | ENSMUSG00000001420.13 | Tmem79 |
| 17.50669033 | 4.629438389 | 0.979649338 | 4.725607635 | 2.29428E-06 | 0.001334814 | Up | ENSMUSG00000072844.6 | G530011O06Rikx |
| 11.56759692 | 6.937879388 | 1.49182663 | 4.650593607 | 3.30981E-06 | 0.001787297 | Up | ENSMUSG00000090338.2 | Gm17081 |
| 4027.345626 | 0.546870808 | 0.120485324 | 4.538899758 | 5.65485E-06 | 0.002862768 | Up | ENSMUSG00000031431.13 | Tsc22d3 |
| 50.89423704 | 2.112389968 | 0.470012654 | 4.494325742 | 6.97907E-06 | 0.003325323 | Up | ENSMUSG00000095366.2 | G530011O06Riky |
| 2319.195107 | 0.650693967 | 0.145506033 | 4.471938068 | 7.75139E-06 | 0.003587785 | Up | ENSMUSG00000020108.4 | Ddit4 |
| 22.76798627 | 3.632631056 | 0.81641962 | 4.449465648 | 8.60842E-06 | 0.003769092 | Up | ENSMUSG00000091542.1 | Gm17167 |
| 716.3524726 | 0.659493896 | 0.153704208 | 4.290669093 | 1.78136E-05 | 0.007399478 | Up | ENSMUSG00000020893.17 | Per1 |
| 1983.98004 | 0.562830555 | 0.137955546 | 4.079796501 | 4.50751E-05 | 0.017810179 | Up | ENSMUSG00000038276.12 | Asic3 |
| 2511.893637 | 0.521207447 | 0.128178891 | 4.066250228 | 4.77756E-05 | 0.018427746 | Up | ENSMUSG00000030284.11 | Creld1 |
| 903.7720527 | 0.651521944 | 0.161391258 | 4.036909755 | 5.41599E-05 | 0.02017395 | Up | ENSMUSG00000007594.10 | Hapln4 |
| 12.01615868 | 5.113856977 | 1.310485793 | 3.902260523 | 9.52985E-05 | 0.032163246 | Up | ENSMUSG00000118434.1 | Gm13301 |
| 837.8537317 | 0.540593255 | 0.139030632 | 3.888303233 | 0.000100947 | 0.033374466 | Up | ENSMUSG00000000958.10 | Slc7a7 |
| 115.3913129 | 1.55435738 | 0.40372934 | 3.849998565 | 0.000118119 | 0.038270399 | Up | ENSMUSG00000072676.12 | Tmem254 |
| 319.7938842 | 0.75331104 | 0.19638764 | 3.83583732 | 0.000125137 | 0.039493352 | Up | ENSMUSG00000034145.14 | Tmem63c |
| 48.93819328 | 1.558728508 | 0.41243401 | 3.779340381 | 0.000157244 | 0.0446905 | Up | ENSMUSG00000063458.13 | Lrmda |
| 230.6124689 | 0.78150239 | 0.207917451 | 3.758714749 | 0.000170788 | 0.047702971 | Up | ENSMUSG00000005994.14 | Tyrp1 |
| 10.66344176 | 4.323590043 | 1.15436608 | 3.745423672 | 0.00018009 | 0.04944833 | Up | ENSMUSG00000095304.9 | Plac9 |
| 4002.113884 | 0.553390821 | 0.148486187 | 3.726884187 | 0.000193861 | 0.051484525 | Up | ENSMUSG00000006342.15 | Susd2 |
| 460.0247566 | 0.61187641 | 0.164045462 | 3.729919761 | 0.000191541 | 0.051484525 | Up | ENSMUSG00000034684.12 | Sema3f |
| 1846.33736 | 0.438121914 | 0.118730928 | 3.690040332 | 0.000224219 | 0.056755308 | Up | ENSMUSG00000002032.17 | Tmem25 |
| 1284.051712 | 0.456721622 | 0.123985169 | 3.683679481 | 0.000229891 | 0.057295981 | Up | ENSMUSG00000058454.15 | Dhcr7 |
| 1285.55709 | 0.570860721 | 0.156021671 | 3.658855314 | 0.000253344 | 0.060355557 | Up | ENSMUSG00000020701.12 | Tmem132e |
| 1274.848388 | 0.510533267 | 0.139456861 | 3.660868781 | 0.000251361 | 0.060355557 | Up | ENSMUSG00000023809.10 | Rps6ka2 |
| 1387.054564 | 0.595418462 | 0.162728699 | 3.658964053 | 0.000253237 | 0.060355557 | Up | ENSMUSG00000041608.8 | Entpd3 |
| 3426.4729 | 0.483028083 | 0.134406968 | 3.59377264 | 0.000325924 | 0.073472562 | Up | ENSMUSG00000025366.8 | Esyt1 |
| 3062.838081 | 0.609446574 | 0.173011293 | 3.522582615 | 0.000427364 | 0.091095972 | Up | ENSMUSG00000034533.10 | Scn10a |
| 949.587146 | 0.462452166 | 0.131497469 | 3.516814203 | 0.000436759 | 0.091889638 | Up | ENSMUSG00000071637.5 | Cebpd |
| 876.5918567 | 0.50780245 | 0.145152217 | 3.498413327 | 0.000468035 | 0.097207308 | Up | ENSMUSG00000045216.7 | Hs6st1 |
| 687.1510179 | 2.037784667 | 0.583912329 | 3.489881212 | 0.000483235 | 0.09909382 | Up | ENSMUSG00000096768.8 | Erdr1y |
| 1580.520619 | -2.346333932 | 0.130152873 | -18.02752322 | 1.18481E-72 | 1.9194E-68 | Down | ENSMUSG00000032679.12 | Cd59a |
| 455.5051916 | -2.646302216 | 0.173818521 | -15.22451234 | 2.43165E-52 | 1.96964E-48 | Down | ENSMUSG00000073643.11 | Wdfy1 |
| 358.0996631 | -1.849588974 | 0.187148888 | -9.882981384 | 4.93426E-23 | 1.5987E-19 | Down | ENSMUSG00000000560.9 | Gabra2 |
| 257.6747186 | -1.669372251 | 0.210140805 | -7.94406517 | 1.95661E-15 | 3.52189E-12 | Down | ENSMUSG00000117465.1 | Gm49980 |
| 586.5231601 | -1.1408797 | 0.160168796 | -7.122983549 | 1.05615E-12 | 1.42581E-09 | Down | ENSMUSG00000096255.2 | Dynlt1b |
| 50.83287407 | -3.311669066 | 0.556964001 | -5.945930182 | 2.74891E-09 | 2.96883E-06 | Down | ENSMUSG00000040592.11 | Cd79b |
| 26.78565016 | -5.32436198 | 1.012912647 | -5.256486819 | 1.46833E-07 | 0.00013215 | Down | ENSMUSG00000008193.13 | Spib |
| 223.5066849 | -1.173294628 | 0.228661341 | -5.131145567 | 2.87984E-07 | 0.000233267 | Down | ENSMUSG00000096780.7 | Tmem181b-ps |
| 220.1680579 | -1.052449598 | 0.212407825 | -4.954853237 | 7.23849E-07 | 0.000509841 | Down | ENSMUSG00000029484.12 | Anxa3 |
| 12.91423538 | -7.177360554 | 1.458265925 | -4.921846167 | 8.57316E-07 | 0.000578688 | Down | ENSMUSG00000005540.10 | Fcer2a |
| 39.33836863 | -3.211822962 | 0.679682843 | -4.725473058 | 2.2958E-06 | 0.001334814 | Down | ENSMUSG00000029819.6 | Npy |
| 12.81943251 | -7.16656525 | 1.516901351 | -4.724476806 | 2.30709E-06 | 0.001334814 | Down | ENSMUSG00000076937.3 | Iglc2 |
| 38.23288032 | -3.471664107 | 0.737464243 | -4.707569404 | 2.50688E-06 | 0.001400395 | Down | ENSMUSG00000104213.5 | Ighd |
| 17.16364914 | -6.612238771 | 1.444202295 | -4.578471309 | 4.68386E-06 | 0.002447697 | Down | ENSMUSG00000034634.7 | Ly6d |
| 18.82217715 | -3.986221151 | 0.881055691 | -4.524369107 | 6.05759E-06 | 0.002973727 | Down | ENSMUSG00000076498.2 | Trbc2 |
| 20.81166089 | -4.985240116 | 1.119386332 | -4.453547423 | 8.4463E-06 | 0.003769092 | Down | ENSMUSG00000042474.6 | Fcmr |
| 52366.58155 | -0.580254252 | 0.132909964 | -4.365769387 | 1.26676E-05 | 0.005400401 | Down | ENSMUSG00000064354.1 | mt-Co2 |
| 90.46608231 | -1.832693205 | 0.454291449 | -4.034179396 | 5.47934E-05 | 0.02017395 | Down | ENSMUSG00000003379.7 | Cd79a |
| 12342.87191 | -0.557374671 | 0.140390657 | -3.970169272 | 7.18216E-05 | 0.025855772 | Down | ENSMUSG00000027273.13 | Snap25 |
| 42643.02486 | -0.573873292 | 0.144748554 | -3.964621926 | 7.35124E-05 | 0.025889137 | Down | ENSMUSG00000064357.1 | mt-Atp6 |
| 44.91424811 | -2.125126711 | 0.544118162 | -3.905634585 | 9.39784E-05 | 0.032163246 | Down | ENSMUSG00000024399.5 | Ltb |
| 47.49643859 | -1.643966762 | 0.428937038 | -3.832652853 | 0.000126769 | 0.039493352 | Down | ENSMUSG00000073791.12 | Efcab7 |
| 580.3382773 | -0.825064028 | 0.21602843 | -3.819238173 | 0.000133864 | 0.040917063 | Down | ENSMUSG00000038418.7 | Egr1 |
| 66679.95271 | -0.556856159 | 0.146286883 | -3.806603494 | 0.000140888 | 0.042266505 | Down | ENSMUSG00000064370.1 | mt-Cytb |
| 36.20480243 | -2.703957838 | 0.714491437 | -3.784451009 | 0.000154048 | 0.0446905 | Down | ENSMUSG00000026581.14 | Sell |
| 94.85206243 | -7.546508008 | 1.995973225 | -3.780866353 | 0.000156284 | 0.0446905 | Down | ENSMUSG00000076609.2 | Igkc |
| 23.40318736 | -7.06471563 | 1.910004869 | -3.698794566 | 0.000216626 | 0.056602237 | Down | ENSMUSG00000030724.7 | Cd19 |
| 32.70561742 | -1.943641182 | 0.526322024 | -3.692874505 | 0.000221733 | 0.056755308 | Down | ENSMUSG00000043263.13 | Ifi209 |

| | | | | | | | | |
|---|---|---|---|---|---|---|---|---|
| 47743.5433 | -0.49892094 | 0.13719033 | -3.636706305 | 0.000276146 | 0.064834392 | Down | ENSMUSG00000064341.1 | mt-Nd1 |
| 1882.679774 | -0.46779691 | 0.130186698 | -3.593277323 | 0.000326545 | 0.073472562 | Down | ENSMUSG00000059824.12 | Dbp |
| 22.5340376 | -2.243028852 | 0.623064799 | -3.599992896 | 0.000318226 | 0.073472562 | Down | ENSMUSG00000110386.1 | Gm42031 |
| 107696.4891 | -0.482320113 | 0.134715287 | -3.580292374 | 0.00034321 | 0.076164403 | Down | ENSMUSG00000064358.1 | mt-Co3 |
| 5.606049755 | -5.972604439 | 1.675476401 | -3.564720121 | 0.000364245 | 0.079740046 | Down | ENSMUSG00000003882.5 | Il7r |
| 575.2149134 | -0.756210397 | 0.212878358 | -3.552312246 | 0.000381861 | 0.082482068 | Down | ENSMUSG00000076617.9 | Ighm |
