## Supplementary material for "CD4^+^ T cells drive corneal nerve damage but not epitheliopathy in an acute aqueous-deficient dry eye model": Dataset S2

| baseMean | log2FoldChange | lfcSE | stat | pvalue | padj | DE | gene_id | external_gene_name |
|---|---|---|---|---|---|---|---|---|
| 24.80591921 | -22.6352404 | 2.763333988 | -8.191279268 | 2.58464E-16 | 4.37244E-12 | Down | ENSMUSG00000093769.4 | H3c14 |
| 6108.638864 | 3.256372982 | 0.42573362 | 7.648850905 | 2.02783E-14 | 1.71524E-10 | Up | ENSMUSG00000025270.13 | Alas2 |
| 26.12085046 | 21.12451323 | 2.848661968 | 7.415591413 | 1.21083E-13 | 6.82788E-10 | Up | ENSMUSG00000089943.1 | Ugt1a5 |
| 196187.102 | 2.279361888 | 0.317695653 | 7.174671311 | 7.24811E-13 | 3.06541E-09 | Up | ENSMUSG00000069917.7 | Hba-a2 |
| 155544.9687 | 2.364053792 | 0.331716806 | 7.126723014 | 1.02787E-12 | 3.47768E-09 | Up | ENSMUSG00000069919.7 | Hba-a1 |
| 110.4582477 | 9.72840486 | 1.383396057 | 7.032262967 | 2.0321E-12 | 5.72952E-09 | Up | ENSMUSG00000079224.5 | Gm6565 |
| 42926.28819 | -1.411675209 | 0.250490085 | -5.635653033 | 1.74396E-08 | 4.21466E-05 | Down | ENSMUSG00000035202.8 | Lars2 |
| 47.88242595 | 8.522315326 | 1.592471826 | 5.351627066 | 8.71669E-08 | 0.000184325 | Up | ENSMUSG00000103442.5 | Pcdha1 |
| 41.37107185 | 8.31158074 | 1.634795214 | 5.084172421 | 3.69232E-07 | 0.000694034 | Up | ENSMUSG00000092368.1 | A930015D03Rik |
| 2509.437651 | -1.131713305 | 0.223686251 | -5.059378031 | 4.20626E-07 | 0.000702863 | Down | ENSMUSG00000001420.13 | Tmem79 |
| 2740.30546 | 1.711510196 | 0.339347792 | 5.043528314 | 4.57025E-07 | 0.000702863 | Up | ENSMUSG00000038871.5 | Bpgm |
| 696.0549796 | 3.050048266 | 0.619923487 | 4.920039857 | 8.65266E-07 | 0.001125977 | Up | ENSMUSG00000039236.18 | Isg20 |
| 36548.05685 | -1.43640297 | 0.291069043 | -4.934921811 | 8.01829E-07 | 0.001125977 | Down | ENSMUSG00000039278.10 | Pcsk1n |
| 3850.644126 | -1.275326682 | 0.260505183 | -4.895590436 | 9.8011E-07 | 0.001184323 | Down | ENSMUSG00000090071.4 | Cdk5r2 |
| 2409.752716 | -1.255033159 | 0.257850004 | -4.86729937 | 1.13134E-06 | 0.001275921 | Down | ENSMUSG00000075227.6 | Znhit2 |
| 4860.2446 | -1.223815122 | 0.252878434 | -4.839539309 | 1.3014E-06 | 0.001295051 | Down | ENSMUSG00000037843.6 | Vstm2l |
| 110.5699207 | 7.117071052 | 1.467299277 | 4.850456322 | 1.23178E-06 | 0.001295051 | Up | ENSMUSG00000059108.4 | Ifitm6 |
| 625.9011203 | 3.001171385 | 0.630865504 | 4.757228549 | 1.96269E-06 | 0.001844601 | Up | ENSMUSG00000078921.3 | Tgtp2 |
| 31.95672573 | 7.938832175 | 1.702992226 | 4.661696074 | 3.13614E-06 | 0.002792321 | Up | ENSMUSG00000020914.17 | Top2a |
| 1602.139392 | 0.976361528 | 0.211344562 | 4.619761764 | 3.84181E-06 | 0.003249594 | Up | ENSMUSG00000040549.16 | Ckap5 |
| 1538.403319 | -1.160449128 | 0.252178028 | -4.601705924 | 4.19045E-06 | 0.003331551 | Down | ENSMUSG00000048385.9 | Scrt1 |
| 5076.706143 | -1.402374139 | 0.305211924 | -4.59475541 | 4.33257E-06 | 0.003331551 | Down | ENSMUSG00000078440.10 | Dohh |
| 431.1394249 | -1.414371366 | 0.30958833 | -4.568555167 | 4.91098E-06 | 0.003612131 | Down | ENSMUSG00000001076.7 | C1ql4 |
| 465.1048951 | 1.4866814 | 0.32766533 | 4.53719471 | 5.70075E-06 | 0.004018314 | Up | ENSMUSG00000035726.8 | Supt16 |
| 1703.559686 | -1.018382716 | 0.225293062 | -4.520257777 | 6.17644E-06 | 0.004179472 | Down | ENSMUSG00000040390.13 | Map3k10 |
| 45.75062422 | 8.456144737 | 1.878146253 | 4.502388843 | 6.71939E-06 | 0.004210071 | Up | ENSMUSG00000009350.13 | Mpo |
| 1336.817112 | -1.162800737 | 0.258073272 | -4.505699988 | 6.61545E-06 | 0.004210071 | Down | ENSMUSG00000022758.14 | P2rx6 |
| 321.0133442 | 2.856738086 | 0.639100016 | 4.469939 | 7.82419E-06 | 0.004412345 | Up | ENSMUSG00000020641.16 | Rsad2 |
| 49.69969827 | 6.526152785 | 1.457928248 | 4.476319596 | 7.59408E-06 | 0.004412345 | Up | ENSMUSG00000027306.15 | Nusap1 |
| 1659.693819 | 0.948539088 | 0.212204687 | 4.46992524 | 7.82469E-06 | 0.004412345 | Up | ENSMUSG00000032018.14 | Sc5d |
| 2587.153831 | -0.98690026 | 0.223783555 | -4.410066059 | 1.03339E-05 | 0.004925961 | Down | ENSMUSG00000000253.13 | Gmpr |
| 3165.001203 | -1.069859347 | 0.241209005 | -4.435403842 | 9.18998E-06 | 0.004925961 | Down | ENSMUSG00000002058.13 | Unc119 |
| 5444.834127 | -1.288337298 | 0.291897604 | -4.413661781 | 1.01637E-05 | 0.004925961 | Down | ENSMUSG00000020308.7 | Tpgs1 |
| 1955.144386 | -1.370230391 | 0.310923403 | -4.406970911 | 1.04826E-05 | 0.004925961 | Down | ENSMUSG00000029725.10 | Ppp1r35 |
| 4856.485456 | -1.009531958 | 0.228561493 | -4.416894309 | 1.00129E-05 | 0.004925961 | Down | ENSMUSG00000035585.16 | Tsen34 |
| 80523.27399 | -1.012210833 | 0.229374835 | -4.412911438 | 1.0199E-05 | 0.004925961 | Down | ENSMUSG00000062380.4 | Tubb3 |
| 33577.22975 | -1.279059812 | 0.291688224 | -4.385023824 | 1.15973E-05 | 0.005248907 | Down | ENSMUSG00000003380.11 | Rabac1 |
| 126.524247 | 4.240503094 | 0.967835912 | 4.38142772 | 1.17904E-05 | 0.005248907 | Up | ENSMUSG00000073489.6 | Ifi204 |
| 435.9163667 | -1.905311538 | 0.435569653 | -4.374298178 | 1.21824E-05 | 0.00528434 | Down | ENSMUSG00000048967.16 | Yjefn3 |
| 7671.806969 | -0.942783816 | 0.216409137 | -4.356488041 | 1.32166E-05 | 0.005453294 | Down | ENSMUSG00000006651.8 | Aplp1 |
| 1159.188008 | -1.60324335 | 0.367595782 | -4.361430213 | 1.29215E-05 | 0.005453294 | Down | ENSMUSG00000065037.1 | |
| 39793.5692 | -1.088015597 | 0.251286067 | -4.329788786 | 1.49252E-05 | 0.006011674 | Down | ENSMUSG00000001270.9 | Ckb |
| 3379.34744 | -1.331721239 | 0.308314547 | -4.319359085 | 1.56483E-05 | 0.006156332 | Down | ENSMUSG00000030588.12 | Yif1b |
| 1656.318268 | -1.449991488 | 0.336479555 | -4.309300417 | 1.63772E-05 | 0.006220893 | Down | ENSMUSG00000005699.16 | Pard6a |
| 25006.25869 | -1.038410869 | 0.24109804 | -4.307006675 | 1.65479E-05 | 0.006220893 | Down | ENSMUSG00000024121.13 | Atp6v0c |
| 7260.991804 | -1.238853024 | 0.28865645 | -4.291790548 | 1.77238E-05 | 0.006518124 | Down | ENSMUSG00000079598.4 | Clec2l |
| 34.26398715 | 8.039502616 | 1.884829487 | 4.265373962 | 1.99568E-05 | 0.006519604 | Up | ENSMUSG00000015437.5 | Gzmb |
| 13234.6067 | -1.274040588 | 0.298306831 | -4.27090651 | 1.9468E-05 | 0.006519604 | Down | ENSMUSG00000022193.7 | Psmb5 |
| 578.0451044 | -1.407986455 | 0.331457418 | -4.247865268 | 2.15817E-05 | 0.006519604 | Down | ENSMUSG00000023232.17 | Serinc2 |
| 15805.80246 | -1.085918526 | 0.254348443 | -4.269412908 | 1.95988E-05 | 0.006519604 | Down | ENSMUSG00000025651.14 | Uqcrc1 |
| 5278.027786 | -0.987069215 | 0.232360854 | -4.248001321 | 2.15686E-05 | 0.006519604 | Down | ENSMUSG00000026817.14 | Ak1 |
| 1265.664569 | -2.262405746 | 0.532070579 | -4.252078271 | 2.11796E-05 | 0.006519604 | Down | ENSMUSG00000030680.6 | Pagr1a |
| 461.090862 | -1.309512959 | 0.305579268 | -4.285346218 | 1.82455E-05 | 0.006519604 | Down | ENSMUSG00000033857.12 | Engase |
| 18890.17027 | -1.098191335 | 0.257544239 | -4.264088142 | 2.0072E-05 | 0.006519604 | Down | ENSMUSG00000041556.8 | Fbxo2 |
| 702.3978545 | -1.35783496 | 0.31839211 | -4.264662715 | 2.00204E-05 | 0.006519604 | Down | ENSMUSG00000064254.6 | Ethe1 |
| 1496.396401 | -1.141736691 | 0.268674414 | -4.249517746 | 2.14231E-05 | 0.006519604 | Down | ENSMUSG00000086784.2 | Isoc2a |
| 650.7782517 | -1.312602517 | 0.310948942 | -4.221279898 | 2.42919E-05 | 0.007209582 | Down | ENSMUSG00000022769.9 | Sdf2l1 |
| 4395.920099 | -0.89318938 | 0.212612857 | -4.20101301 | 2.65723E-05 | 0.007492069 | Down | ENSMUSG00000007950.9 | Abhd8 |
| 121.1065014 | 4.0241189 | 0.957491777 | 4.202771236 | 2.63667E-05 | 0.007492069 | Up | ENSMUSG00000051839.7 | Gypa |
| 5098.485049 | -1.100801057 | 0.261755474 | -4.205455733 | 2.60557E-05 | 0.007492069 | Down | ENSMUSG00000058966.13 | Tlcd3b |
| 29.07911725 | 6.870942787 | 1.641909209 | 4.184727601 | 2.85508E-05 | 0.007887655 | Up | ENSMUSG00000078903.10 | Gm14391 |
| 18.40050804 | -8.186294806 | 1.959256992 | -4.178264944 | 2.93741E-05 | 0.007887655 | Down | ENSMUSG00000091345.9 | Col6a5 |
| 39.04954977 | 7.208760402 | 1.725006615 | 4.178975511 | 2.92825E-05 | 0.007887655 | Up | ENSMUSG00000102037.1 | Bcl2a1a |
| 19226.85135 | -1.198979559 | 0.287806155 | -4.165927435 | 3.10089E-05 | 0.008196533 | Down | ENSMUSG00000033379.13 | Atp6v0b |
| 1465.519059 | -1.150994662 | 0.27665749 | -4.160359669 | 3.17747E-05 | 0.008222248 | Down | ENSMUSG00000002393.14 | Nr2f6 |
| 5103.30996 | -1.277984582 | 0.307403888 | -4.157346846 | 3.21965E-05 | 0.008222248 | Down | ENSMUSG00000007944.8 | Ttc9b |
| 4589.718881 | -0.987688719 | 0.237725179 | -4.154750139 | 3.25643E-05 | 0.008222248 | Down | ENSMUSG00000023909.4 | Paqr4 |
| 7977.662861 | -1.102998358 | 0.265964017 | -4.147171378 | 3.36608E-05 | 0.008252753 | Down | ENSMUSG00000020440.13 | Arf5 |
| 941.8186272 | -1.155222731 | 0.278366526 | -4.150005922 | 3.32467E-05 | 0.008252753 | Down | ENSMUSG00000039199.6 | Zdhhc1 |
| 10495.40611 | -0.976386392 | 0.236114043 | -4.135232199 | 3.54596E-05 | 0.00856957 | Down | ENSMUSG00000004951.10 | Hspb1 |
| 2341.274684 | -0.953467479 | 0.231401128 | -4.120409813 | 3.78199E-05 | 0.009011259 | Down | ENSMUSG00000003657.9 | Calb2 |
| 5244.736795 | -0.97249266 | 0.236739261 | -4.107863876 | 3.99335E-05 | 0.009382716 | Down | ENSMUSG00000062683.11 | Atp5mc2 |
| 212.882525 | 2.551067539 | 0.623196849 | 4.093518032 | 4.24877E-05 | 0.009846088 | Up | ENSMUSG00000079419.4 | Ms4a6c |
| 824.9009646 | -1.010957171 | 0.248188535 | -4.073343558 | 4.6343E-05 | 0.010594382 | Down | ENSMUSG00000036278.7 | Macrod1 |
| 24.41692317 | 7.550621056 | 1.856027722 | 4.068161789 | 4.73855E-05 | 0.010688268 | Up | ENSMUSG00000105881.1 | 4932422M17Rik |

| | | | | | | | | |
|---|---|---|---|---|---|---|---|---|
| 874.7945618 | -1.051994125 | 0.259012186 | -4.061562275 | 4.87454E-05 | 0.01070943 | Down | ENSMUSG00000020921.6 | Tmem101 |
| 2660.947768 | -0.949618834 | 0.233689903 | -4.063585209 | 4.83247E-05 | 0.01070943 | Down | ENSMUSG00000028049.15 | Scamp3 |
| 21625.19576 | -1.045317028 | 0.257867572 | -4.053697091 | 5.04145E-05 | 0.010894457 | Down | ENSMUSG00000024012.18 | Mtch1 |
| 1580.511018 | -1.038890883 | 0.25660343 | -4.048624308 | 5.15196E-05 | 0.010894457 | Down | ENSMUSG00000039483.10 | Asb6 |
| 3018.964592 | -0.849297764 | 0.209709812 | -4.049871372 | 5.12458E-05 | 0.010894457 | Down | ENSMUSG00000117679.1 | Apbb3 |
| 8767.973978 | -1.209516183 | 0.299785532 | -4.034604919 | 5.46942E-05 | 0.011147742 | Down | ENSMUSG00000003346.14 | Abhd17a |
| 1786.041914 | -1.118071706 | 0.277018113 | -4.036096035 | 5.4348E-05 | 0.011147742 | Down | ENSMUSG00000020684.14 | Rasl10b |
| 18154.99274 | 1.313044191 | 0.325073339 | 4.039224481 | 5.36282E-05 | 0.011147742 | Up | ENSMUSG00000064367.1 | mt-Nd5 |
| 1915.015476 | -1.037985307 | 0.257790209 | -4.026472957 | 5.66197E-05 | 0.011402813 | Down | ENSMUSG00000004929.12 | Thop1 |
| 1194.552996 | 0.969748745 | 0.241193893 | 4.020618988 | 5.80454E-05 | 0.011418074 | Up | ENSMUSG00000020124.10 | Usp15 |
| 23.12433611 | 7.471938241 | 1.857831491 | 4.021860044 | 5.77404E-05 | 0.011418074 | Up | ENSMUSG00000114608.1 | Gm36161 |
| 1844.820884 | -0.911641465 | 0.227343146 | -4.009979978 | 6.07239E-05 | 0.011807657 | Down | ENSMUSG00000035278.9 | Plekhj1 |
| 13727.69904 | -1.139634789 | 0.28573991 | -3.988364059 | 6.65305E-05 | 0.012148716 | Down | ENSMUSG00000008140.18 | Emc10 |
| 5410.450661 | -1.068638117 | 0.267303169 | -3.997850531 | 6.39203E-05 | 0.012148716 | Down | ENSMUSG00000021493.15 | Pdlim7 |
| 41.2829842 | 6.213640922 | 1.556535703 | 3.991968131 | 6.55272E-05 | 0.012148716 | Up | ENSMUSG00000031264.13 | Btk |
| 3905.093642 | -0.888033397 | 0.222484867 | -3.991432813 | 6.56753E-05 | 0.012148716 | Down | ENSMUSG00000049339.16 | Retreg2 |
| 5940.751559 | -1.11338295 | 0.279221652 | -3.987452051 | 6.67867E-05 | 0.012148716 | Down | ENSMUSG00000049422.7 | Chchd10 |
| 221.5365723 | -2.059700718 | 0.515660473 | -3.994296299 | 6.48867E-05 | 0.012148716 | Down | ENSMUSG00000092837.1 | Rpph1 |
| 35943.99116 | -0.755542801 | 0.189955928 | -3.977463673 | 6.96543E-05 | 0.012535544 | Down | ENSMUSG00000018865.9 | Sult4a1 |
| 3096.587387 | -0.967132211 | 0.243459223 | -3.972460759 | 7.1134E-05 | 0.012667086 | Down | ENSMUSG00000022516.10 | Nudt16l1 |
| 13272.15845 | -0.956517617 | 0.241050228 | -3.968125758 | 7.24401E-05 | 0.012765305 | Down | ENSMUSG00000022564.7 | Grina |
| 23.45916918 | 7.492846141 | 1.889858765 | 3.964765134 | 7.34682E-05 | 0.012813014 | Up | ENSMUSG00000040026.8 | Saa3 |
| 9891.677487 | -1.173904815 | 0.296870995 | -3.954259037 | 7.67722E-05 | 0.013252605 | Down | ENSMUSG00000093989.1 | Rnasek |
| 821.7770081 | -1.134768696 | 0.287722333 | -3.94397155 | 8.01432E-05 | 0.013694768 | Down | ENSMUSG00000020087.5 | Tysnd1 |
| 3215.091842 | -1.122594916 | 0.28544437 | -3.932797536 | 8.3963E-05 | 0.014204013 | Down | ENSMUSG00000047423.10 | AI837181 |
| 5212.116845 | -1.264006681 | 0.321952589 | -3.92606466 | 8.6347E-05 | 0.014462687 | Down | ENSMUSG00000095098.2 | Ccdc85b |
| 1113.895055 | -0.978830074 | 0.249910333 | -3.9167251 | 8.976E-05 | 0.014886961 | Down | ENSMUSG00000034880.7 | Mrpl34 |
| 2839.432281 | -0.924685866 | 0.236242515 | -3.914138256 | 9.07276E-05 | 0.014901355 | Down | ENSMUSG00000039611.2 | Pgap4 |
| 41.23117285 | 6.20098911 | 1.585793305 | 3.910338813 | 9.21667E-05 | 0.014992162 | Up | ENSMUSG00000021624.9 | Cd180 |
| 11401.88466 | -1.030424777 | 0.263934521 | -3.904092468 | 9.45796E-05 | 0.015094372 | Down | ENSMUSG00000019194.15 | Scn1b |
| 9473.333847 | -1.192440056 | 0.30539213 | -3.904619463 | 9.43738E-05 | 0.015094372 | Down | ENSMUSG00000072772.3 | Grcc10 |
| 5988.91923 | -0.964604095 | 0.247232314 | -3.901610113 | 9.5555E-05 | 0.01510751 | Down | ENSMUSG00000074457.10 | S100a16 |
| 44228.45704 | -0.895183111 | 0.230045217 | -3.891335462 | 9.9694E-05 | 0.01543012 | Down | ENSMUSG00000016349.10 | Eef1a2 |
| 1754.075861 | -1.2012353 | 0.308924583 | -3.888441922 | 0.00010089 | 0.01543012 | Down | ENSMUSG00000028743.7 | Akr7a5 |
| 3601.571036 | -0.833094821 | 0.214295887 | -3.887591284 | 0.000101244 | 0.01543012 | Down | ENSMUSG00000046157.13 | Tmem229b |
| 673.4173656 | -1.172494112 | 0.301574485 | -3.887908865 | 0.000101112 | 0.01543012 | Down | ENSMUSG00000051373.5 | Plpp7 |
| 2471.596791 | -1.116068698 | 0.287561208 | -3.881151791 | 0.000103963 | 0.015468502 | Down | ENSMUSG00000002820.6 | Atg4d |
| 147.9268618 | 3.200490727 | 0.824760943 | 3.880507137 | 0.000104239 | 0.015468502 | Up | ENSMUSG00000006574.15 | Slc4a1 |
| 2590.948122 | -1.129332817 | 0.291018264 | -3.880625232 | 0.000104188 | 0.015468502 | Down | ENSMUSG00000037204.7 | Atg101 |
| 2565.63802 | -1.133018602 | 0.292177036 | -3.877849598 | 0.000105384 | 0.01550242 | Down | ENSMUSG00000050373.13 | Snx21 |
| 66.6277627 | 3.247699708 | 0.839008063 | 3.870880209 | 0.000108443 | 0.015704208 | Up | ENSMUSG00000019936.10 | Epyc |
| 8138.67946 | -0.983284911 | 0.254045941 | -3.870500386 | 0.000108612 | 0.015704208 | Down | ENSMUSG00000056413.16 | Adap1 |
| 3152.870332 | -1.09438728 | 0.28347395 | -3.86062733 | 0.000113096 | 0.016078642 | Down | ENSMUSG00000024958.13 | Gpr137 |
| 5523.166901 | -0.91734109 | 0.237615368 | -3.860613469 | 0.000113103 | 0.016078642 | Down | ENSMUSG00000031765.8 | Mt1 |
| 8932.997223 | -1.00214553 | 0.259944997 | -3.855221452 | 0.000115625 | 0.016300222 | Down | ENSMUSG00000020153.14 | Ndufs7 |
| 690.240337 | -1.618003885 | 0.420323191 | -3.849428057 | 0.000118394 | 0.016416971 | Down | ENSMUSG00000029343.17 | Crybb1 |
| 3741.479534 | -0.797355546 | 0.207099545 | -3.850107672 | 0.000118066 | 0.016416971 | Down | ENSMUSG00000063576.12 | Klhdc3 |
| 2457.862788 | -1.214994046 | 0.315939949 | -3.845648674 | 0.000120234 | 0.016536565 | Down | ENSMUSG00000026820.5 | Ptges2 |
| 22.00440587 | 7.400455522 | 1.92612175 | 3.842153552 | 0.000121959 | 0.016638619 | Up | ENSMUSG00000071068.7 | Treml2 |
| 6500.348601 | 1.257801016 | 0.327775614 | 3.837384367 | 0.000124352 | 0.016829269 | Up | ENSMUSG00000069516.8 | Lyz2 |
| 8109.231001 | -1.080162469 | 0.281767654 | -3.833521888 | 0.000126322 | 0.016960173 | Down | ENSMUSG00000020219.7 | Timm13 |
| 215.0608707 | 1.903067404 | 0.497184227 | 3.827690622 | 0.000129351 | 0.016963056 | Up | ENSMUSG00000003154.15 | Foxj2 |
| 2318.753103 | -1.267169755 | 0.330855044 | -3.82998469 | 0.000128151 | 0.016963056 | Down | ENSMUSG00000005447.12 | Pafah1b3 |
| 799.9033012 | 2.416423245 | 0.631045253 | 3.829239238 | 0.00012854 | 0.016963056 | Up | ENSMUSG00000028268.14 | Gbp3 |
| 5295.715029 | -0.907881464 | 0.23777561 | -3.818227879 | 0.000134414 | 0.017144414 | Down | ENSMUSG00000007721.6 | Ccdc124 |
| 2418.693429 | -0.975914816 | 0.255931607 | -3.813185985 | 0.000137187 | 0.017144414 | Down | ENSMUSG00000011096.17 | Akt1s1 |
| 3115.213054 | -1.073558253 | 0.281086103 | -3.819321698 | 0.000133819 | 0.017144414 | Down | ENSMUSG00000020331.9 | Hcn2 |
| 381.750257 | 2.497846321 | 0.654946585 | 3.813816849 | 0.000136837 | 0.017144414 | Up | ENSMUSG00000024164.15 | C3 |
| 3767.662363 | -0.791901592 | 0.207658743 | -3.813475802 | 0.000137026 | 0.017144414 | Down | ENSMUSG00000027489.15 | Necab3 |
| 314.7450402 | -2.292403802 | 0.600509452 | -3.817431672 | 0.000134848 | 0.017144414 | Down | ENSMUSG00000035699.8 | Slc51a |
| 42.42372638 | 5.582837145 | 1.464529963 | 3.812033408 | 0.000137828 | 0.017144414 | Up | ENSMUSG00000112023.1 | Lilrb4b |
| 102.8751281 | 4.820653281 | 1.265962814 | 3.807894849 | 0.000140155 | 0.017150652 | Up | ENSMUSG00000026536.9 | Ifi211 |
| 3139.713823 | -1.016140347 | 0.266945349 | -3.806548235 | 0.00014092 | 0.017150652 | Down | ENSMUSG00000028959.14 | Fastk |
| 9474.453792 | -1.117955987 | 0.293452437 | -3.809666733 | 0.000139154 | 0.017150652 | Down | ENSMUSG00000059734.7 | Ndufs8 |
| 2979.709398 | -1.351910139 | 0.355629367 | -3.801458102 | 0.000143847 | 0.01732017 | Down | ENSMUSG00000046229.10 | Scand1 |
| 19510.6592 | -1.055039087 | 0.277599814 | -3.800575629 | 0.00014436 | 0.01732017 | Down | ENSMUSG00000068220.6 | Lgals1 |
| 1687.668391 | -1.247926884 | 0.328528215 | -3.798537928 | 0.000145552 | 0.017340182 | Down | ENSMUSG00000044927.6 | H1f10 |
| 3935.771597 | -1.176396539 | 0.310200242 | -3.792377883 | 0.000149212 | 0.01765184 | Down | ENSMUSG00000022577.17 | Ly6h |
| 1488.22578 | -1.228254371 | 0.324025135 | -3.790614486 | 0.000150275 | 0.017654187 | Down | ENSMUSG00000032583.7 | Mon1a |
| 1347.284948 | -1.260879483 | 0.332822041 | -3.788449465 | 0.00015159 | 0.017685897 | Down | ENSMUSG00000002804.4 | Nudt14 |
| 801.0599334 | 1.159659406 | 0.306891325 | 3.778729841 | 0.00015763 | 0.018017785 | Up | ENSMUSG00000006010.14 | Odr4 |
| 3527.391642 | -0.840010371 | 0.2222563 | -3.779467091 | 0.000157164 | 0.018017785 | Down | ENSMUSG00000030678.7 | Maz |
| 13498.60271 | -1.060062856 | 0.280375164 | -3.780872892 | 0.000156279 | 0.018017785 | Down | ENSMUSG00000069744.7 | Psmb3 |
| 1924.16016 | -1.209684321 | 0.320716389 | -3.771819476 | 0.000162061 | 0.018095763 | Down | ENSMUSG00000025226.11 | Fbxl15 |
| 2412.712373 | -1.058969933 | 0.280624643 | -3.77361703 | 0.000160898 | 0.018095763 | Down | ENSMUSG00000034793.15 | G6pc3 |
| 2147.908796 | -0.949251014 | 0.251799162 | -3.769873599 | 0.00016333 | 0.018095763 | Down | ENSMUSG00000042492.12 | Tbc1d10b |

| | | | | | | | | |
|---|---|---|---|---|---|---|---|---|
| 609.0765246 | -1.071467637 | 0.283809346 | -3.775307796 | 0.00015981 | 0.018095763 | Down | ENSMUSG00000049303.10 | Syt12 |
| 1008.06006 | 1.058398227 | 0.280789248 | 3.769368794 | 0.000163661 | 0.018095763 | Up | ENSMUSG00000050017.11 | Pitpnb |
| 4055.171709 | -0.900223691 | 0.2396551 | -3.756330208 | 0.000172423 | 0.018735666 | Down | ENSMUSG00000030284.11 | Creld1 |
| 3290.488469 | -0.908928242 | 0.242004896 | -3.755825835 | 0.000172771 | 0.018735666 | Down | ENSMUSG00000038276.12 | Asic3 |
| 573.8843873 | 2.240318184 | 0.596242795 | 3.757392463 | 0.000171693 | 0.018735666 | Up | ENSMUSG00000104713.4 | Gbp6 |
| 20608.22917 | -0.988646961 | 0.263615911 | -3.750331139 | 0.000176601 | 0.01902906 | Down | ENSMUSG00000006356.10 | Crip2 |
| 11577.31322 | -0.719509504 | 0.192056801 | -3.74633702 | 0.000179435 | 0.019201433 | Down | ENSMUSG00000013593.12 | Ndufs2 |
| 6685.367412 | -0.797762242 | 0.213026737 | -3.74489255 | 0.000180471 | 0.019201433 | Down | ENSMUSG00000055681.14 | Cope |
| 49159.0147 | -0.768218992 | 0.205251045 | -3.742826221 | 0.000181962 | 0.01921323 | Down | ENSMUSG00000023484.14 | Prph |
| 9863.441677 | -1.148926933 | 0.307068457 | -3.741598685 | 0.000182853 | 0.01921323 | Down | ENSMUSG00000039195.4 | Bbln |
| 835.558915 | -1.041514174 | 0.279131543 | -3.731266495 | 0.00019052 | 0.01930407 | Down | ENSMUSG00000004996.9 | Mri1 |
| 430.4307533 | 1.349112378 | 0.361764336 | 3.729257546 | 0.000192045 | 0.01930407 | Up | ENSMUSG00000008658.16 | Rbfox1 |
| 2380.826706 | -0.742837713 | 0.199209065 | -3.7289353 | 0.000192291 | 0.01930407 | Down | ENSMUSG00000024955.15 | Esrra |
| 16176.70118 | 0.652634135 | 0.174976871 | 3.729830874 | 0.000191608 | 0.01930407 | Up | ENSMUSG00000032399.8 | Rpl4 |
| 733.9711201 | -1.250874263 | 0.335348579 | -3.73007176 | 0.000191425 | 0.01930407 | Down | ENSMUSG00000041774.16 | Ydjc |
| 2921.787742 | -1.1483169 | 0.307582584 | -3.733361252 | 0.000188941 | 0.01930407 | Down | ENSMUSG00000051146.2 | Camk2n2 |
| 542.8932751 | -1.717101154 | 0.460570191 | -3.728207309 | 0.000192847 | 0.01930407 | Down | ENSMUSG00000091780.3 | Sco2 |
| 20.43612112 | 7.294063743 | 1.951334946 | 3.737986529 | 0.0001855 | 0.01930407 | Up | ENSMUSG00000097099.2 | Gm9917 |
| 471.0167755 | -1.208847149 | 0.324585307 | -3.724281792 | 0.000195872 | 0.019491573 | Down | ENSMUSG00000041046.7 | Ramp3 |
| 11395.90236 | -1.208512894 | 0.325145342 | -3.71683902 | 0.000201731 | 0.019841171 | Down | ENSMUSG00000020163.12 | Uqcr11 |
| 4832.215641 | -1.047503302 | 0.281791147 | -3.717303798 | 0.00020136 | 0.019841171 | Down | ENSMUSG00000028789.16 | Azin2 |
| 20.96772686 | 7.330908801 | 1.975755654 | 3.71043291 | 0.000206905 | 0.0201715 | Up | ENSMUSG00000056290.16 | Ms4a4b |
| 1043.682104 | -1.257430404 | 0.338954005 | -3.70973756 | 0.000207474 | 0.0201715 | Down | ENSMUSG00000104960.1 | Snhg8 |
| 5019.621707 | -0.958742087 | 0.258692358 | -3.706109048 | 0.000210468 | 0.020345613 | Down | ENSMUSG00000063802.5 | Hspbp1 |
| 11773.33768 | -0.973628595 | 0.263116257 | -3.700374142 | 0.000215282 | 0.020345937 | Down | ENSMUSG00000015094.16 | Npdc1 |
| 869.955809 | -1.06440425 | 0.287594404 | -3.701060371 | 0.0002147 | 0.020345937 | Down | ENSMUSG00000024889.5 | Rce1 |
| 4249.588542 | -0.988982654 | 0.267033685 | -3.703587629 | 0.000212572 | 0.020345937 | Down | ENSMUSG00000028070.7 | Naxe |
| 21.25921161 | -6.469735312 | 1.747907722 | -3.701416974 | 0.000214399 | 0.020345937 | Down | ENSMUSG00000108466.1 | Gm44771 |
| 4615.71779 | -1.004669942 | 0.271676142 | -3.698042577 | 0.000217268 | 0.020419613 | Down | ENSMUSG00000028670.14 | Lypla2 |
| 3527.666208 | -0.87435424 | 0.237314353 | -3.684371497 | 0.000229268 | 0.020821759 | Down | ENSMUSG00000013646.17 | Sh3bp5l |
| 28.47535563 | 7.772127742 | 2.106424149 | 3.689725901 | 0.000224496 | 0.020821759 | Up | ENSMUSG00000013974.3 | Mcemp1 |
| 2981.523327 | -0.89240197 | 0.242282318 | -3.68331448 | 0.000230221 | 0.020821759 | Down | ENSMUSG00000024799.16 | Tm7sf2 |
| 1506.144392 | -0.99100772 | 0.26910086 | -3.68266278 | 0.00023081 | 0.020821759 | Down | ENSMUSG00000027222.14 | Pex16 |
| 2682.726977 | -1.032696929 | 0.279757168 | -3.691404716 | 0.000223019 | 0.020821759 | Down | ENSMUSG00000038055.12 | Dexi |
| 137.7285594 | 2.950727075 | 0.801388273 | 3.682019283 | 0.000231394 | 0.020821759 | Up | ENSMUSG00000061808.4 | Ttr |
| 9480.872621 | -1.023201353 | 0.277887589 | -3.682069277 | 0.000231349 | 0.020821759 | Down | ENSMUSG00000071649.6 | B3gat3 |
| 19.81767788 | 7.249330122 | 1.968505409 | 3.682656948 | 0.000230816 | 0.020821759 | Up | ENSMUSG00000110390.1 | Gm45869 |
| 682.9301012 | -1.490040475 | 0.404854283 | -3.680436488 | 0.000232835 | 0.020840585 | Down | ENSMUSG00000022580.13 | Rhpn1 |
| 597.2090173 | 2.108520459 | 0.573569067 | 3.676140467 | 0.000236789 | 0.020972578 | Up | ENSMUSG00000044468.14 | Tent5c |
| 19.41817978 | 7.219820562 | 1.963502218 | 3.677011665 | 0.000235982 | 0.020972578 | Up | ENSMUSG00000073902.5 | Gvin3 |
| 7838.092532 | -0.745376439 | 0.20295824 | -3.67256062 | 0.000240132 | 0.021157895 | Down | ENSMUSG00000053565.10 | Eif3k |
| 46.85933867 | 7.451359238 | 2.033311731 | 3.664641838 | 0.000247685 | 0.021598383 | Up | ENSMUSG00000005800.3 | Mmp8 |
| 6009.791372 | -0.92063419 | 0.25115378 | -3.665619493 | 0.000246741 | 0.021598383 | Down | ENSMUSG00000006299.13 | Aamp |
| 2587.885311 | -1.32677487 | 0.3622589 | -3.662504555 | 0.000249761 | 0.021667754 | Down | ENSMUSG00000068327.5 | Tlx2 |
| 6073.426243 | -0.590573204 | 0.161727652 | -3.651652619 | 0.000260558 | 0.022489096 | Down | ENSMUSG00000037706.17 | Cd81 |
| 25467.90011 | -1.04137184 | 0.285440894 | -3.648292381 | 0.000263989 | 0.022669562 | Down | ENSMUSG00000027602.9 | Map1lc3a |
| 1700.146086 | -1.244170154 | 0.341521032 | -3.643026451 | 0.000269451 | 0.022702684 | Down | ENSMUSG00000002661.14 | Alkbh7 |
| 10072.0365 | -0.799765872 | 0.219588065 | -3.64211903 | 0.000270403 | 0.022702684 | Down | ENSMUSG00000005716.16 | Pvalb |
| 20213.75112 | -0.932692851 | 0.256130803 | -3.641470839 | 0.000271085 | 0.022702684 | Down | ENSMUSG00000031708.17 | Tecr |
| 398.8690999 | -1.241990028 | 0.340846627 | -3.643838401 | 0.000268602 | 0.022702684 | Down | ENSMUSG00000032291.8 | Crabp1 |
| 381.1143892 | -2.030076022 | 0.556841131 | -3.645700561 | 0.000266664 | 0.022702684 | Down | ENSMUSG00000109523.1 | Gdf1 |
| 340.8951343 | 1.8110482 | 0.497984946 | 3.636752908 | 0.000276097 | 0.023008499 | Up | ENSMUSG00000024659.15 | Anxa1 |
| 2298.23921 | -0.859954493 | 0.236585314 | -3.634859997 | 0.000278132 | 0.023064475 | Down | ENSMUSG00000029162.15 | Khk |
| 758.9056551 | -1.11297868 | 0.306431807 | -3.632059911 | 0.000281168 | 0.023202517 | Down | ENSMUSG00000049932.3 | H2ax |
| 510.37998 | 1.179267184 | 0.325027284 | 3.628209822 | 0.000285393 | 0.023285438 | Up | ENSMUSG00000014496.8 | Ankrd28 |
| 1293.987411 | -0.983376635 | 0.270921969 | -3.629741202 | 0.000283706 | 0.023285438 | Down | ENSMUSG00000030741.15 | Spns1 |
| 713.4627884 | -1.30440851 | 0.359722727 | -3.626149846 | 0.000287678 | 0.023285438 | Down | ENSMUSG00000037904.14 | Ankrd9 |
| 43.71384253 | 5.603884663 | 1.545003365 | 3.627101915 | 0.00028662 | 0.023285438 | Up | ENSMUSG00000054203.7 | Ifi205 |
| 21.39082239 | 7.359360339 | 2.032898574 | 3.620131587 | 0.000294453 | 0.023720311 | Up | ENSMUSG00000026532.7 | Spta1 |
| 426.2759063 | -1.284470398 | 0.3562024 | -3.606012754 | 0.000310938 | 0.024251063 | Down | ENSMUSG00000000214.11 | Th |
| 463.819852 | 1.192739678 | 0.330774747 | 3.605897032 | 0.000311076 | 0.024251063 | Up | ENSMUSG00000006494.11 | Pdk1 |
| 3008.468016 | -1.087629991 | 0.30130966 | -3.60967515 | 0.000306581 | 0.024251063 | Down | ENSMUSG00000024194.16 | Cuta |
| 1933.454822 | -0.920180284 | 0.254976816 | -3.608878246 | 0.000307524 | 0.024251063 | Down | ENSMUSG00000045176.3 | Borcs6 |
| 18425.62488 | -0.95054109 | 0.263272312 | -3.610486347 | 0.000305623 | 0.024251063 | Down | ENSMUSG00000071658.5 | Gng3 |
| 160618.1871 | -0.75158483 | 0.208360983 | -3.60712845 | 0.000309604 | 0.024251063 | Down | ENSMUSG00000072235.6 | Tuba1a |
| 33.34983323 | 8.000372283 | 2.215344384 | 3.611344737 | 0.000304613 | 0.024251063 | Up | ENSMUSG00000102206.5 | Pcdha11 |
| 18.03872143 | 7.113761372 | 1.974761776 | 3.602339005 | 0.000315367 | 0.024472747 | Up | ENSMUSG00000056130.10 | Ticam2 |
| 16604.36892 | -0.877159739 | 0.243621718 | -3.600498949 | 0.000317607 | 0.024493114 | Down | ENSMUSG00000002980.14 | Bcam |
| 150.5082055 | 3.27753707 | 0.910490522 | 3.599748696 | 0.000318525 | 0.024493114 | Up | ENSMUSG00000035042.2 | Ccl5 |
| 4442.497873 | -0.846175277 | 0.2353463 | -3.595447551 | 0.000323834 | 0.024788715 | Down | ENSMUSG00000059518.14 | Znhit1 |
| 4583.048341 | -0.789570231 | 0.220576573 | -3.579574289 | 0.000344154 | 0.026030539 | Down | ENSMUSG00000000743.9 | Chmp1a |
| 103298.7219 | -0.989230773 | 0.276384718 | -3.579180429 | 0.000344673 | 0.026030539 | Down | ENSMUSG00000027581.12 | Stmn3 |
| 1235.891509 | -0.923616269 | 0.257915663 | -3.58107863 | 0.000342179 | 0.026030539 | Down | ENSMUSG00000059540.15 | Tcea2 |
| 169.0027938 | 2.851847416 | 0.798513256 | 3.571446554 | 0.000355015 | 0.026401273 | Up | ENSMUSG00000029322.12 | Plac8 |
| 1154.22357 | 0.834374657 | 0.233732013 | 3.569791937 | 0.000357265 | 0.026401273 | Up | ENSMUSG00000030878.11 | Cdr2 |
| 6289.209672 | -0.816440371 | 0.228494273 | -3.573132762 | 0.000352736 | 0.026401273 | Down | ENSMUSG00000043670.4 | Diras1 |

| | | | | | | | | |
|---|---|---|---|---|---|---|---|---|
| 21.22990984 | 6.315413294 | 1.767172844 | 3.573738311 | 0.000351921 | 0.026401273 | Up | ENSMUSG00000044703.5 | Phf11a |
| 860.9074999 | 0.965451694 | 0.270457114 | 3.569703456 | 0.000357386 | 0.026401273 | Up | ENSMUSG00000067242.11 | Lgi1 |
| 6627.068206 | -0.89346146 | 0.2504713 | -3.5671211 | 0.000360925 | 0.026431876 | Down | ENSMUSG00000038502.17 | Ptov1 |
| 5782.596402 | -0.922809046 | 0.258628386 | -3.568088791 | 0.000359595 | 0.026431876 | Down | ENSMUSG00000044024.16 | Rell2 |
| 2132.89142 | -0.937391396 | 0.263027018 | -3.563859729 | 0.000365441 | 0.026647286 | Down | ENSMUSG00000024875.5 | Yif1a |
| 626.0549148 | -1.01686986 | 0.285519822 | -3.561468526 | 0.000368786 | 0.026775786 | Down | ENSMUSG00000016995.17 | Matn4 |
| 4522.728443 | -0.717483552 | 0.20152452 | -3.560279174 | 0.000370461 | 0.026782411 | Down | ENSMUSG00000036966.15 | Spryd3 |
| 20445.78102 | -1.038015833 | 0.291965893 | -3.555264021 | 0.0003776 | 0.026952972 | Down | ENSMUSG00000003072.15 | Atp5f1d |
| 3945.335679 | -0.943669252 | 0.265304659 | -3.556926798 | 0.000375219 | 0.026952972 | Down | ENSMUSG00000003531.15 | Dgcr6 |
| 1299.700588 | -1.070729837 | 0.301087861 | -3.556203942 | 0.000376252 | 0.026952972 | Down | ENSMUSG00000022561.13 | Gpaa1 |
| 11885.34693 | -1.143690119 | 0.321852561 | -3.553459741 | 0.000380199 | 0.027024508 | Down | ENSMUSG00000036578.7 | Fxyd7 |
| 81333.90259 | -0.844648823 | 0.238133606 | -3.546953478 | 0.000389713 | 0.02758486 | Down | ENSMUSG00000030695.16 | Aldoa |
| 1124.34479 | 1.804215353 | 0.509026626 | 3.544441998 | 0.000393445 | 0.027732959 | Up | ENSMUSG00000078853.8 | Igtp |
| 1261.708744 | -1.011251948 | 0.285772269 | -3.53866368 | 0.000402158 | 0.02822948 | Down | ENSMUSG00000007603.9 | Dus3l |
| 2404.590755 | -1.238781605 | 0.35044396 | -3.534892151 | 0.000407942 | 0.028407351 | Down | ENSMUSG00000023328.14 | Ache |
| 16.95869639 | 7.024935203 | 1.987352017 | 3.534821784 | 0.00040805 | 0.028407351 | Up | ENSMUSG00000108393.1 | Gm32633 |
| 10930.17861 | -0.767829946 | 0.217299071 | -3.533516935 | 0.00041007 | 0.028413054 | Down | ENSMUSG00000000308.14 | Ckmt1 |
| 39.93894032 | 4.438323218 | 1.256394777 | 3.532586491 | 0.000411516 | 0.028413054 | Up | ENSMUSG00000012519.14 | Mlkl |
| 917.7036925 | 1.014154493 | 0.287213761 | 3.531009415 | 0.000413977 | 0.028413054 | Up | ENSMUSG00000021156.17 | Zmynd11 |
| 460.9469491 | 1.116657952 | 0.316293205 | 3.530451917 | 0.00041485 | 0.028413054 | Up | ENSMUSG00000032411.15 | Tfdp2 |
| 2690.820426 | -0.976843114 | 0.277234897 | -3.523521472 | 0.000425853 | 0.028932322 | Down | ENSMUSG00000032997.16 | Chpf |
| 5945.877738 | -0.858058583 | 0.243465473 | -3.524354282 | 0.000424516 | 0.028932322 | Down | ENSMUSG00000039615.9 | Stub1 |
| 3710.091422 | -0.848471989 | 0.24092887 | -3.521670069 | 0.000428837 | 0.029018576 | Down | ENSMUSG00000087006.3 | Gm13889 |
| 12845.24375 | -0.743239655 | 0.211282157 | -3.517758745 | 0.000435208 | 0.029173216 | Down | ENSMUSG00000030120.14 | Mlf2 |
| 4005.842743 | -1.032294896 | 0.293507754 | -3.517095825 | 0.000436296 | 0.029173216 | Down | ENSMUSG00000040883.18 | Tmem205 |
| 2437.423908 | -1.254684493 | 0.356704978 | -3.517429163 | 0.000435749 | 0.029173216 | Down | ENSMUSG00000054716.4 | Zfp771 |
| 736.3247208 | -1.266668012 | 0.360583183 | -3.512831632 | 0.000443358 | 0.029184019 | Down | ENSMUSG00000029715.5 | Pop7 |
| 977.9481803 | -0.902012556 | 0.256681021 | -3.514138107 | 0.000441183 | 0.029184019 | Down | ENSMUSG00000047067.7 | Dusp28 |
| 24409.49213 | -0.844547985 | 0.240411767 | -3.512922826 | 0.000443206 | 0.029184019 | Down | ENSMUSG00000056596.8 | Trnp1 |
| 6984.555575 | -1.0294782 | 0.29298992 | -3.513698355 | 0.000441914 | 0.029184019 | Down | ENSMUSG00000073433.11 | Arhgdig |
| 498.4862154 | 1.058783599 | 0.301584171 | 3.510739956 | 0.000446861 | 0.029300592 | Up | ENSMUSG00000021876.15 | Rnase4 |
| 1486.928112 | 0.788830216 | 0.224839306 | 3.50841776 | 0.000450781 | 0.029443456 | Up | ENSMUSG00000029686.15 | Cul1 |
| 5187.790626 | -0.962224825 | 0.274585794 | -3.504277514 | 0.000457848 | 0.029790054 | Down | ENSMUSG00000006315.9 | Tmem147 |
| 526.1010527 | -1.51785721 | 0.433359942 | -3.502532338 | 0.000460858 | 0.029871004 | Down | ENSMUSG00000048772.15 | Tmem53 |
| 28355.51662 | -0.89136744 | 0.254675043 | -3.500018806 | 0.000465225 | 0.030029598 | Down | ENSMUSG00000019428.16 | Fkbp8 |
| 1011.172227 | -0.984789903 | 0.281441991 | -3.499086617 | 0.000466855 | 0.030029598 | Down | ENSMUSG00000029348.11 | Asphd2 |
| 2187.38828 | -0.812493867 | 0.232344655 | -3.496933752 | 0.000470639 | 0.03014242 | Down | ENSMUSG00000028857.16 | Tmem222 |
| 3464.728165 | -1.165705938 | 0.333433649 | -3.496065685 | 0.000472172 | 0.03014242 | Down | ENSMUSG00000057411.9 | Antkmt |
| 12032.16204 | -0.781541159 | 0.223971052 | -3.489473983 | 0.000483972 | 0.030779538 | Down | ENSMUSG00000026202.13 | Tuba4a |
| 6490.712219 | -0.883186933 | 0.253369129 | -3.485771673 | 0.00049072 | 0.031091777 | Down | ENSMUSG00000028072.6 | Ntrk1 |
| 16765.38762 | -1.067390644 | 0.307754043 | -3.468323706 | 0.000523716 | 0.032217105 | Down | ENSMUSG00000018286.6 | Psmb6 |
| 1281.942014 | -1.002648791 | 0.288943529 | -3.470051027 | 0.000520359 | 0.032217105 | Down | ENSMUSG00000022671.13 | Mzt2 |
| 3763.6933 | -0.627086042 | 0.180620718 | -3.471838942 | 0.000516906 | 0.032217105 | Down | ENSMUSG00000026424.8 | Gpr37l1 |
| 2257.682383 | 0.655993145 | 0.189071194 | 3.469556257 | 0.000521319 | 0.032217105 | Up | ENSMUSG00000027479.14 | Mapre1 |
| 3462.533656 | -0.780015722 | 0.224538357 | -3.473864029 | 0.000513021 | 0.032217105 | Down | ENSMUSG00000033735.9 | Spr |
| 4145.306124 | -1.097576781 | 0.316434877 | -3.468570816 | 0.000523235 | 0.032217105 | Down | ENSMUSG00000036186.5 | Dipk1b |
| 19.84300628 | 7.251144928 | 2.089757386 | 3.469850125 | 0.000520749 | 0.032217105 | Up | ENSMUSG00000058794.12 | Nfe2 |
| 6337.88659 | -0.780713665 | 0.2248581 | -3.472028196 | 0.000516542 | 0.032217105 | Down | ENSMUSG00000115987.1 | Vps28 |
| 22.65320612 | 6.404709807 | 1.847706104 | 3.466303323 | 0.000527668 | 0.032342584 | Up | ENSMUSG00000035910.15 | Dcdc2a |
| 12257.00584 | -1.102326066 | 0.318449942 | -3.461536402 | 0.000537101 | 0.032751238 | Down | ENSMUSG00000008036.11 | Ap2s1 |
| 1725.344023 | -1.111493566 | 0.321239406 | -3.460016255 | 0.000540143 | 0.032751238 | Down | ENSMUSG00000031158.11 | Timm17b |
| 491.4223684 | -1.245757026 | 0.359988358 | -3.460548096 | 0.000539077 | 0.032751238 | Down | ENSMUSG00000034936.2 | Arl4d |
| 7214.413232 | -1.001684193 | 0.289586111 | -3.459020153 | 0.000542144 | 0.032755199 | Down | ENSMUSG00000026688.5 | Mgst3 |
| 150670.4229 | -0.925076671 | 0.26770947 | -3.455524648 | 0.000549223 | 0.033064787 | Down | ENSMUSG00000014846.12 | Tppp3 |
| 20.21403867 | 6.240432394 | 1.808298387 | 3.450997046 | 0.00055852 | 0.033386846 | Up | ENSMUSG00000058488.7 | Kl |
| 1721.013284 | -0.869468551 | 0.251883745 | -3.451864478 | 0.000556727 | 0.033386846 | Down | ENSMUSG00000063954.7 | H2ac19 |
| 1192.616434 | -1.110128457 | 0.321893325 | -3.448746439 | 0.000563195 | 0.033411613 | Down | ENSMUSG00000002308.16 | Cd320 |
| 25706.96396 | -0.7870312 | 0.228281415 | -3.447635893 | 0.000565516 | 0.033411613 | Down | ENSMUSG00000023456.16 | Tpi1 |
| 1187.42154 | -0.8284949 | 0.240295461 | -3.447817522 | 0.000565136 | 0.033411613 | Down | ENSMUSG00000031787.8 | Katnb1 |
| 697.7045982 | 1.933605078 | 0.561104888 | 3.446067073 | 0.000568809 | 0.033411613 | Up | ENSMUSG00000046879.7 | Irgm1 |
| 144.1601259 | 2.485232064 | 0.721166392 | 3.446128511 | 0.00056868 | 0.033411613 | Up | ENSMUSG00000087107.8 | AI662270 |
| 8329.143516 | -1.14532139 | 0.332581429 | -3.443732243 | 0.000573744 | 0.033584846 | Down | ENSMUSG00000061111.8 | Mcrip1 |
| 2610.207031 | -0.709310871 | 0.206029554 | -3.442762731 | 0.000575804 | 0.033589243 | Down | ENSMUSG00000019579.13 | Mydgf |
| 5120.406312 | -0.719202483 | 0.209170051 | -3.438362618 | 0.000585243 | 0.034022553 | Down | ENSMUSG00000022199.12 | Slc22a17 |
| 994.4653154 | -0.900723206 | 0.262281443 | -3.434185787 | 0.000594337 | 0.034351115 | Down | ENSMUSG00000060402.8 | Chst8 |
| 36.72446543 | 5.342787879 | 1.555893506 | 3.433903322 | 0.000594956 | 0.034351115 | Up | ENSMUSG00000073400.6 | Trim10 |
| 971.8713114 | 0.894038477 | 0.260711863 | 3.429220544 | 0.000605317 | 0.03483046 | Up | ENSMUSG00000024601.9 | Isoc1 |
| 10988.87667 | -0.955163163 | 0.278755322 | -3.426528891 | 0.000611349 | 0.035058259 | Down | ENSMUSG00000056665.2 | Them6 |
| 20420.86967 | -0.997542774 | 0.291635259 | -3.42051499 | 0.000625027 | 0.035481813 | Down | ENSMUSG00000028843.8 | Sh3bgrl3 |
| 2257.503908 | -1.208147254 | 0.353040559 | -3.422120275 | 0.00062348 | 0.035481813 | Down | ENSMUSG00000044628.5 | Rnf208 |
| 44343.40868 | -0.861407433 | 0.251805229 | -3.420927501 | 0.00062408 | 0.035481813 | Down | ENSMUSG00000070493.3 | Chchd2 |
| 2009.173878 | -0.862455874 | 0.252440267 | -3.416475054 | 0.000634375 | 0.035892033 | Down | ENSMUSG00000005986.16 | Ankrd13d |
| 13176.80589 | -0.970536952 | 0.284339009 | -3.413309187 | 0.000641791 | 0.036190586 | Down | ENSMUSG00000075702.9 | Selenom |
| 30265.47156 | -0.742000483 | 0.217489304 | -3.411664254 | 0.000645676 | 0.0362887 | Down | ENSMUSG00000001794.13 | Capns1 |
| 14680.4122 | -0.770945824 | 0.226162017 | -3.408820964 | 0.000652443 | 0.036347441 | Down | ENSMUSG00000025510.14 | Cd151 |
| 2346.451734 | 0.849210684 | 0.249143639 | 3.40851842 | 0.000653167 | 0.036347441 | Up | ENSMUSG00000026229.17 | Psmd1 |

| | | | | | | | | |
|---|---|---|---|---|---|---|---|---|
| 3860.734301 | -0.668850546 | 0.196223824 | -3.408610293 | 0.000652947 | 0.036347441 | Down | ENSMUSG00000053398.11 | Phgdh |
| 1226.001618 | -1.108099144 | 0.325770907 | -3.401467472 | 0.000670251 | 0.037175859 | Down | ENSMUSG00000033256.14 | Shf |
| 21089.00049 | -0.750813666 | 0.220911034 | -3.398715095 | 0.000677032 | 0.037307332 | Down | ENSMUSG00000005161.15 | Prdx2 |
| 262.9129304 | 2.050812763 | 0.603322292 | 3.39919938 | 0.000675834 | 0.037307332 | Up | ENSMUSG00000049103.14 | Ccr2 |
| 989.8026149 | -0.99610323 | 0.293167367 | -3.39772888 | 0.000679477 | 0.037307622 | Down | ENSMUSG00000021018.8 | Polr2h |
| 738.1182541 | -0.939691206 | 0.276629023 | -3.396936426 | 0.000681448 | 0.037307622 | Down | ENSMUSG00000038517.15 | Tbkbp1 |
| 1130.74757 | 0.770190535 | 0.226864807 | 3.39493175 | 0.000686457 | 0.037460629 | Up | ENSMUSG00000026275.13 | Ppp1r7 |
| 18863.28776 | -0.933798095 | 0.275152653 | -3.393745568 | 0.000689437 | 0.037502277 | Down | ENSMUSG00000005779.12 | Psmb4 |
| 5601.563852 | -1.126396793 | 0.332319195 | -3.389502648 | 0.000700195 | 0.037965399 | Down | ENSMUSG00000019158.9 | Tmem160 |
| 3665.336369 | -1.122109632 | 0.331381962 | -3.386151815 | 0.000708802 | 0.03818725 | Down | ENSMUSG00000028445.7 | Enho |
| 5461.930657 | -1.050069814 | 0.310034531 | -3.386944706 | 0.000706756 | 0.03818725 | Down | ENSMUSG00000053291.15 | Rab4b |
| 16.90215748 | 7.02016097 | 2.075797734 | 3.38190993 | 0.000719837 | 0.038551746 | Up | ENSMUSG00000062939.11 | Stat4 |
| 21.32154433 | 6.31442319 | 1.867178039 | 3.381800266 | 0.000720125 | 0.038551746 | Up | ENSMUSG00000092021.8 | Gbp11 |
| 535.3370778 | -1.25590264 | 0.37155627 | -3.380114242 | 0.000724557 | 0.038658886 | Down | ENSMUSG00000071073.5 | Lrrc73 |
| 2203.466833 | -1.114572453 | 0.3298231 | -3.379303799 | 0.000726697 | 0.038658886 | Down | ENSMUSG00000078695.8 | Cisd3 |
| 602.3666487 | -0.889958233 | 0.263479514 | -3.377713199 | 0.000730913 | 0.038761285 | Down | ENSMUSG00000035206.10 | Sppl2b |
| 28.43181727 | -6.880525631 | 2.039462389 | -3.373695768 | 0.000741663 | 0.039086327 | Down | ENSMUSG00000036185.9 | Sapcd1 |
| 1819.65688 | -1.078112929 | 0.319561518 | -3.373725768 | 0.000741582 | 0.039086327 | Down | ENSMUSG00000061286.7 | Exosc5 |
| 483.058496 | -1.123702186 | 0.333200184 | -3.372453682 | 0.000745016 | 0.039141113 | Down | ENSMUSG00000055629.5 | B4galnt4 |
| 18.39166354 | -8.186204945 | 2.428575019 | -3.370785288 | 0.000749543 | 0.039257 | Down | ENSMUSG00000084771.8 | A230072E10Rik |
| 260.9812394 | 1.345083442 | 0.399761996 | 3.364710637 | 0.00076624 | 0.040007646 | Up | ENSMUSG00000042487.6 | Leo1 |
| 3745.467718 | -0.822989563 | 0.244940245 | -3.359960557 | 0.000779536 | 0.04045218 | Down | ENSMUSG00000036622.15 | Atp13a2 |
| 7264.551053 | -0.955304161 | 0.28425373 | -3.360744507 | 0.000777327 | 0.04045218 | Down | ENSMUSG00000067925.4 | Rtl8a |
| 1041.971192 | -1.122060646 | 0.33424158 | -3.35703489 | 0.000787832 | 0.040633381 | Down | ENSMUSG00000021773.11 | Comtd1 |
| 4873.300921 | -1.035720985 | 0.308469508 | -3.357612204 | 0.000786188 | 0.040633381 | Down | ENSMUSG00000110841.1 | Gpx4-ps2 |
| 101571.6026 | -0.661684613 | 0.197306179 | -3.353592962 | 0.000797696 | 0.040741512 | Down | ENSMUSG00000023004.8 | Tuba1b |
| 3304.44648 | -0.949399782 | 0.283135152 | -3.353168185 | 0.000798921 | 0.040741512 | Down | ENSMUSG00000039183.6 | Nubp2 |
| 4622.225788 | -0.831324564 | 0.247938509 | -3.352946532 | 0.000799562 | 0.040741512 | Down | ENSMUSG00000071076.6 | Jund |
| 866.8262742 | -0.912838302 | 0.272117231 | -3.354577357 | 0.000794863 | 0.040741512 | Down | ENSMUSG00000078794.4 | Dact3 |
| 20321.89208 | -0.839925711 | 0.250590594 | -3.351784674 | 0.000802924 | 0.040790007 | Down | ENSMUSG00000022658.10 | Tagln3 |
| 546.7312954 | -1.152170176 | 0.344156817 | -3.347805759 | 0.000814541 | 0.041149727 | Down | ENSMUSG00000024873.7 | Cnih2 |
| 5481.029681 | -0.925248325 | 0.276383805 | -3.347693715 | 0.00081487 | 0.041149727 | Down | ENSMUSG00000037499.9 | Nenf |
| 6931.765128 | -0.858831914 | 0.256914323 | -3.34287284 | 0.000829159 | 0.041703284 | Down | ENSMUSG00000024767.11 | Otub1 |
| 951.3521421 | 0.813505807 | 0.243394335 | 3.342336657 | 0.000830762 | 0.041703284 | Up | ENSMUSG00000041133.11 | Smc1a |
| 1460.917111 | -1.016707465 | 0.30435985 | -3.340478272 | 0.000836342 | 0.041859177 | Down | ENSMUSG00000079478.9 | Znrd2 |
| 5803.261482 | -0.90552342 | 0.27120867 | -3.338843924 | 0.000841278 | 0.041982014 | Down | ENSMUSG00000058833.11 | Rex1bd |
| 1478.658615 | -1.103122456 | 0.33055673 | -3.337165318 | 0.000846376 | 0.042112179 | Down | ENSMUSG00000003872.9 | Lin7b |
| 5450.117497 | 0.639834409 | 0.191784142 | 3.336221664 | 0.000849254 | 0.042131476 | Up | ENSMUSG00000020719.14 | Ddx5 |
| 17.53911666 | -6.177166159 | 1.852250204 | -3.334952344 | 0.00085314 | 0.042200505 | Down | ENSMUSG00000106825.1 | 2510016D11Rik |
| 353.0166549 | -1.140488245 | 0.34222664 | -3.332552498 | 0.000860532 | 0.042442057 | Down | ENSMUSG00000036957.8 | Lrfn3 |
| 5204.769038 | -0.777982882 | 0.233684899 | -3.329196223 | 0.00087097 | 0.042461679 | Down | ENSMUSG00000028779.16 | Pef1 |
| 664.6326601 | 0.872130063 | 0.26184422 | 3.33072108 | 0.000866213 | 0.042461679 | Up | ENSMUSG00000032497.19 | Lrrfip2 |
| 1467.38304 | -0.747956637 | 0.224629579 | -3.329733516 | 0.000869291 | 0.042461679 | Down | ENSMUSG00000035781.14 | R3hdm4 |
| 2196.776706 | -0.968452799 | 0.290834415 | -3.329911278 | 0.000868737 | 0.042461679 | Down | ENSMUSG00000044709.6 | Gemin7 |
| 3230.152947 | -0.759817255 | 0.228479287 | -3.325541084 | 0.000882471 | 0.042898738 | Down | ENSMUSG00000029001.15 | Fbxo44 |
| 56767.21397 | -0.853715152 | 0.256832716 | -3.324012481 | 0.000887322 | 0.043010976 | Down | ENSMUSG00000031818.12 | Cox4i1 |
| 352.1018113 | -1.633890603 | 0.491879775 | -3.321727556 | 0.00089462 | 0.043240819 | Down | ENSMUSG00000050212.4 | Eva1b |
| 923.7220142 | -1.062999817 | 0.320430319 | -3.317413353 | 0.000908551 | 0.043789048 | Down | ENSMUSG00000064264.14 | Zfp428 |
| 5539.757228 | -0.973936744 | 0.293939958 | -3.313386686 | 0.000921734 | 0.044298245 | Down | ENSMUSG00000030401.16 | Rtn2 |
| 9095.066658 | -0.726188678 | 0.219245932 | -3.312210496 | 0.000925619 | 0.044358901 | Down | ENSMUSG00000024914.16 | Drap1 |
| 260.3767199 | 1.51180551 | 0.457094202 | 3.307426572 | 0.000941574 | 0.044869313 | Up | ENSMUSG00000026896.14 | Ifih1 |
| 39.05379838 | 4.966338272 | 1.501252401 | 3.308130111 | 0.000939212 | 0.044869313 | Up | ENSMUSG00000078763.2 | Slfn1 |
| 3010.42439 | -0.892851886 | 0.270071416 | -3.305984397 | 0.000946434 | 0.044974205 | Down | ENSMUSG00000040813.16 | Tex264 |
| 1895.758525 | -1.046197741 | 0.317094027 | -3.299329702 | 0.00096916 | 0.045925158 | Down | ENSMUSG00000038520.15 | Tbc1d17 |
| 766.8245298 | 0.8974669 | 0.272348076 | 3.2952937 | 0.000983189 | 0.046459783 | Up | ENSMUSG00000020171.8 | Yeats4 |
| 61.77096716 | 4.364373512 | 1.325801118 | 3.291876475 | 0.000995213 | 0.046756268 | Up | ENSMUSG00000027398.13 | Il1b |
| 2729.607332 | -0.805306268 | 0.244667946 | -3.291425308 | 0.000996811 | 0.046756268 | Down | ENSMUSG00000028763.18 | Hspg2 |
| 1958.331843 | -0.810682106 | 0.24632115 | -3.291159149 | 0.000997754 | 0.046756268 | Down | ENSMUSG00000035640.18 | Cbarp |
| 3618.209591 | -0.905946376 | 0.275402036 | -3.289541311 | 0.001003508 | 0.046766802 | Down | ENSMUSG00000020477.10 | Mrps24 |
| 1563.276851 | -0.926935843 | 0.281719423 | -3.290280214 | 0.001000877 | 0.046766802 | Down | ENSMUSG00000026857.9 | Ntmt1 |
| 597.5804188 | -1.061835681 | 0.323228277 | -3.285095258 | 0.001019479 | 0.047380566 | Down | ENSMUSG00000021265.3 | Slc25a29 |
| 724.3923144 | -1.43987952 | 0.438545066 | -3.28331027 | 0.001025957 | 0.047550989 | Down | ENSMUSG00000044287.6 | Nrn1l |
| 2287.115265 | -0.828110039 | 0.25237829 | -3.28122534 | 0.001033571 | 0.047773023 | Down | ENSMUSG00000040610.7 | Tlx3 |
| 1116.402638 | -1.20117463 | 0.366584441 | -3.276665605 | 0.001050407 | 0.048418894 | Down | ENSMUSG00000044876.15 | Zfp444 |
| 1152.634166 | 0.913342496 | 0.278983362 | 3.273824255 | 0.001061026 | 0.048586254 | Up | ENSMUSG00000026353.9 | Ubxn4 |
| 6237.782082 | -0.826638604 | 0.252474532 | -3.274146493 | 0.001059817 | 0.048586254 | Down | ENSMUSG00000030688.15 | Stard10 |
| 758.2163815 | 1.500606176 | 0.458425587 | 3.27339097 | 0.001062654 | 0.048586254 | Up | ENSMUSG00000035692.7 | Isg15 |
| 6938.604396 | -0.746486635 | 0.228634529 | -3.264977688 | 0.001094727 | 0.048935173 | Down | ENSMUSG00000019087.13 | Atp6ap1 |
| 1621.301336 | -0.968754434 | 0.296479419 | -3.267526752 | 0.001084916 | 0.048935173 | Down | ENSMUSG00000022557.11 | Bop1 |
| 19868.27506 | -0.72674146 | 0.222336674 | -3.26865311 | 0.001080607 | 0.048935173 | Down | ENSMUSG00000028937.14 | Acot7 |
| 6318.486425 | 0.635746812 | 0.194781516 | 3.263897035 | 0.001098911 | 0.048935173 | Up | ENSMUSG00000029622.16 | Arpc1b |
| 2822.950892 | -0.928167931 | 0.283943124 | -3.268851582 | 0.001079849 | 0.048935173 | Down | ENSMUSG00000033152.13 | Podxl2 |
| 400.9873073 | 1.592572742 | 0.487448588 | 3.26716044 | 0.001086321 | 0.048935173 | Up | ENSMUSG00000034248.7 | Slc25a37 |
| 1910.078165 | -0.881485213 | 0.269918179 | -3.265749706 | 0.001091747 | 0.048935173 | Down | ENSMUSG00000035828.11 | Pim3 |
| 577.5335055 | -1.18701449 | 0.363688769 | -3.263819478 | 0.001099212 | 0.048935173 | Down | ENSMUSG00000037349.9 | Nudt22 |
| 9171.633957 | -0.958941849 | 0.293619254 | -3.265936537 | 0.001091027 | 0.048935173 | Down | ENSMUSG00000040907.15 | Atp1a3 |

| | | | | | | | | |
|---|---|---|---|---|---|---|---|---|
| 24.49557274 | 5.409606072 | 1.65602731 | 3.266616461 | 0.00108841 | 0.048935173 | Up | ENSMUSG00000060336.6 | Zfp937 |
| 4540.264895 | -0.782282691 | 0.239799218 | -3.262240382 | 0.001105354 | 0.049079445 | Down | ENSMUSG00000031820.9 | Babam1 |
| 3887.244915 | -0.880528801 | 0.270071467 | -3.260354787 | 0.001112729 | 0.049102186 | Down | ENSMUSG00000023011.8 | Faim2 |
| 454.1801809 | -1.238675436 | 0.380024437 | -3.259462594 | 0.001116235 | 0.049102186 | Down | ENSMUSG00000044216.7 | Kcnj4 |
| 4671.83183 | -0.816337469 | 0.250376918 | -3.260434208 | 0.001112418 | 0.049102186 | Down | ENSMUSG00000059316.2 | Slc27a4 |
| 1890.767888 | -0.817843901 | 0.250937993 | -3.259147381 | 0.001117476 | 0.049102186 | Down | ENSMUSG00000083282.3 | Ctsf |
| 2734.24575 | -0.999372934 | 0.307050924 | -3.254746542 | 0.001134936 | 0.049740189 | Down | ENSMUSG00000035754.8 | Wdr18 |
| 54.99994578 | 4.581906385 | 1.409367188 | 3.251038071 | 0.001149845 | 0.050263362 | Up | ENSMUSG00000044827.10 | Tlr1 |
| 1617.024368 | -1.04272609 | 0.320953107 | -3.24884249 | 0.001158756 | 0.050522368 | Down | ENSMUSG00000035559.9 | Mpv17l2 |
| 7383.870598 | -0.936475442 | 0.288812028 | -3.242508455 | 0.001184824 | 0.051496017 | Down | ENSMUSG00000023020.3 | Cox14 |
| 9292.234232 | -0.814426236 | 0.251215429 | -3.241943535 | 0.001187175 | 0.051496017 | Down | ENSMUSG00000024847.15 | Aip |
| 75243.13303 | 0.502479896 | 0.155042288 | 3.24092158 | 0.00119144 | 0.051548805 | Up | ENSMUSG00000037742.14 | Eef1a1 |
| 2522.770105 | -0.980536884 | 0.302716 | -3.239131341 | 0.001198943 | 0.051582483 | Down | ENSMUSG00000024925.11 | Rnaseh2c |
| 1024.925256 | -1.133256374 | 0.350082626 | -3.237111157 | 0.001207464 | 0.051582483 | Down | ENSMUSG00000029029.14 | Wrap73 |
| 129.934475 | 7.092841468 | 2.190975539 | 3.237298336 | 0.001206672 | 0.051582483 | Up | ENSMUSG00000040809.10 | Chil3 |
| 1490.08995 | -0.987217681 | 0.30492162 | -3.237611296 | 0.001205349 | 0.051582483 | Down | ENSMUSG00000045777.14 | Ifitm10 |
| 3287.113086 | -0.803890962 | 0.248335855 | -3.237111946 | 0.00120746 | 0.051582483 | Down | ENSMUSG00000071657.12 | Bscl2 |
| 2293.384665 | -1.050662796 | 0.32538875 | -3.228946288 | 0.001242472 | 0.052944342 | Down | ENSMUSG00000038880.13 | Mrps34 |
| 184.1564971 | 1.589269052 | 0.492475705 | 3.227101433 | 0.001250511 | 0.053019795 | Up | ENSMUSG00000031684.11 | Slc10a7 |
| 43576.27064 | -0.877423597 | 0.271887995 | -3.227150935 | 0.001250295 | 0.053019795 | Down | ENSMUSG00000041571.9 | Selenow |
| 4991.271002 | -0.816193847 | 0.253047926 | -3.22545163 | 0.001257741 | 0.053193006 | Down | ENSMUSG00000033423.16 | Eri3 |
| 979.9759374 | -0.851189165 | 0.2640897 | -3.223106263 | 0.001268085 | 0.053444058 | Down | ENSMUSG00000020388.12 | Pdlim4 |
| 970.8467709 | -1.06306565 | 0.330011658 | -3.221297257 | 0.001276117 | 0.053444058 | Down | ENSMUSG00000028958.15 | Tmub1 |
| 408.338725 | -1.260571427 | 0.391329518 | -3.221253113 | 0.001276314 | 0.053444058 | Down | ENSMUSG00000040904.10 | Gm21988 |
| 560.477752 | 1.038108929 | 0.322140534 | 3.222534328 | 0.001270619 | 0.053444058 | Up | ENSMUSG00000068794.7 | Col28a1 |
| 1726.81385 | -0.920926201 | 0.286186192 | -3.217926747 | 0.001291208 | 0.053801389 | Down | ENSMUSG00000037428.14 | Vgf |
| 29178.73885 | -0.837796326 | 0.260301512 | -3.218561122 | 0.001288355 | 0.053801389 | Down | ENSMUSG00000059412.7 | Fxyd2 |
| 2278.564703 | -0.900909248 | 0.28015772 | -3.215721655 | 0.00130117 | 0.053818796 | Down | ENSMUSG00000003873.11 | Bax |
| 1873.274805 | -0.782589781 | 0.24332094 | -3.216286191 | 0.001298613 | 0.053818796 | Down | ENSMUSG00000024797.14 | Vps51 |
| 1977.744284 | -0.935770963 | 0.290881302 | -3.217019988 | 0.001295296 | 0.053818796 | Down | ENSMUSG00000031622.16 | Sin3b |
| 14.29265264 | 6.778110406 | 2.110370505 | 3.211810623 | 0.001319013 | 0.05442376 | Up | ENSMUSG00000094595.1 | Fsbp |
| 3999.701891 | -1.028593701 | 0.320342964 | -3.210913979 | 0.001323135 | 0.054461024 | Down | ENSMUSG00000031813.8 | Mvb12a |
| 1157.275662 | -0.777538698 | 0.242278705 | -3.209273793 | 0.001330707 | 0.054527504 | Down | ENSMUSG00000010609.15 | Psen2 |
| 54943.36601 | -0.769344247 | 0.239733243 | -3.209167975 | 0.001331197 | 0.054527504 | Down | ENSMUSG00000075706.10 | Gpx4 |
| 4608.193926 | 0.626461574 | 0.195419684 | 3.205724014 | 0.001347231 | 0.055050997 | Up | ENSMUSG00000027012.15 | Dync1i2 |
| 548.6055882 | 0.96965944 | 0.302705204 | 3.203312752 | 0.001358564 | 0.055380288 | Up | ENSMUSG00000032547.12 | Ryk |
| 27198.60627 | -0.843555464 | 0.263438492 | -3.202096464 | 0.001364313 | 0.055465358 | Down | ENSMUSG00000035242.15 | Oaz1 |
| 689.5848356 | -1.260674775 | 0.393778022 | -3.201485874 | 0.001367208 | 0.055465358 | Down | ENSMUSG00000039450.11 | Dcxr |
| 102.2881467 | 1.89639525 | 0.592796015 | 3.199068822 | 0.001378723 | 0.055798682 | Up | ENSMUSG00000022762.18 | Ncam2 |
| 654.0361079 | 0.953848091 | 0.298556035 | 3.19487125 | 0.001398932 | 0.056481475 | Up | ENSMUSG00000021000.17 | Mia2 |
| 1435.735634 | -1.030199127 | 0.322582064 | -3.193603242 | 0.001405091 | 0.056595052 | Down | ENSMUSG00000066235.7 | Pomgnt2 |
| 4573.884276 | -1.007246423 | 0.315846773 | -3.189035025 | 0.001427486 | 0.057360516 | Down | ENSMUSG00000042380.8 | Smim12 |
| 115.833296 | 2.902161059 | 0.911517061 | 3.18388013 | 0.001453152 | 0.05792556 | Up | ENSMUSG00000023132.8 | Gzma |
| 150.6392518 | 1.93805361 | 0.609108377 | 3.181787814 | 0.00146369 | 0.05792556 | Up | ENSMUSG00000026471.14 | Mr1 |
| 3174.451041 | -0.728446892 | 0.228968628 | -3.181426641 | 0.001465516 | 0.05792556 | Down | ENSMUSG00000027603.15 | Ggt7 |
| 2232.361196 | -0.847999815 | 0.266409438 | -3.183069721 | 0.001457225 | 0.05792556 | Down | ENSMUSG00000033020.7 | Polr2f |
| 6174.798319 | -0.700665817 | 0.220009796 | -3.184702814 | 0.001449027 | 0.05792556 | Down | ENSMUSG00000034993.7 | Vat1 |
| 2381.986871 | -0.928630139 | 0.291822265 | -3.182177139 | 0.001461724 | 0.05792556 | Down | ENSMUSG00000042532.14 | Golga7b |
| 965.9173837 | 1.030124172 | 0.323731754 | 3.182030061 | 0.001462466 | 0.05792556 | Up | ENSMUSG00000053279.8 | Aldh1a1 |
| 4629.491609 | -0.950945027 | 0.299173664 | -3.178571989 | 0.001480025 | 0.058100647 | Down | ENSMUSG00000024181.9 | Mrpl28 |
| 21705.5268 | -0.7529344 | 0.236880267 | -3.178544198 | 0.001480166 | 0.058100647 | Down | ENSMUSG00000030122.12 | Ptms |
| 744.39461 | -0.855807821 | 0.269272825 | -3.1782183 | 0.001481831 | 0.058100647 | Down | ENSMUSG00000049482.16 | Ctu2 |
| 2118.747512 | -1.204155931 | 0.378920853 | -3.177856069 | 0.001483684 | 0.058100647 | Down | ENSMUSG00000070858.12 | Nicol1 |
| 1434.801322 | -1.084663374 | 0.34152745 | -3.175918576 | 0.001493629 | 0.058355003 | Down | ENSMUSG00000074738.2 | Fndc10 |
| 1313.415213 | -0.90141757 | 0.28429185 | -3.17074714 | 0.001520474 | 0.059266968 | Down | ENSMUSG00000029060.17 | Mib2 |
| 533.4705417 | -0.997491898 | 0.31495045 | -3.167139146 | 0.001539466 | 0.059869317 | Down | ENSMUSG00000020829.9 | Slc46a1 |
| 2577.760097 | -0.8149184 | 0.257551512 | -3.164098686 | 0.00155564 | 0.06035956 | Down | ENSMUSG00000013858.14 | Tmem259 |
| 2196.099933 | 0.676842416 | 0.214008655 | 3.162687112 | 0.001563202 | 0.060514177 | Up | ENSMUSG00000019699.16 | Akt3 |
| 2342.268331 | -0.827541617 | 0.261828576 | -3.160623754 | 0.001574317 | 0.060666794 | Down | ENSMUSG00000075467.4 | Dnlz |
| 2282.622202 | -0.981255016 | 0.310430323 | -3.160950927 | 0.00157255 | 0.060666794 | Down | ENSMUSG00000096606.2 | Tpbgl |
| 11342.5846 | -0.85212897 | 0.26998685 | -3.156186937 | 0.001598464 | 0.061427854 | Down | ENSMUSG00000022551.8 | Cyc1 |
| 568.5437132 | 2.048378445 | 0.649111591 | 3.155664563 | 0.001601329 | 0.061427854 | Up | ENSMUSG00000078920.3 | Ifi47 |
| 3657.69921 | -0.854934397 | 0.27108809 | -3.153714342 | 0.001612068 | 0.061699896 | Down | ENSMUSG00000034891.13 | Sncb |
| 48792.79303 | -0.635312335 | 0.201500826 | -3.152901889 | 0.001616561 | 0.061732206 | Down | ENSMUSG00000032294.17 | Pkm |
| 10780.12959 | -0.862464066 | 0.273817867 | -3.149772786 | 0.001633975 | 0.062116748 | Down | ENSMUSG00000026860.16 | Sh3glb2 |
| 858.1701127 | -0.910963985 | 0.289198992 | -3.149955597 | 0.001632953 | 0.062116748 | Down | ENSMUSG00000078317.6 | F8a |
| 5300.341808 | -0.794279625 | 0.25222911 | -3.149040275 | 0.001638076 | 0.062133037 | Down | ENSMUSG00000024939.6 | Fam89b |
| 5384.72645 | -0.675374464 | 0.215001712 | -3.141251559 | 0.001682275 | 0.063666758 | Down | ENSMUSG00000053929.17 | Zftraf1 |
| 23036.90555 | -0.755438596 | 0.240741115 | -3.137970829 | 0.001701218 | 0.063954458 | Down | ENSMUSG00000004267.16 | Eno2 |
| 445.6567986 | 0.988881853 | 0.315103684 | 3.138274492 | 0.001699456 | 0.063954458 | Up | ENSMUSG00000031015.8 | Swap70 |
| 14.27976394 | 6.776702949 | 2.158754551 | 3.139172513 | 0.001694257 | 0.063954458 | Up | ENSMUSG00000032739.16 | Pram1 |
| 5498.339109 | -0.981998329 | 0.313279682 | -3.134573944 | 0.001721039 | 0.064556125 | Down | ENSMUSG00000027613.15 | Eif6 |
| 6486.763146 | -0.698527387 | 0.222987256 | -3.132588829 | 0.00173272 | 0.064850491 | Down | ENSMUSG00000008690.15 | Ncaph2 |
| 968.463409 | -0.791098353 | 0.252804495 | -3.129289111 | 0.001752298 | 0.065438468 | Down | ENSMUSG00000004748.5 | Mtfp1 |
| 5781.14704 | -0.865241748 | 0.276655082 | -3.127510769 | 0.001762934 | 0.065690634 | Down | ENSMUSG00000067924.4 | Rtl8b |
| 962.5529822 | -0.927634076 | 0.296747398 | -3.126005762 | 0.001771981 | 0.065882635 | Down | ENSMUSG00000078636.4 | Gm7336 |

| | | | | | | | | |
|---|---|---|---|---|---|---|---|---|
| 622.5063414 | -1.030702473 | 0.329982458 | -3.123506861 | 0.001787097 | 0.066019253 | Down | ENSMUSG00000036114.3 | Rpp25l |
| 1088.746857 | -0.861792098 | 0.275890822 | -3.123670778 | 0.001786102 | 0.066019253 | Down | ENSMUSG00000042729.11 | Wdr74 |
| 4164.969679 | -0.79712121 | 0.2552043 | -3.123463081 | 0.001787363 | 0.066019253 | Down | ENSMUSG00000061046.9 | Haghl |
| 5608.747258 | -0.954005666 | 0.305555912 | -3.122196721 | 0.001795069 | 0.066159456 | Down | ENSMUSG00000023904.10 | Hcfc1r1 |
| 801.7166718 | 0.885012331 | 0.283582089 | 3.120832964 | 0.001803403 | 0.066178238 | Up | ENSMUSG00000011958.17 | Bnip2 |
| 2564.980584 | -0.733609371 | 0.235064368 | -3.12088717 | 0.001803071 | 0.066178238 | Down | ENSMUSG00000027466.15 | Rbck1 |
| 4058.346643 | 1.031931803 | 0.330807217 | 3.119435581 | 0.001811979 | 0.066349015 | Up | ENSMUSG00000027078.14 | Ube2l6 |
| 3112.550141 | -0.963146005 | 0.309006955 | -3.116907204 | 0.001827591 | 0.066488936 | Down | ENSMUSG00000026209.15 | Dnpep |
| 10350.56743 | -0.714948607 | 0.22930948 | -3.117832752 | 0.001821862 | 0.066488936 | Down | ENSMUSG00000039347.7 | Atp6v0e2 |
| 191.2380723 | 2.093467472 | 0.671636605 | 3.116964525 | 0.001827236 | 0.066488936 | Up | ENSMUSG00000053063.11 | Clec12a |
| 33413.98307 | -0.691702278 | 0.222011427 | -3.115615653 | 0.001835614 | 0.066637498 | Down | ENSMUSG00000032959.12 | Pebp1 |
| 7081.02925 | -0.79275586 | 0.254562396 | -3.114190754 | 0.001844502 | 0.066816788 | Down | ENSMUSG00000003199.16 | Mpnd |
| 134576.683 | -0.644678369 | 0.207096214 | -3.112941355 | 0.001852328 | 0.066956915 | Down | ENSMUSG00000002985.16 | Apoe |
| 723.4692862 | -0.919913793 | 0.296305018 | -3.104617665 | 0.001905251 | 0.068264021 | Down | ENSMUSG00000000792.2 | Slc5a5 |
| 369.0511005 | -1.07446542 | 0.346027861 | -3.105141348 | 0.001901881 | 0.068264021 | Down | ENSMUSG00000007837.11 | Prrg2 |
| 1180.599887 | -0.843974453 | 0.271691343 | -3.106372262 | 0.001893981 | 0.068264021 | Down | ENSMUSG00000019470.12 | Xab2 |
| 1420.31497 | 0.829345737 | 0.267102937 | 3.104966748 | 0.001903004 | 0.068264021 | Up | ENSMUSG00000033186.9 | Mzt1 |
| 2635.497925 | -0.835230295 | 0.269098176 | -3.103812549 | 0.001910443 | 0.068264021 | Down | ENSMUSG00000042078.14 | Svop |
| 2580.43646 | -0.609806723 | 0.19649233 | -3.103463232 | 0.0019127 | 0.068264021 | Down | ENSMUSG00000072694.8 | 1500011B03Rik |
| 2210.362739 | 0.631779733 | 0.203622445 | 3.102701831 | 0.001917627 | 0.068295795 | Up | ENSMUSG00000030616.16 | Sytl2 |
| 3027.697562 | -0.811824018 | 0.261792727 | -3.101018226 | 0.001928564 | 0.068397318 | Down | ENSMUSG00000000753.15 | Serpinf1 |
| 4018.526478 | -0.830711883 | 0.267862893 | -3.101257789 | 0.001927004 | 0.068397318 | Down | ENSMUSG00000035203.16 | Epn1 |
| 808.6324359 | -1.307755641 | 0.421889582 | -3.099758085 | 0.001936788 | 0.068422686 | Down | ENSMUSG00000041199.3 | Rpusd1 |
| 955.9618158 | -1.057036846 | 0.341016015 | -3.099669222 | 0.001937369 | 0.068422686 | Down | ENSMUSG00000044991.10 | Shld1 |
| 2069.933577 | 1.231081704 | 0.397258019 | 3.098947393 | 0.001942095 | 0.068446711 | Up | ENSMUSG00000026104.14 | Stat1 |
| 8952.3731 | -0.819465892 | 0.264488613 | -3.098303108 | 0.001946323 | 0.068453093 | Down | ENSMUSG00000037916.14 | Ndufv1 |
| 1604.724537 | -0.782834849 | 0.25287549 | -3.095732403 | 0.001963275 | 0.068590137 | Down | ENSMUSG00000002767.13 | Mrpl2 |
| 2053.424248 | -0.854423969 | 0.275898758 | -3.096875012 | 0.001955723 | 0.068590137 | Down | ENSMUSG00000003423.15 | Pih1d1 |
| 856.6101566 | 1.021544362 | 0.330033997 | 3.095270092 | 0.001966337 | 0.068590137 | Up | ENSMUSG00000026023.16 | Cdk15 |
| 4908.537036 | -0.717867202 | 0.231925055 | -3.095255069 | 0.001966437 | 0.068590137 | Down | ENSMUSG00000033475.15 | Tomm6 |
| 1531.431023 | -0.815699086 | 0.263808669 | -3.092010167 | 0.00198806 | 0.069201683 | Down | ENSMUSG00000006442.10 | Srm |
| 552.7878823 | -0.986201571 | 0.31926325 | -3.088991832 | 0.00200837 | 0.069765069 | Down | ENSMUSG00000015377.10 | Dennd6b |
| 300.3189071 | 1.191861005 | 0.385930577 | 3.088278243 | 0.002013199 | 0.069789517 | Up | ENSMUSG00000063785.12 | Utp14a |
| 36.61041988 | 4.508464355 | 1.460309333 | 3.087335165 | 0.002019597 | 0.069868159 | Up | ENSMUSG00000107724.1 | Gm16042 |
| 1995.457842 | -0.823903223 | 0.26698078 | -3.086002007 | 0.002028675 | 0.069896308 | Down | ENSMUSG00000003438.17 | Timm50 |
| 2029.380239 | -0.878615808 | 0.284700129 | -3.08610962 | 0.00202794 | 0.069896308 | Down | ENSMUSG00000006517.5 | Mvd |
| 486.7440565 | -1.105057122 | 0.358223782 | -3.084823444 | 0.00203673 | 0.070031229 | Down | ENSMUSG00000033809.15 | Alg3 |
| 3731.5121 | -0.631949224 | 0.205147468 | -3.080463187 | 0.002066789 | 0.070920629 | Down | ENSMUSG00000109324.2 | Prmt1 |
| 1699.030684 | 0.686504129 | 0.222956684 | 3.079091943 | 0.002076326 | 0.071103655 | Up | ENSMUSG00000024712.9 | Rfk |
| 860.2223266 | 0.860120219 | 0.279434242 | 3.07807738 | 0.002083408 | 0.07120205 | Up | ENSMUSG00000026095.15 | Asnsd1 |
| 278.7585366 | 1.443837089 | 0.469240736 | 3.076964505 | 0.002091202 | 0.071324323 | Up | ENSMUSG00000025006.18 | Sorbs1 |
| 187.8281438 | 1.482460023 | 0.482057405 | 3.075276946 | 0.002103072 | 0.071584837 | Up | ENSMUSG00000062210.13 | Tnfaip8 |
| 981.1316098 | -0.897116883 | 0.291796112 | -3.074464823 | 0.002108806 | 0.071635882 | Down | ENSMUSG00000073436.11 | Eme2 |
| 786.0074781 | 0.83230014 | 0.270854803 | 3.07286461 | 0.002120147 | 0.071838244 | Up | ENSMUSG00000022772.12 | Senp5 |
| 435.434066 | 1.087524185 | 0.353962553 | 3.072427224 | 0.002123256 | 0.071838244 | Up | ENSMUSG00000048234.13 | Rnf149 |
| 10316.84692 | -0.674708855 | 0.219768392 | -3.070090515 | 0.002139939 | 0.072114241 | Down | ENSMUSG00000027546.15 | Atp9a |
| 261.6002309 | 1.20575619 | 0.392731968 | 3.070175814 | 0.002139328 | 0.072114241 | Up | ENSMUSG00000031578.5 | Mak16 |
| 89.25373258 | 3.685253055 | 1.200919525 | 3.068692762 | 0.002149976 | 0.072308429 | Up | ENSMUSG00000031722.10 | Hp |
| 1530.909138 | -0.715429995 | 0.233274499 | -3.066901854 | 0.002162899 | 0.072573332 | Down | ENSMUSG00000041528.15 | Rnf123 |
| 1957.214747 | -0.683582734 | 0.22292579 | -3.066413868 | 0.002166432 | 0.072573332 | Down | ENSMUSG00000049960.4 | Mrps16 |
| 2037.164024 | -0.916306973 | 0.299024255 | -3.064323239 | 0.002181631 | 0.072938031 | Down | ENSMUSG00000037966.15 | Ninj1 |
| 1070.79725 | -0.84416821 | 0.275655284 | -3.06240533 | 0.002195659 | 0.073124677 | Down | ENSMUSG00000048481.15 | Mypop |
| 536.3269469 | 1.136157106 | 0.371004834 | 3.062378177 | 0.002195858 | 0.073124677 | Up | ENSMUSG00000074749.10 | Kiz |
| 265.5667503 | 1.551911953 | 0.506899702 | 3.061575982 | 0.002201751 | 0.073176861 | Up | ENSMUSG00000034422.14 | Parp14 |
| 1267.825684 | -0.839894466 | 0.274408025 | -3.060750372 | 0.002207831 | 0.073235047 | Down | ENSMUSG00000048807.2 | Slc35e4 |
| 333.6431104 | -1.108383309 | 0.362323216 | -3.059100988 | 0.002220023 | 0.07349536 | Down | ENSMUSG00000074006.3 | Omp |
| 11549.84517 | -0.858159228 | 0.280850352 | -3.055574695 | 0.002246296 | 0.073882411 | Down | ENSMUSG00000002379.7 | Ndufa11 |
| 1782.905168 | -0.85034075 | 0.278378094 | -3.054625232 | 0.002253419 | 0.073882411 | Down | ENSMUSG00000002763.16 | Pex6 |
| 896.6743317 | -0.918637459 | 0.300700395 | -3.054992521 | 0.002250661 | 0.073882411 | Down | ENSMUSG00000009633.3 | G0s2 |
| 417.9007685 | 0.946526039 | 0.309902199 | 3.054273388 | 0.002256064 | 0.073882411 | Up | ENSMUSG00000027433.5 | Xrn2 |
| 89.09598904 | 3.408749135 | 1.116149027 | 3.054026883 | 0.002257918 | 0.073882411 | Up | ENSMUSG00000030921.17 | Trim30a |
| 768.9010829 | -1.353589006 | 0.442835208 | -3.056642701 | 0.002238309 | 0.073882411 | Down | ENSMUSG00000068264.11 | Ap5s1 |
| 1571.055196 | -1.025808665 | 0.335994816 | -3.053049085 | 0.002265289 | 0.073980484 | Down | ENSMUSG00000116165.1 | Pdxp |
| 12509.13942 | -0.850252388 | 0.278601284 | -3.051860976 | 0.002274274 | 0.074130818 | Down | ENSMUSG00000039515.11 | Ptpa |
| 193.1210967 | -1.446131121 | 0.474290628 | -3.049040051 | 0.002295739 | 0.074658955 | Down | ENSMUSG00000039628.9 | Hs3st6 |
| 934.6867008 | -0.748455232 | 0.245509951 | -3.048573919 | 0.002299303 | 0.074658955 | Down | ENSMUSG00000061118.8 | Dnajc30 |
| 11972.75524 | -0.710979231 | 0.233280911 | -3.047738577 | 0.002305704 | 0.074702567 | Down | ENSMUSG00000009863.14 | Sdhb |
| 3086.549706 | -0.808385299 | 0.265333805 | -3.046672844 | 0.002313894 | 0.074702567 | Down | ENSMUSG00000015013.9 | Trappc2l |
| 12.82416291 | 6.621674982 | 2.173382548 | 3.046713975 | 0.002313577 | 0.074702567 | Up | ENSMUSG00000086753.1 | Gm15751 |
| 1875.680161 | -0.738372917 | 0.242620204 | -3.043328232 | 0.00233977 | 0.075010528 | Down | ENSMUSG00000018820.13 | Zfyve27 |
| 6683.293284 | 0.704721855 | 0.231580583 | 3.043095606 | 0.002341579 | 0.075010528 | Up | ENSMUSG00000027199.14 | Gatm |
| 1430.369699 | 0.66828979 | 0.219508572 | 3.04448152 | 0.002330817 | 0.075010528 | Up | ENSMUSG00000036323.14 | Srp72 |
| 130.8387455 | 1.672406351 | 0.549667374 | 3.042578895 | 0.002345603 | 0.075010528 | Up | ENSMUSG00000040613.14 | Apobec1 |
| 797.0113824 | -0.866005227 | 0.284536451 | -3.043565152 | 0.002337928 | 0.075010528 | Down | ENSMUSG00000072941.5 | Sod3 |
| 351.2553379 | 1.089259587 | 0.358153784 | 3.041318101 | 0.002355448 | 0.075051168 | Up | ENSMUSG00000022718.12 | Dgcr8 |
| 7472.217038 | -0.679164458 | 0.223315337 | -3.041279947 | 0.002355747 | 0.075051168 | Down | ENSMUSG00000028798.16 | Eif3i |

| | | | | | | | | |
|---|---|---|---|---|---|---|---|---|
| 2078.427083 | -1.034126174 | 0.340187777 | -3.039868693 | 0.002366813 | 0.075261988 | Down | ENSMUSG00000033434.15 | Gtpbp6 |
| 189.938092 | 2.024364753 | 0.666295629 | 3.03823808 | 0.002379659 | 0.075528491 | Up | ENSMUSG00000015340.10 | Cybb |
| 13.1789033 | 6.660739776 | 2.194773417 | 3.034818867 | 0.002406802 | 0.076246942 | Up | ENSMUSG00000060131.11 | Atp8b4 |
| 1972.543233 | 0.654931639 | 0.21588016 | 3.033774104 | 0.002415152 | 0.07636846 | Up | ENSMUSG00000014195.16 | Dnajc7 |
| 1876.427621 | -0.848568227 | 0.279893429 | -3.031754733 | 0.002431367 | 0.076630857 | Down | ENSMUSG00000024856.10 | Cdk2ap2 |
| 741.8358063 | 1.670147021 | 0.550910395 | 3.03161283 | 0.00243251 | 0.076630857 | Up | ENSMUSG00000035929.11 | H2-Q4 |
| 1664.100198 | -1.273007862 | 0.420277595 | -3.02896913 | 0.002453898 | 0.077017783 | Down | ENSMUSG00000015337.5 | Endog |
| 8703.580549 | -0.628319294 | 0.20741204 | -3.029328927 | 0.002450977 | 0.077017783 | Down | ENSMUSG00000022048.8 | Dpysl2 |
| 527.4113606 | -1.189541143 | 0.3928103 | -3.02828399 | 0.002459468 | 0.077049679 | Down | ENSMUSG00000042293.8 | Gm5617 |
| 325.3754626 | -1.539548671 | 0.509236405 | -3.023249428 | 0.00250076 | 0.077340699 | Down | ENSMUSG00000000154.16 | Slc22a18 |
| 14296.8681 | -0.608447152 | 0.201118796 | -3.02531223 | 0.002483766 | 0.077340699 | Down | ENSMUSG00000019179.10 | Mdh2 |
| 671.5855321 | 1.136186177 | 0.37551832 | 3.025647795 | 0.002481011 | 0.077340699 | Up | ENSMUSG00000024908.15 | Ppp6r3 |
| 181.7924447 | 2.041540987 | 0.675089367 | 3.02410479 | 0.0024937 | 0.077340699 | Up | ENSMUSG00000026395.16 | Ptprc |
| 2416.61405 | -0.721033994 | 0.238331716 | -3.025337989 | 0.002483554 | 0.077340699 | Down | ENSMUSG00000041115.16 | Iqsec2 |
| 687.4292417 | -1.019728925 | 0.337141846 | -3.024628761 | 0.002489385 | 0.077340699 | Down | ENSMUSG00000041506.15 | Rrp9 |
| 2590.188478 | -0.711571315 | 0.235332186 | -3.023688879 | 0.002497131 | 0.077340699 | Down | ENSMUSG00000057375.13 | Yipf1 |
| 1591.546167 | -1.020025471 | 0.337564999 | -3.021715738 | 0.002513465 | 0.077591753 | Down | ENSMUSG00000003444.8 | Med29 |
| 2432.055808 | 0.605304882 | 0.200400856 | 3.020470539 | 0.002523823 | 0.07769594 | Up | ENSMUSG00000015536.14 | Mocs2 |
| 698.6565014 | -1.060507044 | 0.351367758 | -3.018225263 | 0.002542598 | 0.077784919 | Down | ENSMUSG00000022947.8 | Cbr3 |
| 3489.173344 | -0.64899981 | 0.215027929 | -3.018211701 | 0.002542712 | 0.077784919 | Down | ENSMUSG00000031781.14 | Ciapin1 |
| 246.1256847 | 1.326366268 | 0.439401919 | 3.018571862 | 0.002539692 | 0.077784919 | Up | ENSMUSG00000036572.16 | Upf3b |
| 5559.27884 | -0.643691743 | 0.21317779 | -3.01950659 | 0.002531868 | 0.077784919 | Down | ENSMUSG00000052456.9 | Get3 |
| 28656.85376 | -0.747986734 | 0.248045727 | -3.01551953 | 0.002565394 | 0.078195988 | Down | ENSMUSG00000041697.8 | Cox6a1 |
| 1482.986583 | -0.795674406 | 0.263835817 | -3.015793745 | 0.002563075 | 0.078195988 | Down | ENSMUSG00000048644.8 | Ctxn1 |
| 14.5652288 | 6.805007958 | 2.258984635 | 3.012418878 | 0.002591747 | 0.078857165 | Up | ENSMUSG00000106438.1 | Gm32051 |
| 2933.62321 | -0.878328283 | 0.291687441 | -3.011196779 | 0.002602202 | 0.079033113 | Down | ENSMUSG00000030083.11 | Abtb1 |
| 2298.064238 | -0.814111527 | 0.270608427 | -3.008448538 | 0.002625853 | 0.079182799 | Down | ENSMUSG00000004788.11 | Eif2b2 |
| 10803.48267 | -0.815404828 | 0.270994087 | -3.008939554 | 0.002621613 | 0.079182799 | Down | ENSMUSG00000031760.9 | Mt3 |
| 1201.586797 | -1.10489918 | 0.367214847 | -3.008863036 | 0.002622273 | 0.079182799 | Down | ENSMUSG00000042462.4 | Dctpp1 |
| 122.6872847 | 2.369514492 | 0.787224354 | 3.00996086 | 0.002612814 | 0.079182799 | Up | ENSMUSG00000078771.10 | Evi2a |
| 5038.559979 | -0.757838494 | 0.252086528 | -3.006263369 | 0.002644798 | 0.079587752 | Down | ENSMUSG00000030682.4 | Cdipt |
| 7674.405727 | -0.873375751 | 0.290561933 | -3.005816149 | 0.002648691 | 0.079587752 | Down | ENSMUSG00000033307.7 | Mif |
| 9273.445176 | -0.824989882 | 0.275002595 | -2.999934901 | 0.002700373 | 0.080568276 | Down | ENSMUSG00000023495.13 | Pcbp4 |
| 4678.209551 | -0.741064177 | 0.24701625 | -3.00006246 | 0.002699242 | 0.080568276 | Down | ENSMUSG00000025156.17 | Gps1 |
| 1463.615194 | -0.858193048 | 0.285918149 | -3.001534009 | 0.00268623 | 0.080568276 | Down | ENSMUSG00000045752.13 | Tssc4 |
| 12.37040245 | 6.569692024 | 2.189840912 | 3.000077306 | 0.002699111 | 0.080568276 | Up | ENSMUSG00000090031.2 | 4732440D04Rik |
| 7717.834916 | -0.73429305 | 0.244892021 | -2.998435992 | 0.002713692 | 0.080823098 | Down | ENSMUSG00000040479.12 | Dgkz |
| 5445.666667 | 0.621139891 | 0.207301655 | 2.996309366 | 0.00273269 | 0.081245912 | Up | ENSMUSG00000028639.14 | Ybx1 |
| 277.6928147 | 1.209256108 | 0.403776135 | 2.99486771 | 0.002745639 | 0.081487673 | Up | ENSMUSG00000024006.17 | Stk38 |
| 1413.424517 | 0.84740742 | 0.283126232 | 2.993037468 | 0.002762158 | 0.081834384 | Up | ENSMUSG00000024231.15 | Cul2 |
| 4126.613795 | -0.627109717 | 0.209673992 | -2.990879853 | 0.002781749 | 0.082270718 | Down | ENSMUSG00000023353.14 | Agap3 |
| 486.257566 | 1.568728278 | 0.524784533 | 2.989280705 | 0.002796351 | 0.082414407 | Up | ENSMUSG00000033355.6 | Rtp4 |
| 11584.9045 | -0.748236613 | 0.250271175 | -2.989703522 | 0.002792483 | 0.082414407 | Down | ENSMUSG00000037032.16 | Apbb1 |
| 2555.144241 | -0.664506423 | 0.222508387 | -2.986433155 | 0.002822525 | 0.082945537 | Down | ENSMUSG00000020225.10 | Tmbim4 |
| 3613.085125 | -0.805091466 | 0.269599115 | -2.986254109 | 0.002824179 | 0.082945537 | Down | ENSMUSG00000060376.7 | Bckdha |
| 1500.670268 | 0.95004854 | 0.318427206 | 2.983565862 | 0.002849107 | 0.08338814 | Up | ENSMUSG00000028906.17 | Epb41 |
| 1623.845182 | 0.752348486 | 0.25215873 | 2.98363053 | 0.002848505 | 0.08338814 | Up | ENSMUSG00000038668.14 | Lpar1 |
| 376.9208615 | -1.025827597 | 0.344443184 | -2.978220047 | 0.002899278 | 0.084709979 | Down | ENSMUSG00000039648.14 | Kyat1 |
| 916.1593665 | 0.759518527 | 0.255113933 | 2.977173841 | 0.00290919 | 0.084765612 | Up | ENSMUSG00000001036.17 | Epn2 |
| 142.100724 | 1.992827349 | 0.669416504 | 2.976961783 | 0.002911203 | 0.084765612 | Up | ENSMUSG00000020380.16 | Rad50 |
| 7764.451259 | -0.736105677 | 0.247317955 | -2.976353565 | 0.002916984 | 0.08478799 | Down | ENSMUSG00000024870.6 | Rab1b |
| 1866.264763 | -0.710968557 | 0.238987205 | -2.974923105 | 0.00293062 | 0.084892634 | Down | ENSMUSG00000011589.9 | Fsd1 |
| 2801.020189 | -0.965530664 | 0.32454982 | -2.974984434 | 0.002930034 | 0.084892634 | Down | ENSMUSG00000078570.10 | 1110065P20Rik |
| 2735.00843 | -0.85101314 | 0.286350503 | -2.971928215 | 0.002959359 | 0.085373874 | Down | ENSMUSG00000023939.7 | Mrpl14 |
| 182.5906957 | 1.881384037 | 0.633118234 | 2.971615628 | 0.002962373 | 0.085373874 | Up | ENSMUSG00000024672.11 | Ms4a7 |
| 971.9538278 | -0.806071788 | 0.271193193 | -2.972315716 | 0.002955626 | 0.085373874 | Down | ENSMUSG00000032609.12 | Klhdc8b |
| 9545.94538 | -0.827111799 | 0.278484474 | -2.970046367 | 0.002977548 | 0.085665272 | Down | ENSMUSG00000002416.13 | Ndufb2 |
| 332.6915348 | 1.090341865 | 0.36725021 | 2.968934626 | 0.002988342 | 0.08582984 | Up | ENSMUSG00000027883.15 | Gpsm2 |
| 6495.762971 | -0.641879419 | 0.216294285 | -2.967620797 | 0.003001143 | 0.086051426 | Down | ENSMUSG00000065990.12 | Aurkaip1 |
| 1041.869137 | -0.75611327 | 0.254846632 | -2.966934524 | 0.00300785 | 0.086097799 | Down | ENSMUSG00000031970.16 | Dbndd1 |
| 2141.524593 | -0.828576032 | 0.279478337 | -2.964723637 | 0.00302955 | 0.086572449 | Down | ENSMUSG00000078681.10 | Tm2d3 |
| 11066.64346 | -0.745834816 | 0.251698949 | -2.963201945 | 0.003044568 | 0.086854889 | Down | ENSMUSG00000055839.6 | Elob |
| 2444.586684 | 0.628731481 | 0.21221856 | 2.962660204 | 0.003049931 | 0.086861405 | Up | ENSMUSG00000021998.16 | Lcp1 |
| 1527.835912 | -0.63325855 | 0.213809248 | -2.961792135 | 0.003058542 | 0.086960258 | Down | ENSMUSG00000004961.7 | Syt5 |
| 5384.458106 | -0.757525562 | 0.255895913 | -2.960287852 | 0.003073517 | 0.08713631 | Down | ENSMUSG00000001313.12 | Rnd2 |
| 732.89696 | 0.79922733 | 0.270119846 | 2.958787891 | 0.003088516 | 0.08713631 | Up | ENSMUSG00000021681.8 | Aggf1 |
| 3184.561083 | -0.994631052 | 0.336024663 | -2.959994197 | 0.003076448 | 0.08713631 | Down | ENSMUSG00000031807.10 | Pgls |
| 775.3912205 | 0.800500939 | 0.270497672 | 2.959363499 | 0.003082752 | 0.08713631 | Up | ENSMUSG00000038371.15 | Sbf2 |
| 597.4136007 | 0.922138454 | 0.311681607 | 2.958591176 | 0.003090488 | 0.08713631 | Up | ENSMUSG00000042363.14 | Lgalsl |
| 1087.746052 | -0.647878062 | 0.219150744 | -2.956312395 | 0.003113416 | 0.087636695 | Down | ENSMUSG00000004846.10 | Plod3 |
| 9856.762824 | -0.826021074 | 0.279585306 | -2.954450955 | 0.003132259 | 0.08802065 | Down | ENSMUSG00000032171.7 | Pin1 |
| 7325.5166 | -0.772481639 | 0.261512359 | -2.953901082 | 0.003137846 | 0.088031401 | Down | ENSMUSG00000030707.15 | Coro1a |
| 1582.676238 | -0.58598228 | 0.198422408 | -2.95320617 | 0.003144918 | 0.088083751 | Down | ENSMUSG00000040964.16 | Arhgef10l |
| 2881.507837 | -0.848164929 | 0.287562785 | -2.949494762 | 0.00318294 | 0.089001307 | Down | ENSMUSG00000078348.4 | Sf3b5 |
| 1806.96096 | -0.775276286 | 0.263096438 | -2.946738055 | 0.003211451 | 0.089636957 | Down | ENSMUSG00000031458.8 | Coprs |
| 1996.395597 | -0.784837966 | 0.266383189 | -2.94627438 | 0.00321627 | 0.089636957 | Down | ENSMUSG00000036372.14 | Tmem258 |

| | | | | | | | | |
|---|---|---|---|---|---|---|---|---|
| 1651.509973 | -0.90791598 | 0.308221978 | -2.945656198 | 0.003222704 | 0.089668555 | Down | ENSMUSG00000071078.6 | Nr2c2ap |
| 4323.371241 | -0.722267533 | 0.245307681 | -2.944333137 | 0.003236514 | 0.089887195 | Down | ENSMUSG00000032112.10 | Trappc4 |
| 10420.10794 | -0.763936595 | 0.25949934 | -2.943886471 | 0.003241189 | 0.089887195 | Down | ENSMUSG00000073616.10 | Cops9 |
| 12.35127006 | 6.567193669 | 2.232187099 | 2.942044451 | 0.003260531 | 0.090275624 | Up | ENSMUSG00000034438.16 | Gbp8 |
| 1339.106164 | -0.759069061 | 0.258245342 | -2.939333021 | 0.003289195 | 0.090916465 | Down | ENSMUSG00000026799.15 | Med27 |
| 521.5920334 | 0.889132775 | 0.302545431 | 2.938840533 | 0.003294425 | 0.090916465 | Up | ENSMUSG00000045969.8 | Ing1 |
| 7420.266265 | -0.717416695 | 0.244268344 | -2.937002327 | 0.003314016 | 0.091174148 | Down | ENSMUSG00000039640.7 | Mrpl12 |
| 1549.422629 | -0.665365819 | 0.226549687 | -2.936953157 | 0.003314542 | 0.091174148 | Down | ENSMUSG00000041241.12 | Mul1 |
| 5232.092102 | -0.841987745 | 0.287006968 | -2.933683985 | 0.003349651 | 0.091910095 | Down | ENSMUSG00000035964.8 | Tmem59l |
| 2519.758983 | -0.868629938 | 0.296111933 | -2.933451314 | 0.003352162 | 0.091910095 | Down | ENSMUSG00000037960.11 | Card19 |
| 21247.01266 | -0.742024457 | 0.253013962 | -2.932741145 | 0.003359839 | 0.091971512 | Down | ENSMUSG00000028931.12 | Kcnab2 |
| 1335.369384 | -0.918578565 | 0.313275237 | -2.932177385 | 0.003365944 | 0.09198979 | Down | ENSMUSG00000020778.12 | Ten1 |
| 577.8798136 | 1.21204666 | 0.413694294 | 2.929812372 | 0.003391667 | 0.092543283 | Up | ENSMUSG00000030220.13 | Arhgdib |
| 5005.678292 | -0.780357851 | 0.266815337 | -2.924711377 | 0.003447758 | 0.093448067 | Down | ENSMUSG00000005881.13 | Ergic3 |
| 11.89602126 | 6.513457235 | 2.226982725 | 2.924790193 | 0.003446885 | 0.093448067 | Up | ENSMUSG00000021356.10 | Irf4 |
| 2519.212358 | 0.790334341 | 0.270248075 | 2.924477229 | 0.003450353 | 0.093448067 | Up | ENSMUSG00000026234.12 | Ncl |
| 13.20054886 | 6.663003712 | 2.277202782 | 2.925959762 | 0.003433953 | 0.093448067 | Up | ENSMUSG00000030142.10 | Clec4e |
| 19422.7075 | -0.549383045 | 0.187892287 | -2.923925472 | 0.003456475 | 0.093448067 | Down | ENSMUSG00000031144.15 | Syp |
| 15824.2443 | -0.751963206 | 0.25718776 | -2.923790794 | 0.003457971 | 0.093448067 | Down | ENSMUSG00000044894.14 | Uqcrq |
| 551.9532827 | -1.219637121 | 0.417235082 | -2.923141349 | 0.003465192 | 0.093493857 | Down | ENSMUSG00000063897.3 | |
| 1797.721012 | 0.776297648 | 0.265771461 | 2.920921779 | 0.003489974 | 0.093849509 | Up | ENSMUSG00000003031.14 | Cdkn1b |
| 4849.657485 | -0.496477675 | 0.169999123 | -2.920471974 | 0.003495016 | 0.093849509 | Down | ENSMUSG00000020265.16 | Sumo3 |
| 809.614643 | -0.953327193 | 0.326340705 | -2.92126351 | 0.003486148 | 0.093849509 | Down | ENSMUSG00000025175.12 | Fn3k |
| 61.04504615 | -2.652525681 | 0.908712804 | -2.918992304 | 0.003511649 | 0.093997729 | Down | ENSMUSG00000035274.13 | Tpbg |
| 1719.525399 | -0.951101845 | 0.325807933 | -2.91921021 | 0.003509195 | 0.093997729 | Down | ENSMUSG00000059895.13 | Ptp4a3 |
| 3303.724598 | -0.877963438 | 0.300842605 | -2.918348078 | 0.003518913 | 0.094043367 | Down | ENSMUSG00000070570.5 | Slc17a7 |
| 38.0760446 | 4.550141997 | 1.560801645 | 2.915259611 | 0.003553928 | 0.094829331 | Up | ENSMUSG00000053835.17 | H2-T24 |
| 23464.6161 | -0.767649399 | 0.263504531 | -2.913230353 | 0.003577106 | 0.095135866 | Down | ENSMUSG00000024953.17 | Prdx5 |
| 4706.71854 | -0.700658033 | 0.240466869 | -2.913740414 | 0.003571267 | 0.095135866 | Down | ENSMUSG00000040722.7 | Scamp5 |
| 3351.360762 | -0.78196134 | 0.268458916 | -2.912778429 | 0.003582287 | 0.095135866 | Down | ENSMUSG00000061451.13 | Tmem151a |
| 5804.032286 | -0.804293774 | 0.276247682 | -2.911495106 | 0.003597035 | 0.095377818 | Down | ENSMUSG00000029544.16 | Cabp1 |
| 4990.850386 | -0.754563826 | 0.259305178 | -2.909945073 | 0.003614923 | 0.095660817 | Down | ENSMUSG00000033287.16 | Kctd17 |
| 701.7981284 | -0.738383339 | 0.253775629 | -2.909591209 | 0.003619018 | 0.095660817 | Down | ENSMUSG00000038524.13 | Fchsd1 |
| 68018.90815 | -0.966041586 | 0.33221239 | -2.907903541 | 0.003638606 | 0.096028533 | Down | ENSMUSG00000024608.11 | Rps14 |
| 112854.5852 | -0.515992122 | 0.177673095 | -2.904165777 | 0.003682331 | 0.096575468 | Down | ENSMUSG00000019505.7 | Ubb |
| 158.4376233 | 1.436823991 | 0.494698323 | 2.904444837 | 0.00367905 | 0.096575468 | Up | ENSMUSG00000025395.14 | Prim1 |
| 598.2193809 | -1.017073882 | 0.350163985 | -2.904564506 | 0.003677644 | 0.096575468 | Down | ENSMUSG00000029875.5 | Ccdc184 |
| 3572.477616 | -0.649537125 | 0.223688622 | -2.903755755 | 0.003687157 | 0.096575468 | Down | ENSMUSG00000074247.10 | Dda1 |
| 13.39376363 | 6.683788299 | 2.301821811 | 2.90369492 | 0.003687873 | 0.096575468 | Up | ENSMUSG00000099757.1 | BE692007 |
| 1970.817912 | 0.666277394 | 0.229606601 | 2.901821601 | 0.003709998 | 0.09700468 | Up | ENSMUSG00000024588.9 | Fech |
| 2682.120484 | -0.860355776 | 0.296631463 | -2.900419821 | 0.003726632 | 0.09728924 | Down | ENSMUSG00000040759.9 | Cmtm5 |
| 368.3092567 | -1.279298409 | 0.441588824 | -2.897035295 | 0.003767074 | 0.097896526 | Down | ENSMUSG00000020877.11 | Scrn2 |
| 15313.79581 | -0.524553677 | 0.18113807 | -2.895877592 | 0.003780999 | 0.097896526 | Down | ENSMUSG00000026223.15 | Itm2c |
| 3776.737304 | -0.662672018 | 0.228810023 | -2.896166911 | 0.003777514 | 0.097896526 | Down | ENSMUSG00000026965.12 | Anapc2 |
| 1345.671205 | -0.753577105 | 0.26025105 | -2.895577574 | 0.003784615 | 0.097896526 | Down | ENSMUSG00000052926.16 | Rnaseh2a |
| 447.0244813 | -1.105185857 | 0.381500909 | -2.896941609 | 0.003768199 | 0.097896526 | Down | ENSMUSG00000070699.5 | Sars2 |
| 945.1434262 | 1.046446714 | 0.361112869 | 2.897838336 | 0.003757442 | 0.097896526 | Up | ENSMUSG00000072946.12 | Ptgr2 |
| 14197.58946 | -0.678473744 | 0.234708983 | -2.890702078 | 0.003843823 | 0.099171714 | Down | ENSMUSG00000020496.10 | Rnf187 |
| 391.6945394 | 1.126192631 | 0.389611374 | 2.890553786 | 0.003845637 | 0.099171714 | Up | ENSMUSG00000042742.7 | Bmt2 |
| 19.09611835 | 7.195848135 | 2.49074977 | 2.889028927 | 0.003864335 | 0.099502221 | Up | ENSMUSG00000069892.9 | 9930111J21Rik2 |
| 82365.57302 | -0.657551413 | 0.22767993 | -2.888051715 | 0.003876361 | 0.09966019 | Down | ENSMUSG00000057666.18 | Gapdh |
